# Early endothelial activation at the blood–nerve barrier defines a hallmark of ALS

**DOI:** 10.1101/2025.08.31.673224

**Authors:** Ganesh Parameshwar Bhat, Santo Diprima, Cristian G Beccaria, Paola Podini, Aurora Badaloni, Alessia Motta, Ilaria Brambilla, Luis Martins, Daniela Boselli, Stefano de Pretis, Giovanni Nardo, Caterina Bendotti, Nilo Riva, Matteo Iannacone, Angelo Quattrini, Dario Bonanomi

## Abstract

Vascular defects are common in Amyotrophic Lateral Sclerosis (ALS). The prevailing view is that breakdown of the blood–brain and blood–spinal cord barriers contributes to neurodegeneration. Here, we reveal that selective and early vulnerability of peripheral nerve endothelium —manifested as endothelial cell activation and interlinked blood–nerve barrier dysfunction— constitutes a core feature of ALS pathogenesis, arising before vascular alterations in the central nervous system (CNS) and motor neuron pathology.

Vascular changes in ALS patients have been largely studied in postmortem samples, limiting insight into their onset, causes, and pathogenic role. Surveying diagnostic motor nerve samples from ALS patients across clinical stages, we observed endothelial damage that preceded axonal loss and demyelination, marking vascular dysfunction as an early disease event. Similar ultrastructural abnormalities were detected in pre-symptomatic ALS mouse models (*SOD1* and *TARDBP* mutants). Notably, endothelial cells became dysfunctional even when not carrying mutant TDP-43, indicating they respond to non-cell autonomous disease signals. Transcriptional, histological, and functional analyses revealed that these alterations were largely confined to peripheral nerves, while spinal cord vessels exhibited delayed and more focal changes. Single-cell sequencing identified ALS-susceptible endothelial cell subsets prone to a pro-inflammatory phenotype and impaired blood–nerve barrier function, increasing permeability via the transcellular route. These changes coincided with reactivity of nerve-resident macrophages and neutrophil infiltration. Neutrophil depletion attenuated endothelial activation and barrier leakiness, mitigating axonopathy in ALS mice.

Our work unmasks the greater susceptibility of the peripheral nerve vasculature in ALS relative to the CNS. The early activation of peripheral nerve endothelium, combined with its potential reversibility, identifies a therapeutic window and suggests strategies for targeting the vascular–immune axis to protect the motor system.

## INTRODUCTION

The architecture, composition, and molecular properties of vascular beds reflect the structural and functional requirements of the surrounding tissue^1^. Endothelial cells (ECs) that line these vessels embody this diversity and specialization^2,3^. Far from being an inert interface, the endothelium is metabolically active and plastic. Beyond their circulatory role in delivering oxygen and nutrients, ECs —together with associated cells in tissue-specific vascular niches— maintain organ homeostasis, contributing to vascular regulation, immune surveillance, and tissue repair^4–6^. At the same time, ECs mount maladaptive responses to pathological stimuli that exacerbate disease^7^. This specialization and adaptive capacity also shape how ECs respond to pathophysiological stress^7^. These observations raise the question of whether the mechanisms that confer EC specialization and plasticity may also render them selectively susceptible to damage, generating focal patterns of vulnerability.

A prototypical example of highly specialized endothelium is found in the vascular barriers of the CNS — the blood-brain barrier (BBB) and blood-spinal cord barrier (BSCB). CNS ECs acquire a barrier phenotype characterized by tight junctions, low rates of transcellular transport, restricted leukocyte adhesion molecule expression, and specialized transporter systems, all of which are essential for maintaining CNS homeostasis^8–10^. These barriers are disrupted in neurological disorders, including ALS, where BBB and BSCB defects are associated with neuroinflammation and motor neuron dysfunction^11–13^. In ALS patients and mouse models, BSCB abnormalities include reduced tight junction protein expression, decreased pericyte and astrocytic coverage, and activation of perivascular fibroblasts^14–23^. Increased levels of serum proteins have been detected in the cerebrospinal fluid of ALS patients, indicating BBB/BSCB dysfunction^24–30^, although these changes do not clearly align with disease aggressiveness^31^ or the spatial pattern of motor neuron loss^32^. In ALS rodent models, BSCB impairment has been observed prior to motor neuron degeneration^19,20,22,33–35^, whereas other studies reported an intact barrier to systemically administered IgG^36^. While the relationship between CNS barrier leakage and neurodegeneration remains to be resolved, the expression of pathogenic gene variants in ECs is sufficient to trigger vascular dysfunction —including BBB/BSCB alterations^37–39^— and motor neuron degeneration^40^ via non-cell autonomous disease mechanisms^41^.

Although ALS research has primarily focused on CNS vascular damage and barrier dysfunction, spinal motor neurons —the main disease targets— extend most of their volume into the peripheral nervous system (PNS), where motor axons travel long distances within peripheral nerves to innervate skeletal muscles and autonomic ganglia^42^. This unique positioning at the CNS–PNS interface, together with the considerable heterogeneity in ALS onset and progression^43,44^, raises questions about where the disease initiates and how it propagates^45,46^. Early pathogenic events, including direct axonal damage and a “dying-back” phenomenon from the PNS, have been proposed^47–49^ and likely act in concert with “dying-forward” mechanisms of disease spreading from the CNS^45^. Supporting PNS involvement, transcriptomic analyses and histopathological assessment of diagnostic motor nerve biopsies from ALS patients have revealed early alterations, including accumulation of pathological phopho-TDP-43 aggregates in axon fibers and Schwann cells —the PNS glia— before neurodegeneration^50,51^. Schwann cells can trigger neuroinflammation via chemokine secretion^52^, while both nerve-resident macrophages and infiltrating immune cells appear to play context-dependent roles that may be either protective or deleterious ^53–57^. Collectively, these observations underscore the critical role of the peripheral nerve microenvironment in ALS pathogenesis.

In the PNS, motor axons run alongside capillaries within the endoneurial compartment —the innermost layer of the nerve— whose ECs form the blood–nerve barrier (BNB), analogous to CNS barriers but comparatively less well studied^58,59^. The BNB shares key features with CNS barriers, including endothelial tight junctions, low rates of transcellular transport, and pericyte coverage^60^, but, unlike CNS barriers, it lacks astrocytic endfeet —critical components of neurovascular units^61^. The baseline permeability of the BNB is higher than that of the BBB^62,63^, and its ECs exhibit enhanced angiogenic capacity, supporting regeneration after injury^64,65^ —a distinctive feature of PNS nerves that is absent in the mammalian CNS. The importance of the BNB for normal nerve physiology is underscored by the fact that its alteration is associated with several peripheral neuropathies, including diabetic and inflammatory neuropathies^58,59^. Although BNB changes in ALS remain poorly explored, studies in *SOD1* mutant rat models have shown endoneurial edema and macrophage activation in motor nerves, indicating early BNB impairment that precedes axonal degeneration^66^.

We show that peripheral nerves harbor a vulnerable vasculature in ALS. Endothelial activation —defined by an inflammatory and procoagulant phenotype^7^— coincided with BNB disruption, driven primarily by aberrant transcellular permeability in motor nerves before CNS vascular pathology and neurodegeneration. Venous capillaries were intrinsically more susceptible, displaying an elevated inflammatory tone that distinguished them from CNS counterparts. ECs became dysfunctional even in the absence of ALS-linked mutations, indicating that they responded to non-cell-autonomous disease signals. These vascular changes were accompanied by remodeling of nerve-resident macrophages and neutrophil infiltration, which further compromised barrier integrity. Collectively, these findings identify the peripheral nerve endothelium as an active sensor and amplifier of ALS pathology, revealing a previously underappreciated site of vulnerability in motor neuron disease.

## RESULTS

### Early microvascular changes in peripheral nerves of ALS patients and mouse models

Retrospective transmission electron microscopy (TEM) analysis of motor nerve (*obturator n.*) biopsies collected to assist in diagnosis of motor neuron disease revealed a high incidence of microvascular abnormalities in clinically confirmed cases of sporadic ALS (n=12) (Fig. 1a, i). Nerve samples were categorized from “normal” to “severe” based on the extent of axonal loss, assessed in semithin sections (Fig. 1d-h). Biopsies from non-ALS neuropathy or myopathy patients, without evidence of demyelination or axonal pathology, served as controls (n=3). In ALS nerves, ECs lining endoneurial blood vessels (Fig. 1c) exhibited abnormal accumulation of pinocytotic vesicles, tight junctions (TJ) alterations, marked hypertrophy leading to luminal narrowing, central microfilament bundles resembling atherosclerotic lesions^67^, detachment from the basal lamina, and hyperplasia (Fig. 1d’-h’, i and Suppl. Fig. 1a-l). While the overall severity of endothelial damage correlated with axonal pathology, the aberrant increase in cytoplasmic vesicles —indicative of enhanced transcytosis (transcellular transport)— was already evident in histologically “normal” ALS cases, prior to significant axonal loss (Fig. 1e, e’, f, f’, i, j). Conversely, TJ opening —expected to increase paracellular permeability— along with EC hyperplasia and basal lamina detachment, were observed in cases with more extensive axonal loss (Fig. 1h, h’, i and Suppl. Fig. 1j-k).

**Fig. 1:**
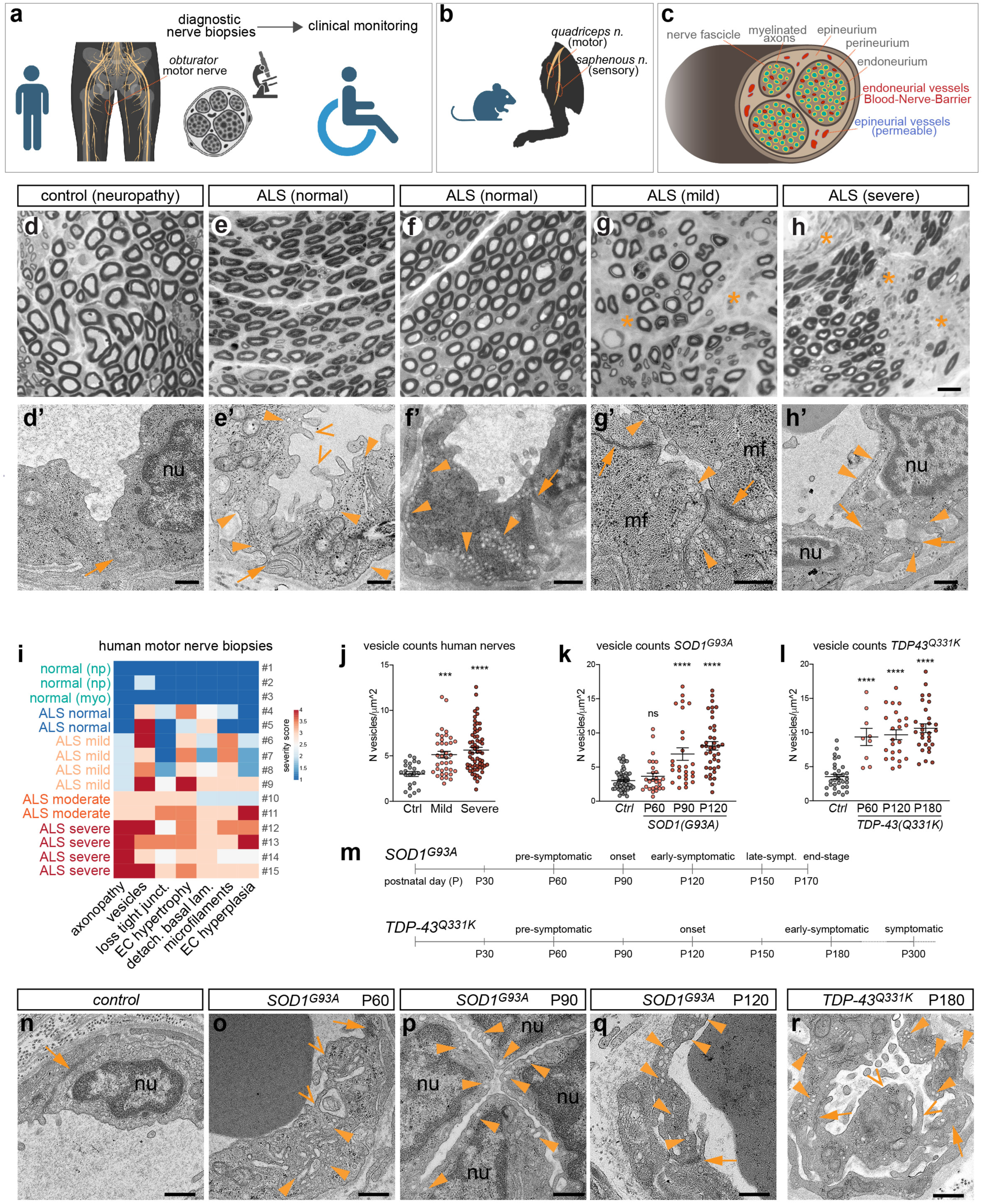
Microvascular alterations in peripheral nerves of ALS patients and mouse models. **a**, Diagnostic workflow for ALS based on histopathological analysis of motor nerve (*obturator n*.) biopsies. **b**, Schematic of *quadriceps* motor nerve and *saphenous* sensory nerve in mouse hindlimb. **c**, Schematic of peripheral nerve showing blood-nerve-barrier localization in endoneurial microvessels and permeable vasculature of the epineurial compartment. **d-h**, Toluidine Blue-stained semithin transverse sections of diagnostic *obturator* nerve biopsies from non-ALS (control, **d**) and sporadic ALS cases (**e-h**) with varying degrees of fiber loss (asterisks), reflecting different stages of disease progression. **d’-h’**, Corresponding TEM images of endoneurial ECs. Pinocytotic vesicles (closed arrowheads) accumulate in ALS nerves. TJ (arrows) are compromised in severe cases (**h’**). EC hypertrophy, open arrowhead, **e’**. Microfilaments (mf; **g’**). Multiple EC nuclei (nu; **h’**) indicate hyperplasia. **i**, Heatmap of severity score for axonopathy (fiber loss, demyelination) and morphological alterations in endoneurial ECs from *obturator* nerve biopsies. Controls are non-ALS neuropathy (np) and myopathy (myo) cases with normal nerve fiber morphology. Case numbers are reported (see Suppl. Fig. 1). **j**, Vesicle density in human motor nerve ECs. ALS “normal” cases are included in the mild group; “moderate" ALS cases in the severe group. Mean ± SEM. **Sample size and statistics for all figures are reported in Methods.** **k**, **l**, Vesicle density in motor nerve ECs from *SOD1^G93A^* (**k**) and *TDP-43^Q331K^* (**l**) mice at progressive disease stages. Mean ± SEM **m**, Timeline of disease progression in *SOD1^G93A^* and *TDP-43^Q331K^* mouse models. **n-r**, TEM images of endoneurial ECs from control (**n**), *SOD1^G93A^* (**o**, **p**, **q**) and *TDP43^Q331K^* (**r**) motor nerves at the indicated stages. ECs in mutants show increased cytoplasmic vesicles (closed arrowheads, **o-r**), hypertrophy (open arrowheads, **o**, **r**), hyperplasia (nu, nuclei, **p**). TJs (arrows) are maintained through the early-symptomatic stage (**q**, **r**). Scale bars: d-h, 20μm; d’-h’, 500nm; n-r, 500nm.

A spectrum of EC alterations similar to those observed in human ALS nerve biopsies was identified by TEM in the microvasculature of motor nerves (*quadriceps n.*) of two transgenic mouse models of ALS, *SOD1^G93A^*and *TDP-43^Q331K^* (Fig. 1b, n-r and Suppl. Fig. 2i-n, u-w). These models develop progressive motor neuron disease with distinct temporal patterns, as defined by histological and behavioral assays (Fig. 1m, Suppl. Fig. 2g, h and Suppl. Fig. 3o-q)^68–70^. Axonal loss in *SOD1^G93A^*motor nerves began approximately at 90 days of age (postnatal, P90; onset stage) and worsened by P120 (early-symptomatic) (Suppl. Fig. 2a-f)^53^. In *TDP-43^Q331K^* mice, which moderately overexpress human mutant TDP-43 without mislocalization and potentially mirror an early stage of ALS^68,70^, only a few degenerating axons were observed by the onset of motor symptoms at P120 (Suppl. Fig. 2o-t). As noted in patient nerve biopsies, endoneurial ECs in both *SOD1^G93A^* and *TDP-43^Q331K^* motor nerves showed a conspicuous increase in cytoplasmic vesicle density by the onset stage (Fig. 1k, l, o-r and Suppl. Fig. 2m’, n, u’, v’), while TJs remained relatively preserved through the early-symptomatic phase (Fig. 1q, r and Suppl. Fig. 2m’, u’, v’). These defects were never detected in control motor nerves from non-transgenic littermates, nor in sensory nerves (*saphenous n.*) of mutant mice, supporting the regional specificity of ALS pathology in these models (Fig. 1b, n and data not shown)^69^.

These findings indicate that endothelial alterations in the peripheral nerve microvasculature arise at the pre-symptomatic stage, preceding axonal degeneration, and may represent a shared pathogenic feature of both familial and sporadic ALS, as observed across different transgenic mouse models and in patients with distinct clinical presentations.

### Heightened vulnerability of peripheral nerve endothelium in ALS

The structural changes identified in the endothelium of human and mouse peripheral nerves point to a compromised blood-nerve barrier (BNB) (Fig. 1c). To assess vascular barrier integrity, we intravenously administered the permeability tracer sulfo-NHS-biotin^64,71,72^ to *SOD1^G93A^* mice at different disease stages and examined its distribution in sciatic nerves, which carry both motor and sensory fibers (Fig. 2a-e). In control mice, sulfo-biotin was confined to the lumen of endoneurial vessels, confirming an intact BNB (Fig. 2b). In contrast, focal leakage into the nerve parenchyma was already evident in *SOD1^G93A^*mice at presymptomatic stage (P60) and worsened from onset (P90) onward (Fig. 2c-f and Suppl. Fig. 3a-b’). BNB dysfunction was limited to motor nerves, with no leakage observed in sensory nerves at any stage (Fig. 2h-j’), aligning with the pattern of ultrastructural EC alterations. Notably, in the spinal cord of *SOD1^G93A^*mice, sulfo-biotin was retained in the microvasculature until the symptomatic phase (P120-140) (Fig. 2g), when only sporadic leaky hotspots became visible despite widespread astrogliosis, microglial activation, and motor neuron loss (Fig. 2k, l’ and Suppl. Fig. 3c-q). In parallel, *SOD1^G93A^* sciatic nerves showed progressive capillary dilation and tortuosity, features associated with inflammation, hypoxia, and barrier permeability in other conditions^73,74^. By contrast, spinal cord capillary size and morphology remained largely unchanged, with only localized alterations observed at advanced stages (Suppl. Fig. 3r-x). Together, these findings demonstrate early BNB disruption, preceding blood-spinal cord barrier (BSCB) impairment^19,20,33^.

**Fig. 2:**
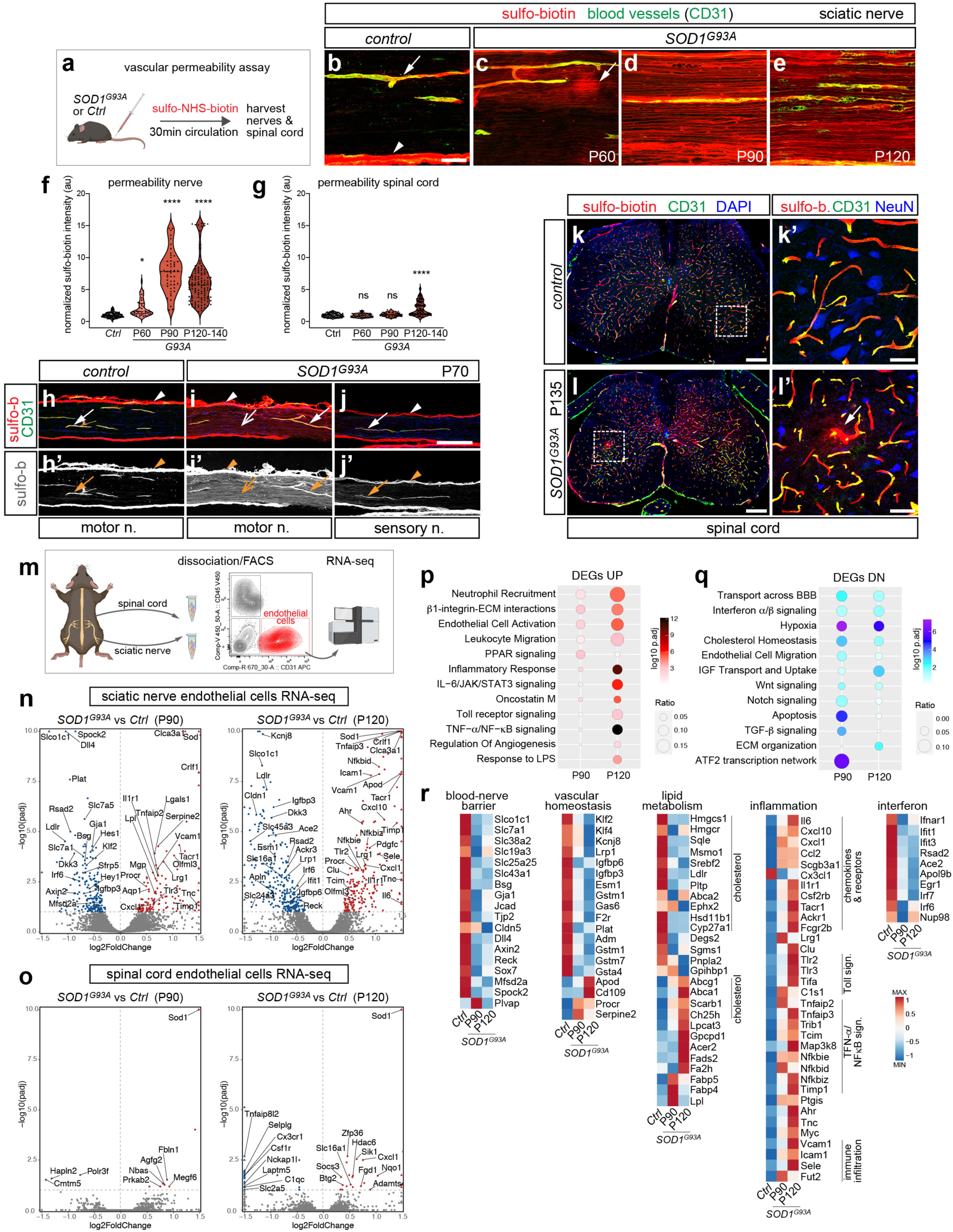
Blood-nerve barrier disruption and transcriptional dysregulation in the peripheral nerve endothelium of *SOD1^G93A^* ALS mice. **a**, Vascular permeability assessed through intravenous injection of sulfo-NHS-biotin. **b-e**, Longitudinal views of sciatic nerves from control (**b**) and *SOD1^G93A^* (**c-e**) mice injected with sulfo-biotin (550 Da). The tracer is retained in BNB-forming endoneurial vessels (CD31^+^) in controls (**b**, arrow) but extravasates in mutants (**c-e**). Leakage is focal at pre-symptomatic stage (P60; **c**, arrow) and widespread from onset (P90) onward (**d**, **e**). Sulfo-biotin accumulates in the epineurium that contains permeable vessels (**b**, arrowhead). **f**, **g** Extravascular sulfo-biotin signal in control and *SOD1^G93A^*sciatic nerves (**f**) and spinal cord (**g**) at progressive stages, normalized to average control value. **h-j’**, Vascular leakage in *quadriceps* motor nerve of mice injected with sulfo-biotin at P70 (**i**, **i’**; open arrows). The tracer is retained in endoneurial vessels (CD31^+^) of controls (**h**, **h’**, closed arrows) and *SOD1^G93A^ saphenous* sensory nerve (**j**, **j’**^+^, closed arrows). Sulfo-biotin is shown separately in **h’-j’**. Arrowheads point to epineurium. **k-l’**, Spinal cord transverse sections from control (**k**, **k’**) and *SOD1^G93A^* (**l**, **l’**) mice injected with sulfo-biotin at P135. Boxed areas in **k**, **l** are magnified in **k’**, **l’**. The tracer is confined to vessels (CD31^+^) in controls. Focal leakage is visible in *SOD1^G93A^* (**l’**, arrow). DAPI labels nuclei (**k**, **l**). NeuN labels neurons (**k’**, **l’**). **m**, Schematic of bulk RNA-seq of ECs FACS-purified separately from sciatic nerves and spinal cord using anti-CD31 antibody. **n**, **o**, Volcano plots of EC genes upregulated (red) or downregulated (blue) (padj <0.1) in *SOD1^G93A^* sciatic nerve (**n**) or spinal cord (**o**) vs. control at P90 and P120. Selected genes are labeled. **p**, **q**, GO enrichment from upregulated (**p**) and downregulated (**q**) genes (padj ≤0.05) in *SOD1^G93A^* sciatic nerve vs. control at P90 and P120. Spinal cord DEGs are too few for GO analysis. **r**, Average expression heatmaps (z-score) of representative genes for deregulated biological processes. Scale bars: b-e, 50μm; h-j’, 200μm; k, l, 200μm; k’, l’, 50μm.

We asked whether the enhanced susceptibility of the BNB in *SOD1^G93A^* mice, compared to the BSCB, was reflected in distinct transcriptional changes in the endothelium. To address this, we isolated ECs from the sciatic nerve and spinal cord of *SOD1^G93A^*mice and non-transgenic controls at P90 and P120 using fluorescence-activated cell sorting (FACS) with an anti-CD31 antibody, followed by bulk RNA-sequencing (Fig. 2m). The sciatic nerve epineurium was mechanically removed to enrich BNB-forming endoneurial ECs. After confirming EC population purity (Suppl. Fig. 3y), differential gene expression (DGE) analysis showed divergent transcriptional responses of sciatic nerve versus spinal cord ECs in *SOD1^G93A^* mice. In the sciatic nerve, hundreds of genes were affected (both upregulated and downregulated) at P90, with further changes observed at P120 (Fig. 2n). In contrast, only a few genes were deregulated in the spinal cord at either time point (Fig. 2o). In nerve ECs, some differentially expressed genes (DEGs) changed progressively between P90 and P120, while others appeared only at a single time point, reflecting an evolving and partly stage-specific endothelial response.

Gene Ontology (GO) annotation of nerve DEGs and Gene Set Enrichment Analysis (GSEA) pointed to impaired BNB function, already evident at P90, consistent with early vascular leakage in *SOD1^G93A^* nerves (Fig. 2p-r and Suppl. Fig. 3z). This impairment was indicated by decreased expression of solute transporters (*Slco1c1*, *Slc2a1*, *Slc7a1*, *Slc19a3*, *Slc45a3*), key components of Wnt and Notch pathways required for barrier maintenance (*Axin2*, *Reck*, *Lef1*, *Dkk3*, *Sfrp5*, *Dll4*, *Hey1*, *Hes1*), as well as selected junction proteins (*Jcad*, *Tjp2*, *Cldn5*) and endothelial barrier modulators (*Bsg*, *Spock2*, *Mfsd2a*) (Fig. 2q, r). Downregulation of vasoprotective factors (*Klf2*, *Klf4*, *Igfbp3*, *Igfbp6*, *Lrp1*, *Plat*) indicated altered vascular homeostasis (Fig. 2r). Among metabolism-related DEGs, those involved in lipid biosynthesis, transport, and processing were especially affected. Decreased cholesterol availability was inferred from reduced expression of biosynthetic enzymes (*Hmgcs1*, *Hmgcr*, *Sqle*, *Msmo1*) and uptake receptors/transporters (*Ldlr*, *Abca2*, *Pltp*), along with increased expression of efflux transporters (*Abcg1*, *Scarb1*, *Abca1*) (Fig. 2r). A similar change in cholesterol homeostasis has been observed in *SOD1^G93A^* motor neurons, highlighting a shared metabolic imbalance^75^. In parallel, GO analysis of upregulated DEGs revealed an inflammatory profile in *SOD1^G93A^* nerve ECs, evident at P90 and more pronounced at P120 (Fig. 2p, r). This signature included pro-inflammatory cytokines/chemokines (*Il6*, *Cxcl1*, *Cxcl10*, *Ccl2*), receptors (*Il1r1*, *Tacr1/NK1*, *Tlr2*, *Trl3*), key signaling regulators –particularly within the TNFα-NFκB axis– and cell adhesion molecules (*Vcam1*, *Icam1*, *Sele*), which are expressed on activated endothelium and involved in immune cell invasion^76^ (Fig. 2r). Conversely, downregulation of a type-I interferon signature suggested suppression of immunoregulatory mechanisms active in capillaries of healthy nerves, as well as in the brain and other organs^64,77^ (Fig. 2q, r).

In summary, peripheral nerve ECs in *SOD1^G93A^* mice are more sensitive than those in the spinal cord, adopting a pro-inflammatory, activated state^76,78^ with concomitant BNB loss before axonal or motor neuron degeneration.

### Dysregulated transcytosis drives blood-nerve barrier breakdown

Several lines of evidence indicate that paracellular transport in the nerve endothelium remains relatively preserved during the early phase of ALS. Structural alterations in EC contacts were only evident in severe human cases (Fig.1h’, i), with only few junctional genes found downregulated in *SOD1^G93A^* mice (Fig. 2r). Furthermore, the distribution and levels of the adherens junction protein VE-cadherin (CDH5) did not change in *SOD1^G93A^* nerves (Suppl. Fig. 4d-f) and the TJ component ZO-1 decreased only in late-stage symptomatic mice (P140) (Suppl. Fig. 4g-i). In contrast, the increased density of cytoplasmic vesicles prior to disease onset (Fig. 1i-l) suggests that dysregulated transcellular transport may drive BNB impairment. To investigate this possibility, we immunostained *SOD1^G93A^* sciatic nerves for PLVAP, which forms stomatal and fenestral diaphragms on ECs, and regulates trans-endothelial transport^79^, and Claudin-5 (CLDN5), a TJ component expressed by barrier-forming ECs. In control nerves, PLVAP was restricted to permeable vessels of the epineurium, which correspondingly lacked CLDN5. Conversely, BNB endoneurial vessels, in which both paracellular and transcellular permeability are suppressed, expressed CLDN5 but not PLVAP (Fig. 3a)^60,64^. These non-overlapping patterns of PLVAP and CLDN5 were drastically altered in *SOD1^G93A^* nerves (Fig. 3b). At the pre-symptomatic stage (P60), ectopic expression of PLVAP was observed in endoneurial vessels that retained normal levels of CLDN5. PLVAP induction was initially focal and became progressively widespread at the symptomatic stage (P120) when CLDN5 expression began to decline (Fig. 3c-g, j, k). The aberrant upregulation of PLVAP in CLDN5^+^ ECs was exclusive to motor fiber-carrying nerves of *SOD1^G93A^* mice and absent in pure sensory nerves (Suppl. Fig. 4a-c). Notably, PLVAP was never induced in spinal cord microvessels, despite a localized decrease in CLDN5 in the ventral horn by P120, suggesting TJ alterations (Fig. 3h-i’, l, m). These results point to distinct mechanisms underlying the response of PNS and CNS barriers to ALS pathology.

**Fig. 3:**
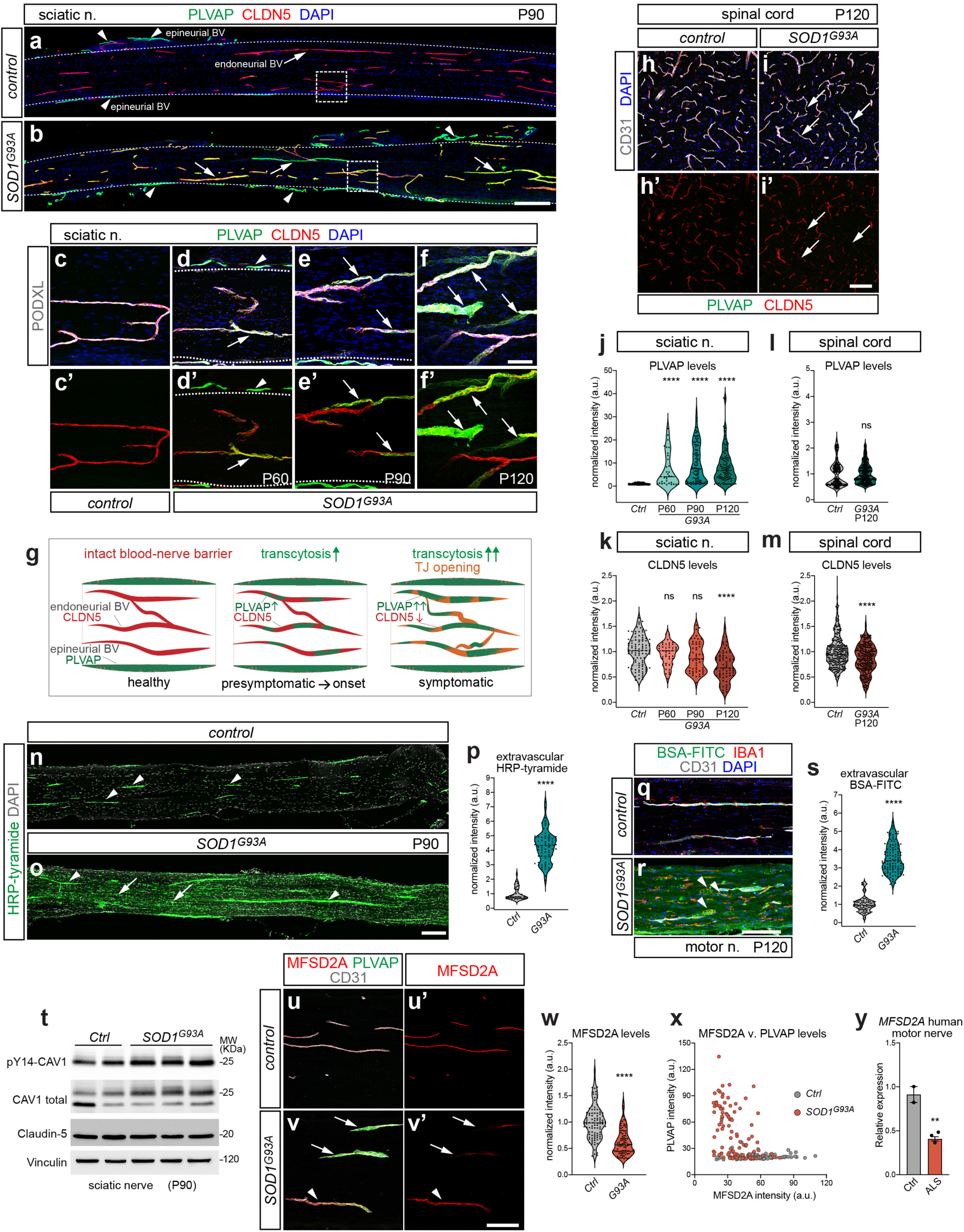
Transcytosis impairment in the nerve endothelium but not spinal cord of *SOD1^G93A^* ALS mice. **a**, **b**, Sciatic nerves from control (**a**) and *SOD1^G93A^*mice at P90 stained for PLVAP and CLDN5. DAPI, nuclei. CLDN5^+^ endoneurial vessels are PLVAP^−^ in controls (**a**, arrow) but express PLVAP in mutants (**b**, arrows). Permeable epineurial vessels express PLVAP in both genotypes (arrowheads). Dashed contours outline the endoneurium. Boxed areas in **a** and **b** are magnified in **c** and **e**, respectively. **c-f’**, Ectopic expression of PLVAP in endoneurial vessels of *SOD1^G93A^*sciatic nerves during disease course (P60, P90, P120) (**d-f’**, arrows), but not in controls (**c**). Podocalyxin (PODXL) marks all vessels; DAPI, nuclei. Lower panels (**c’-f’**) show only PLVAP and Claudin-5 staining. Dashed contours outline the endoneurium. Arrowheads point to epineurial vessels. **g**, Schematic of progressive BNB impairment in *SOD1^G93A^* nerves. Transcytosis increases at the presymptomatic stage, marked by focal PLVAP upregulation in endoneurial vessels, which transition into a hybrid PLVAP^+^/CLDN5^+^ state. At the symptomatic stage, PLVAP expression spreads and CLDN5 declines, indicating tight junction impairment. **h-i’**, Spinal cords from control (**h**, **h’**) and *SOD1^G93A^*(**i**, **i’**) mice at P120 stained for PLVAP and CLDN5. CD31 marks all vessels; DAPI, nuclei. Lower panels (**h’**, **i’**) show only PLVAP and CLDN5 staining. PLVAP is not upregulated in mutants, while focal tight junction loss is observed (**i**, **i’**, arrows). **j**, **k**, PLVAP (**j**) and CLDN5 (**k**) signal in endoneurial vessels of control and *SOD1^G93A^* sciatic nerves during disease progression, normalized to average control value. **l**, **m,** PLVAP (**l**) and CLDN5 (**m**) signal in spinal cord vessels of control and *SOD1^G93A^* mice at P120. **n**, **o**, Sciatic nerves from P90 control (**n**) of *SOD1^G93A^* (**o**) mice injected systemically with HRP, detected by Tyramide amplification. DAPI, nuclei. HRP is confined to vessels in controls (**n**, arrowheads) but extravasate in mutant nerves (**o**, arrows). **p**, Extravascular HRP-tyramide signal in control and *SOD1^G93A^*sciatic nerves (P90), normalized to average control value. **q**, **r**, Motor nerves from P120 control (**q**) and *SOD1^G93A^*(**r**) mice injected systemically with BSA-FITC. The tracer extravasates and is phagocytosed by macrophages (IBA1^+^) in mutants (**r**, arrowheads). CD31 marks vessels; DAPI, nuclei. **s**, Extravascular BSA-FITC signal in control and *SOD1^G93A^* motor/sciatic nerves (P120), normalized to average control value. **t**, Western blotting for phoshoY14-CAV1, total CAV1 and CLDN5 in sciatic nerve lysates from control and *SOD1^G93A^* mice at P90. Vinculin is a loading control. **u-v’**, Endoneurial sciatic nerve vessels from P120 control (**u**, **u’**) and *SOD1^G93A^* (**v**, **v’**) mice stained for MFSD2A, PLVAP and CD31. MFSD2A is shown separately in **u’**, **v’**. MFSD2A expression decreases in mutant vessels that upregulate PLVAP (arrows). A subset of vessels retain MFSD2A expression (arrowheads). **w**, MFSD2A signal in endoneurial vessels of P120 control and *SOD1^G93A^*sciatic nerves, normalized to average control value. **x**, Scatter plot showing MFSD2A (x-axis) and PLVAP (y-axis) signal intensity in individual endoneurial vessels of control and *SOD1^G93A^*sciatic nerve. **y**, qRT-PCR for *MFSD2A* in control and ALS *obturator* motor nerve biopsies. Mean (normalized to control) ± SEM. Scale bars: a, b, 500μm; c-f’, 50μm; h-i’, 100μm; n, o, 200μm; q, r, 100μm; u-v’, 100μm.

Endoneurial ECs that induced PLVAP expression at an early disease stage exhibited abnormal transcytosis. This was monitored through intravenous injection of horseradish peroxidase (HRP-tyramide) and fluorescent-albumin (BSA-FITC), established markers of transcellular transport^60,80–82^. In control mice, the tracers were confined to the lumen of endoneurial vessels, reflecting a low transcytosis rate (Fig. 3n, q). By contrast, in *SOD1^G93A^* sciatic and motor nerves —but not sensory nerves— both tracers diffused into the parenchyma and were subsequently phagocytosed by macrophages (Fig. 3o, p, r, s and Suppl. Fig. 4m-o). Leakage of plasma proteins in the endoneurium was further supported by accumulation of extravascular fibrinogen in *SOD1^G93A^*nerves by onset stage (Suppl. Fig. 4j-l). A main route of endothelial transcytosis occurs via caveolae^83^, whose formation is enhanced by Src-dependent phosphorylation of caveolin-1 (CAV1), their key structural component^84,85^. Consistent with increased transcytosis at the BNB, the levels of both total and phosphorylated (pY14) CAV1 were elevated in *SOD1^G93A^* sciatic nerves, while CLDN5 was unaffected (Fig. 3t). The subcellular distribution of CAV1 reflected BNB permeability changes: in control endoneurial ECs, CAV1 localized to the plasma membrane, similar to BBB and BSCB capillaries^86^, but redistributed to the cytoplasm in PLVAP^+^ ECs of *SOD1^G93A^* nerves (Suppl. Fig. 4s-v and data not shown). This pattern resembled that observed in wild-type dorsal root ganglia vessels, which are naturally permeable (PLVAP^+^) (Suppl. Fig. 4w-y). Increased caveolae-mediated transcytosis in *SOD1^G93A^* endoneurial vessels was further evidenced by a reduction in MFSD2A —a key inhibitor of the pathway^87,88^— correlating with PLVAP induction (Fig. 3u-x; see also Fig. 2r, heatmap), whereas MFSD2A expression in the spinal cord remained unchanged (Suppl. Fig. 4p-r). *MFSD2A* levels were also decreased in motor nerve biopsies from ALS patients (Fig. 3y).

We conclude that the early increase in BNB permeability in ALS nerves is primarily driven by a nerve-specific enhancement of transcytosis, which is not observed in the spinal cord.

### Peripheral nerves harbor ALS-susceptible capillaries with distinctive para-inflammatory tone

The observation that BNB leakage and PLVAP induction in endoneurial vessels of *SOD1^G93A^* mice first appeared in sparse hotspots raises the question of whether specific EC subtypes are inherently more susceptible to ALS. To address this, we performed droplet-based single-cell RNA-seq (scRNA-seq) on CD31^+^ ECs FACS-isolated from the sciatic nerves of onset-stage (P90) *SOD1^G93A^* mice and non-transgenic controls (Fig. 4a, b). Unsupervised subclustering of BNB-forming endoneurial ECs (*Cldn5*^+^) identified a cell subset with increased *Plvap* expression in *SOD1^G93A^* samples (Fig. 4c-e). This activated population corresponded to capillary-venous ECs expressing carbonic anhydrase *Car4*^64,77^ (Fig. 4e, f and Suppl. Fig. 5a, b). The other two endoneurial clusters were “capillary-barrier” ECs, characterized by junctional and transport genes (*Scl2a1*, *Mfsd2a*, *Spock2*), and arterial ECs (*Gja4*, *Stmn2*, *Sema3g*) (Fig. 4f and Suppl. Fig. 5b). The remaining EC clusters were assigned to epineural vessels (*Plvap*^+^ in controls), including veins (*Nr2f2*, *Ackr1*, *Sepl*, *Vwf*) and capillaries (*Rgcc*, *Cd300lg*, *Thrsp*, *Cxcl12*; “capillary-Plvap^+^”). A smaller “capillary-Plvap^−^” cluster, possibly associated with epi/perineurial compartments^64^, lacked *Plvap* and expressed genes such as *Gpihbp1*, *Aqp7* and *Timp4* (Fig. 4f and Suppl. Fig. 5b). The relative abundance of EC subtypes was similar between *SOD1^G93A^* and control nerves, except for a modest expansion of the capillary-venous cluster (p=0.0495) (Fig. 4g). Most genes upregulated in *SOD1^G93A^* ECs were found in the capillary-venous subtype, whereas downregulated genes were more prominent in capillary-barrier ECs (Fig. 4h), as confirmed by Expression Weighted Celltype Enrichment (EWCE) analysis of bulk RNA-seq DEGs (Fig. 4i).

**Fig. 4:**
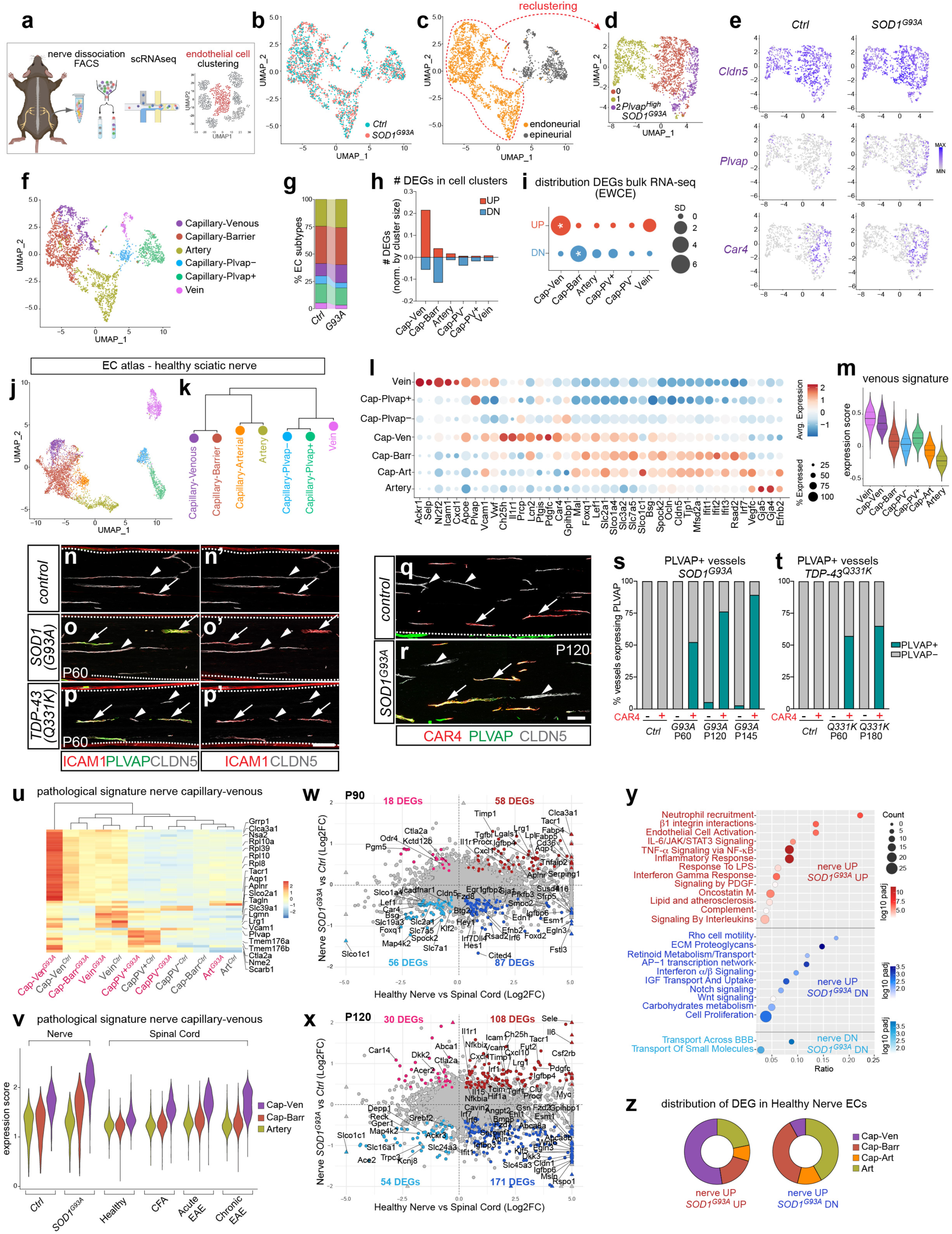
Primed state and vulnerability of capillary subtypes in peripheral nerves of ALS mice. **a**, Workflow for scRNA-seq of mouse sciatic nerve endothelium. **b**, **c**, Uniform manifold approximation and projection (UMAP) plot of integrated control and *SOD1^G93A^* nerve ECs, color coded by genotype (**b**) or by endoneurial (orange) vs. epineurial (gray) identity (**c**). The dataset comprises 2611 cells (control, 1250 cells; *SOD1^G93A^*, 1361cells). **d**, Reclustering of endoneurial ECs from integrated control and *SOD1^G93A^* dataset. Cluster 2 has high *Plvap* expression in *SOD1^G93A^*. **e**, UMAP plots showing expression of *Cldn5*, *Plvap* and *Car4* in control and *SOD1^G93A^* endoneurial ECs. **f**, UMAP plot of integrated control and *SOD1^G93A^* dataset color-coded by EC subtype. Capillary-venous ECs correspond to venous-Plvap^−^ ECs annotated in a previous nerve dataset{Bhat, 2024 #1}. **g**, Relative abundance of EC subtypes in control and *SOD1^G93A^* nerves. **h**, Number of upregulated (red) and downregulated (blue) DEGs (padj <0.1) identified by scRNA-seq in nerve EC subtypes in *SOD1^G93A^*vs. control, normalized by cluster size. **i**, Distribution of upregulated (red) and downregulated (blue) DEGs (padj <0.1, concordant at P90-P120) from bulk RNA-seq of *SOD1^G93A^*vs. control sciatic nerve ECs computed by EWCE. **j**, UMAP plot of ECs from healthy sciatic nerve color-coded by subtype. The atlas comprises 6 independent scRNA-seq datasets. The capillary-arterial subtype was not resolved in our scRNA-seq dataset alone (Fig. 4f) **k**, Dendrogram for hierarchical clustering of EC subtypes from healthy sciatic nerve atlas based on the top 200 markers for each celltype. **l**, Dot plot showing the expression of EC subtype markers from healthy sciatic nerve atlas. **m**, Expression of “venous signature” in nerve EC subtypes from healthy sciatic nerve atlas. The gene set includes 58 venous markers identified in various organs{Kalucka, 2020 #2} with padj <10^-8^ in nerve capillary-venous ECs. **n-p’**, ICAM1/CLDN5 co-staining identifies venous capillaries (arrows) in sciatic nerves from P60 control (**n**, **n’**), *SOD1^G93A^* (**o**, **o’**) and *TDP-43^Q331K^*(**p**, **p’**) mice. Venous capillaries become PLVAP^+^ in mutants. CLDN5^+^/ICAM1^−^ vessels remain PLVAP^−^ (arrowheads). **n’, o’**, **p’** show only ICAM1 and CLDN5 staining. Dashed contours outline the endoneurium. **q-r**, CAR4/CLDN5 co-staining identifies venous capillaries (arrows) in sciatic nerves from P120 control (**q**) and *SOD1^G93A^* (**r**) mice. CAR4^+^ vessels become PLVAP^+^ in *SOD1^G93A^*. CAR4^−^ vessels remain PLVAP^−^ (arrowheads) Dashed contours outline the endoneurium. **s**, **t**, Percentage of CAR4^+^ vs. CAR4^−^ endoneurial vessels expressing PLVAP in control, *SOD1^G93A^* (**s**) or *TDP-43^Q331K^* (**t**) sciatic nerves at the indicated stages. **u**, Expression heatmap (z-score) of capillary-venous pathological signature across EC subtypes from controls and *SOD1^G93A^* nerves. The signature includes 50 genes upregulated in *SOD1^G93A^* capillary-venous ECs (padj <0.05). The dendrogram illustrates hierarchical clustering of EC subtypes in the two conditions. **v**, Expression of the nerve capillary-venous pathological signature across EC subtypes from nerve and spinal cord in the indicated conditions. **w**, **x**, Log2 fold-change (Log2FC) vs. Log2FC scatterplot comparing DEGs (padj <0.1, colored) in ECs from healthy nerve vs. spinal cord (x-axis) and *SOD1^G93A^* vs. control nerve (y-axis) at P90 (**w**) and P120 (**x**). Selected DEGs are labeled. **y**, GO analysis of DEGs (P90 and P120 combined) from the comparisons in **w**, **x**. Genes downregulated in healthy nerve and upregulated in *SOD1^G93A^*(top/left quadrants) did not yield significant GO results. **z**, Distribution of DEGs (from comparisons in **w**, **x**) across EC subtypes in healthy sciatic nerve, based on expression strength index (ESI, see Methods). Scale bars: n-p’, 100μm; q, r, 100μm.

To better characterize steady-state features of these EC subtypes, we generated a scRNA-seq atlas of the healthy nerve endothelium by integrating our control samples with five published adult mouse sciatic nerve datasets^64,89–92^. This yielded a total of ∼5000 ECs, divided into seven clusters, which were annotated based on previously defined markers^64,77^ (Fig. 4j and Suppl. Fig. 5e). The four endoneurial EC subtypes exhibited an arteriovenous zonation similar to the brain vasculature^93^, encompassing capillary-venous, capillary-barrier, capillary-arterial, and arterial populations (Fig. 4j). Hierarchical clustering revealed a close relationship between capillary-venous and capillary-barrier ECs, the two subtypes most affected in *SOD1^G93A^* nerves (Fig. 4k). Accordingly, capillary-venous ECs expressed vascular barrier genes (*Cldn5, Ocln*, *Tjp1*, *Bsg*, *Spock2*, *Mfsd2a*, *Lef1*, *Scl2a1*), though often at lower levels than *bona fide* barrier capillaries (Fig. 4l and Suppl. Fig. 5e, f). At the same time, they exhibited a venous phenotype defined by highly conserved markers (*Apoe*, *Vcam1*, *Vwf*, *Ctla2a*, *Ptgs1*, *Tmsb10*, *Icam1*), along with others shared across venous beds in different organs (*Car4*, *Lcn2*, *Ch25h*, *Ptgis*, *Adgrg6*)^77^ (Fig. 4l, m and Suppl. Fig. 5e, f). In addition, enrichment of ribosomal protein genes suggested elevated basal translational activity (Suppl. Fig. 5e).

Venous capillaries in the sciatic nerve were identified by co-staining for the venous adhesion molecule ICAM1^77^ and the barrier marker CLDN5 (Fig. 4l, n, n’ and Suppl. Fig. 5e). Consistent with the predicted susceptibility of this EC subtype, PLVAP upregulation was restricted to ICAM1^+^/CLDN5^+^ vessels in pre-symptomatic *SOD1^G93A^*and *TDP-43^Q331K^* mice (Fig. 4o-p’). However, because ICAM1 levels increased in mutants (Fig. 5b and Suppl. Fig. 11h), we instead labeled venous capillaries with CAR4, which remained stable throughout disease progression (Fig. 4e, l, q). As expected, ectopic PLVAP expression was again confined to CAR4^+^/CLDN5^+^ capillaries in both *SOD1^G93A^*and *TDP-43^Q331K^* models at pre-symptomatic stage, and this association persisted as the proportion of PLVAP^+^ endoneurial vessels rose in the symptomatic phase (Fig. 4r-t, Suppl. Fig. 5g-m’).

**Fig. 5:**
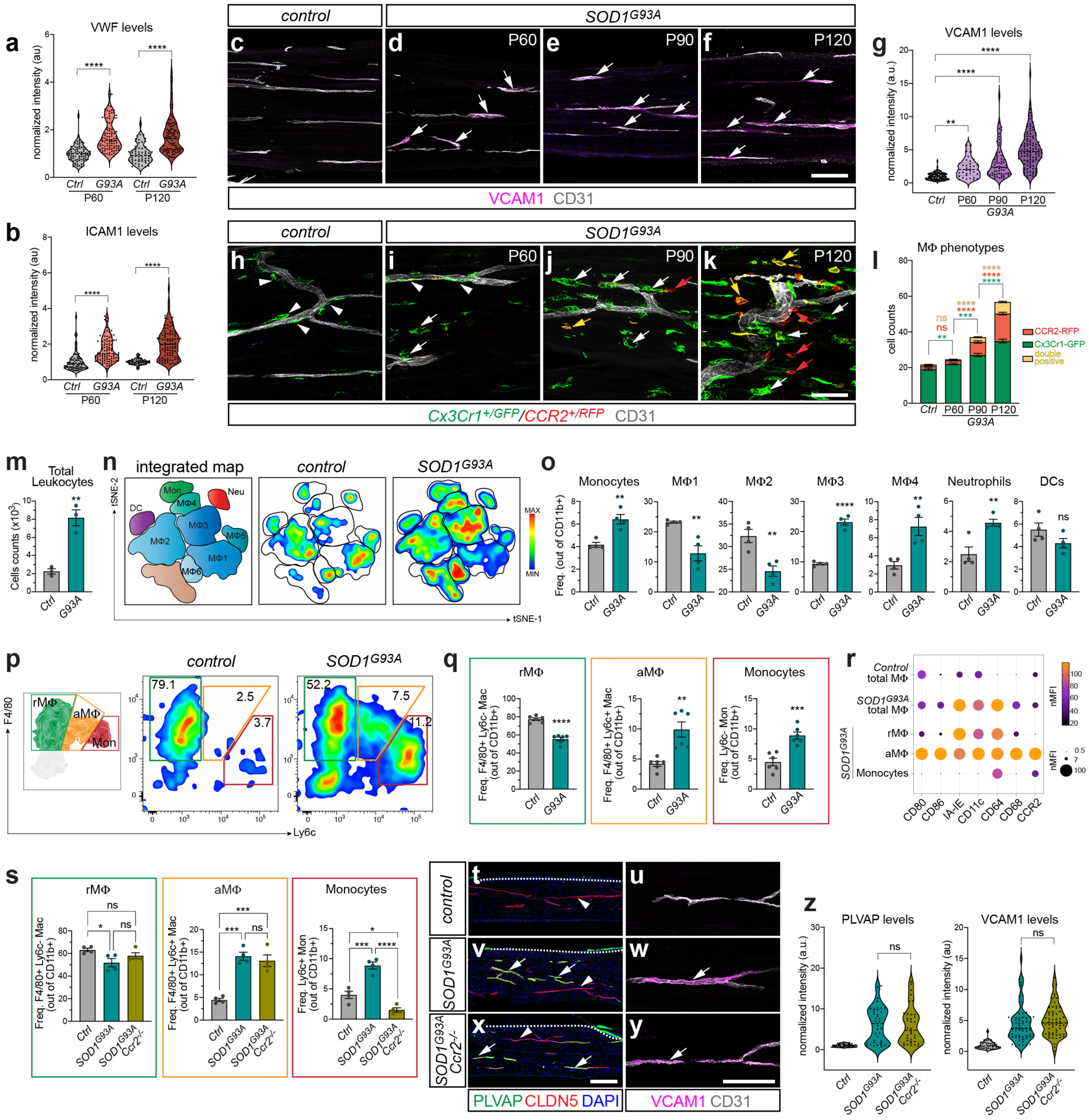
Endothelial activation in peripheral nerves of *SOD1^G93A^* ALS mice parallels myeloid remodeling but is independent of monocyte infiltration. **a**, **b**, VWF (**a**) and ICAM1 (**b**) signal in endoneurial vessels of control and *SOD1^G93A^* sciatic nerves at P60 and P120, normalized to average control value. **c-f**, Sciatic nerves from control (**c**) and *SOD1^G93A^* (**d-f**) mice at the indicated stages stained for VCAM1. CD31 marks all vessels. VCAM1 is upregulated in endoneurial vessels of mutants (arrows). **g**, VCAM1 signal in endoneurial vessels of control and *SOD1^G93A^*sciatic nerves during disease progression, normalized to average control value. **h-k**, Sciatic nerves of control (**h**) and *SOD1^G93A^* (**i-k**) mice at the indicated stages, expressing *Cx3cr1^GFP^*/*Ccr2^RFP^*reporter. CD31 marks vessels. In controls, elongated GFP^+^ MΦ associate with vessels (**h**, arrowheads). In *SOD1^G93A^* nerves, rare globular GFP^+^ MΦ appear at P60 (**i**, white arrows) and increase in the parenchyma at P90 (**j**) and P120 (**k**). RFP^+^ monocytes infiltrate *SOD1^G93A^* nerves at P120 (**k**, red arrows); some acquire GFP expression (**k**, yellow arrows). Panels are magnified from Suppl. Fig. 7l. **l**, Number of GFP^+^, RFP^+^, GFP^+^/RFP^+^ double-positive MΦ in control and *SOD1^G93A^* sciatic nerves at the indicated stages. Mean ± SEM. **m**, Number of leukocytes (CD45^+^) measured by flow cytometry in control and *SOD1^G93A^* sciatic nerves at P120. Mean ± SEM. **n**, t-SNE density maps of multi-parametric flow cytometry profiling of sciatic nerve myeloid cells showing changes in abundance of discrete populations in *SOD1^G93A^* mice compared to controls (P120). MΦ subsets 1-6; Mon, monocytes; DC, dendritic cells; Neu, neutrophils. **o**, Frequency of myeloid populations identified by unsupervised t-SNE analysis (gated from CD11b^+^ cells). Small MΦ5 and MΦ6 clusters that did not display significant changes in mutants are not shown. Mean ± SEM. **p**, Flow cytometry plots from control and *SOD1^G93A^* (P120) sciatic nerves showing resident macrophages (rMΦ, green gate), activated macrophages (aMΦ, organe gate), and monocytes (Mon, red gate) discriminated based on F4/80 and Ly6c levels (schematic, left panel). **q**, Frequency of rMΦ, aMΦ and monocytes (gated from CD11b^+^ cells) in control and *SOD1^G93A^* nerves. Mean ± SEM. **r**, Normalized mean fluorescence intensity (MFI) of selected inflammatory markers upregulated in *SOD1^G93A^*nerve macrophages. Since aMΦ and monocytes are nearly absent in control nerves, the total F4/80^+^ population is used for comparison with *SOD1^G93A^*. **s**, Frequency of rMΦ, aMΦ and monocytes (gated from CD11b^+^ cells) in control, *SOD1^G93A^*, and *SOD1^G93A^; Ccr2^-/-^* sciatic nerves. Mean ± SEM. Flow cytometry plots are shown in Suppl. Fig. 8f. **t-y**, Sciatic nerves from control (**t**, **u**), *SOD1^G93A^* (**v**, **w**), and *SOD1^G93A^; Ccr2^-/-^* (**x**, **y**) mice at P120 stained for PLVAP and CLDN5 (DAPI, nuclei) (**t**, **v**, **x**), or VCAM1 (CD31, all vessels) (**u**, **w**, **y**). Dashed contours outline the endoneurium. PLVAP and VCAM1 are upregulated in CNDN5^+^ vessels in *SOD1^G93A^* and *SOD1^G93A^; Ccr2^-/-^* nerves (arrows). Some vessels remain PLVAP^−^ (arrowheads). **z**, PLVAP and VCAM1 signal in endoneurial vessels of control, *SOD1^G93A^*, and *SOD1^G93A^; Ccr2^-/-^* sciatic nerves, normalized to average control value. Scale bars: c-f, 100μm; h-k, 50μm; t, v, x, 200μm; u, w, y, 100μm.

Next, we defined a “pathological” signature enriched in *SOD1^G93A^* capillary-venous ECs, comprising genes linked to vascular inflammation, permeability and activation (*Plvap*, *Vcam1*, *Lrg1*, *Tacr1*), along with others with less defined endothelial functions (e.g., water channel-forming *Aqp1*) and ribosomal protein genes (Fig. 4u and Suppl. Fig. 5c, d). *SOD1^G93A^* capillary-barrier ECs also upregulated this gene set, albeit to a lesser extent (Fig. 4u). Unexpectedly, based on this signature control capillary-venous ECs clustered with mutant barrier ECs (Fig. 4u), suggesting a basal state of activation further amplified in disease.

To assess whether the ALS signature identified in nerve capillary-venous ECs was shared across other pathological conditions, we turned to experimental autoimmune encephalomyelitis (EAE), a model characterized by BSCB breakdown and inflammatory changes primarily affecting venous ECs^94^. Using scRNA-seq data from spinal cord ECs of healthy controls, Freund’s adjuvant (CFA)-immunized mice, and mice with acute or chronic EAE, we annotated *Cldn5*^+^ EC clusters corresponding to capillary-venous, capillary-barrier, and arterial subtypes (Suppl. Fig. 6a-c)^94^. Among these, only venous capillaries in CFA and EAE spinal cords, but not in healthy controls, upregulated the pathological gene set, though at much lower levels than in nerves (Fig. 4v). While key mediators of vascular inflammation and damage, such as *Vcam1* and *Lrg1*, were induced in both tissues, other genes were specific to venous capillaries in *SOD1^G93A^* nerves (e.g., *Plvap*, *Tacr1*, *Aqp1*, *Aplnr*) (Suppl. Fig. 6d). That venous ECs in CFA mice paralleled changes seen in EAE (Fig. 4v) underscores the sensitivity of the signature, as CFA/pertussis toxin —used as a control for EAE— *per se* increases vascular permeability and immune activation^95–97^. Furthermore, this comparison reinforced the notion that capillary-venous ECs in healthy nerves exhibit an activated gene profile, comparable to spinal cords of CFA-treated and EAE mice (Fig. 4v).

We therefore asked whether the nerve vasculature exhibits a basal activation state. Compared with spinal cord ECs, healthy nerve ECs were enriched for inflammatory pathways (Fig. 2m and Suppl. Fig. 6f), which were further upregulated in *SOD1^G93A^* (Fig. 4w, x, top-right quadrants and Fig. 4y) and predominantly detected in capillary-venous ECs (Fig. 4z). In contrast, nerve-enriched genes downregulated in *SOD1^G93A^*were associated with vascular homeostasis and enriched in capillary-barrier and arterial ECs (Fig. 4w, x, bottom-right quadrants; Fig. 4y, z).

Altogether, these findings suggest that the nerve endothelium maintains a basal para-inflammatory tone^98^, partly driven by the distinct transcriptional profile of capillary-venous ECs compared with their spinal cord counterparts (Suppl. Fig. 6e), possibly making the nerve vasculature more susceptible to pathological changes in ALS.

### Endothelial activation precedes and is independent of monocyte infiltration in ALS nerves

In line with our transcriptional analysis pointing to inflammatory activation of the nerve endothelium, immunostaining of *SOD1^G93A^*sciatic nerves revealed increased expression of VWF, ICAM1, and VCAM1—key mediators of vascular inflammation and immune cell recruitment— in endoneurial microvessels from the pre-symptomatic stage (P60) onward (Fig. 5a-g and Suppl. Fig. 7a-e”). To explore the spatiotemporal relationship between EC activation and remodeling of the myeloid compartment in peripheral nerves, we crossed *SOD1^G93A^* with the *Cx3cr1^GFP^*/*Ccr2^RFP^*dual-reporter allele (Fig. 5h-k and Suppl. Fig. 7l). This model distinguished GFP^+^ resident macrophages expressing fractalkine receptor CX3CR1 from RFP^+^ infiltrating inflammatory monocytes expressing CCR2, the receptor for CCL2 —a monocyte chemoattractant upregulated in *SOD1^G93A^* nerve ECs (Fig. 2r). In control sciatic nerves, GFP^+^ resident macrophages with an elongated morphology were closely associated with endoneurial vessels, while blood-derived RFP^+^ monocytes were virtually absent (Fig. 5h). In *SOD1^G93A^* nerves, GFP^+^ macrophages began to increase in numbers at the presymptomatic stage (P60) (Fig. 5i, l). By disease onset (P90), many had adopted a globular shape consistent with phagocytic activity and were scattered in the nerve parenchyma, away from vessels (Fig. 5j, l). At this stage, RFP^+^ monocytes began infiltrating, and by the early-symptomatic phase (P120) both populations had expanded further. A subset of RFP^+^ monocytes co-expressed GFP, reflecting their differentiation into activated macrophages (Fig. 5k, l).

We concluded that in *SOD1^G93A^* nerves carrying motor fibers (*sciatic* and *quadriceps n.*), but not in sensory nerves, EC activation marked by VCAM1 induction coincided with the early expansion and activation of resident macrophages, whereas infiltration of inflammatory monocytes occurred later (Fig. 5g, l and Suppl. Fig. 7n-t). In contrast, EC activation was not observed in the spinal cord of symptomatic *SOD1^G93A^* mice, as VCAM1 and VWF were not induced in the microvasculature but remained confined to larger veins (Suppl. Fig. 7f-k). At this stage, microglia marked by *Cx3cr1^GFP^*expanded without recruitment of *Ccr2^RFP^* monocytes (Suppl. Fig. 7m).

Given the activated EC profile and increased leukocyte counts in *SOD1^G93A^*peripheral nerves (Fig. 5m), we used high-dimensional spectral flow cytometry to profile the immune compartment in sciatic nerves from control and mutant mice (P120) (Fig. 5n, o and Suppl. Fig. 8a). The expansion of leukocytes was not due to lymphocytic infiltration, as B cells (CD19^+^) and T cells (CD4^+^ and CD8^+^) were not detected (Suppl. Fig. 8b-e). Instead, unsupervised analysis of myeloid composition using t-distributed stochastic neighbor embedding (t-SNE) dimensionality reduction revealed a substantial increase in the frequencies of monocytes, specific macrophage subpopulations, and neutrophils in *SOD1^G93A^* nerves, while dendritic cells remained stable (Fig. 5n-o). Macrophage clusters MF1 and MF2, exhibiting a resting phenotype, were reduced in *SOD1^G93A^* mice, whereas MF3 and MF4 with an activated phenotype —marked by high expression of IA-IE, CD64, CD68, CX3CR1, CD86 and CD80— were enriched (Fig. 5n, o and Suppl. Fig. 8a). Considering the role of infiltrating monocytes and macrophages in modulating ALS progression^53,54,99,100^, we conducted a supervised analysis to further characterize these lineages, distinguishing three main subpopulations based on F4/80 and Ly6c expression: tissue-resident MF (rMF; F4/80^+^ Ly6c^−^), activated MF (aMF) displaying intermediate Ly6c levels (F4/80^+^ Ly6c^low^), and monocytes (F4/80^low^ Ly6c^hi^) (Fig. 5p). In healthy sciatic nerves, most CD11b^+^ myeloid cells were rMF, but this subpopulation declined in *SOD1^G93A^* mice (Fig. 5p, q) and began to express pro-inflammatory markers such as IA-IE, CD11c and CD64 (Fig. 5r). In parallel, inflammatory monocytes increased, along with a substantial expansion of the aMF subset, which upregulated several pro-inflammatory molecules (e.g., CD80, CD86, IA-IE, CD11c, CD64, CD68 and CCR2) (Fig. 5r). The aMΦ population likely comprised both infiltrating monocyte-derived macrophages (CCR2^+^) and resident macrophages that may have transitioned toward a pro-inflammatory state, in contrast to the pro-resolving traits typically associated with Ly6c^low^ macrophages in other contexts^53,54,101,102^. Collectively, these findings reveal a shift of the macrophage compartment toward an activated state in ALS-affected nerves.

In light of these dynamics, we investigated whether EC activation was sustained by monocytes/macrophages infiltration by crossing *SOD1^G93A^*mice with a *Ccr2* knockout allele, which prevents monocyte recruitment to the sites of inflammation^103^. As expected, the numbers of inflammatory monocytes in sciatic nerves were drastically reduced in *SOD1^G93A^; Ccr2^-/-^* mice compared to *SOD1^G93A^* mutants at P120 (Fig. 5s and Suppl. Fig. 8f). However, there were no significant differences in the frequencies of rMΦ and aMΦ populations in the two models, further supporting the idea that aMΦ were not derived from inflammatory monocytes, but likely emerged from rMΦ that had acquired a pro-inflammatory profile (Fig. 5s and Suppl. Fig. 8f). Notably, despite near-complete absence of inflammatory monocytes and their derivatives in *SOD1^G93A^; Ccr2^-/-^* sciatic nerves, EC activation and BNB impairment — reflected by ectopic induction of VCAM1 and PLVAP on endoneurial vessels, respectively— occurred to the same extent as in *SOD1^G93A^* mice (Fig. 5t-z).

Thus, ALS-induced vascular inflammation in peripheral nerves precedes extensive reconfiguration and activation of tissue-resident macrophages and develops independently of circulating monocyte infiltration, which takes place only after disease onset. Whether the earliest instances of focal EC activation and resident macrophage responses are interdependent or just temporally aligned remains to be addressed.

### Neutrophils sustain endothelial activation and contribute to axonal pathology in ALS peripheral nerves

High-dimensional unsupervised and supervised spectral flow cytometry analysis of leukocytes in sciatic nerve revealed a significant increase in neutrophils (CD11b^+^ Ly6G^+^) in *SOD1^G93A^*mice at P120 (Fig. 5n, o; Fig. 6a, b). Since this approach did not differentiate between circulating and extravasated neutrophils, we performed tissue immunostaining for Ly6g and Ly-6B.2 (7/4) to assess neutrophil infiltration into the nerve parenchyma. In control mice, extravasated neutrophils were virtually absent. In contrast, they were present in the sciatic and motor nerves of *SOD1^G93A^* mice as early as P60 and became increasingly abundant by disease onset (P90) (Fig. 6c, d). However, in the spinal cord of mutants, neutrophils remained rare, even at advanced stages, and were confined to the vessel lumen or meninges without infiltrating the parenchyma (Suppl. Fig. 9a-c). The accumulation of neutrophils in *SOD1^G93A^* peripheral nerves after P90 followed the early induction of inflammatory markers on endoneurial vessels, which became evident at P60 (Fig. 5d), suggesting that their initial ingress from the circulation was mediated by focal activation of the nerve endothelium^104,105^.

**Fig. 6:**
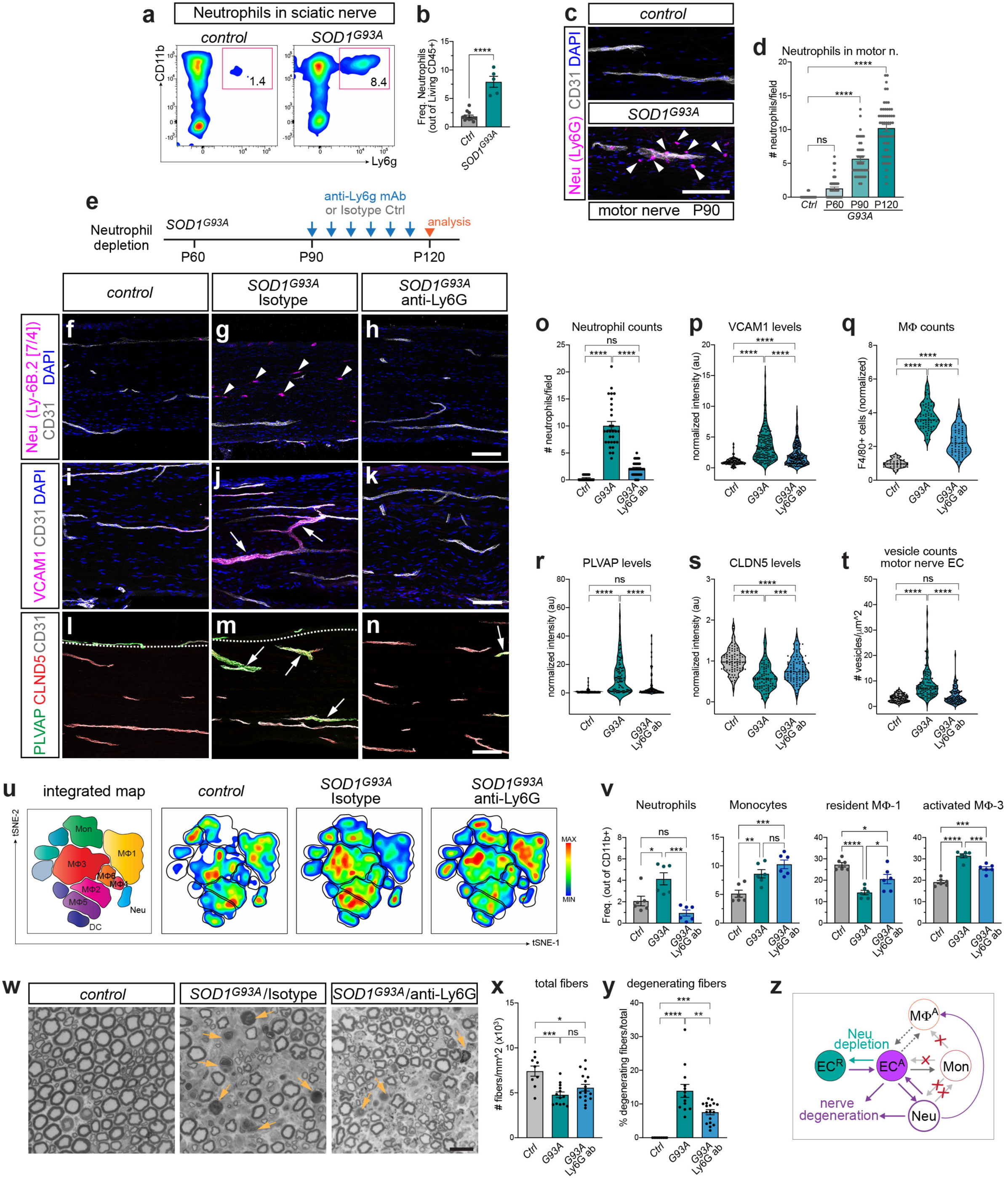
Neutrophil depletion reverses vascular damage and mitigates axonal degeneration in *SOD1^G93A^*ALS mice. **a**, **b**, Representative flow cytometry plots (**a**) and corresponding frequency measurements (**b**) showing increase of neutrophils (CD11b^+^ Ly6G^+^) in sciatic nerves of *SOD1^G93A^*mice at P120. **c**, Neutrophils (Ly6G^+^, arrowheads) infiltrate SOD1G93A motor nerves at P90. CD31 marks vessels; DAPI, nuclei. **d**, Number of extravasated neutrophils in motor nerves of control and *SOD1^G93A^*mice at different stages. Mean ± SEM. **e**, Timeline of neutrophil depletion with anti-Ly6G treatment. **f-h**, Neutrophils detected by Ly-6B.2 (7/4) staining in sciatic nerves of control (**f**), *SOD1^G93A^*/isotype antibody-treated (**g**) and *SOD1^G93A^*/anti-Ly6G-treated (**h**) mice at P120. Infiltrating neutrophils (arrowheads) are depleted by anti-Ly6G treatment. CD31, vessels; DAPI, nuclei. **i-k**, VCAM1 induction in endoneurial vessels of *SOD1^G93^*sciatic nerves (**j**, arrows; compare to control mice, **i**) is rescued by anti-Ly6G treatment (**k**). **l-n**, PLVAP upregulation in CLDN5^+^ vessels of *SOD1^G93A^*sciatic nerves (**m**, arrows; compare to control mice, **l**) is rescued by anti-Ly6G treatment (**n**). Dashed contours outline the endoneurium. **o**, Neutrophil counts (Ly-6B.2 7/4 and Ly6G staining) in sciatic nerves of control, *SOD1^G93A^*/isotype-treated and *SOD1^G93A^*/anti-Ly6G-treated mice at P120. Mean ± SEM. **p**, VCAM1 signal in sciatic nerve endoneurial vessels at P120, normalized to average control value. **q**, Macrophages counts (F4/80 staining) in sciatic nerves at P120, normalized to average control value. **r**, **s**, PLVAP (**r**) and CLDN5 (**s**) signal in sciatic nerve endoneurial vessels at P120, normalized to average control value. **t**, Vesicle density in motor nerve ECs at P120 assessed by TEM. **u**, t-SNE density maps of multi-parametric flow cytometry profiling of myeloid cells from sciatic nerves of control, *SOD1^G93A^*/isotype-treated and *SOD1^G93A^*/anti-Ly6G-treated mice at P120. MΦ subsets 1-6; Mon, monocytes; DC, dendritic cells; Neu, neutrophils. **v**, Frequency of neutrophils, monocytes, resident (non-activated) MΦ1 and activated MΦ3 subsets identified by unsupervised t-SNE analysis (gated from CD11b^+^ cells). Mean ± SEM. **w**, Toluidine Blue-stained semithin sections of *quadriceps* (motor) nerves from control, *SOD1^G93A^*/isotype-treated and *SOD1^G93A^*/anti-Ly6G-treated mice at P120. Degenerating fibers (arrows) are reduced by anti-Ly6G treatment. **x**, **y**, Axon density (**x**) and percentage of degenerating axons (**y**) assessed in semithin sections from control and *SOD1^G93A^* motor nerves. Mean ± SEM. **z**, Model: an exacerbating loop between nerve EC activation (EC^A^) and neutrophil (Neu) infiltration promotes vascular damage and neurodegeneration. EC dysfunction reverses upon neutrophil depletion (EC^R^, resting). Neutrophils amplify EC responses and modulate activation of tissue-resident macrophages (MΦ^A^) independently of monocyte (Mon) infiltration. Monocytes do not contribute to EC activation. Potential interactions between EC and MΦ activation remain to be defined. Scale bars: c, 100μm; f-n, 100μm; w, 20μm.

Given that infiltrating neutrophils are known to increase vascular permeability^106,107^, we tested their contribution to sustained EC activation and BNB dysfunction. *SOD1^G93A^*mice were treated with an anti-Ly6G depleting antibody, or isotype control via intraperitoneal injection from P90 to P120, targeting the window of significant neutrophil accumulation (Fig. 6e). The treatment efficiently depleted neutrophils from blood and nerve tissue, as confirmed by flow cytometry and immunofluorescence for the Ly-6B.2 (7/4) neutrophil marker, which showed a substantial reduction of neutrophil infiltration in *SOD1^G93A^* sciatic nerves (Fig. 6f-h, o and Suppl. Fig. 9d-f). Neutrophil depletion significantly improved vascular pathology in sciatic and motor nerves of *SOD1^G93A^* mice. The aberrant induction of VCAM1 and PLVAP in endoneurial vessels was substantially attenuated, while CLDN5 expression was partially rescued in anti-Ly6G-treated mutants (Fig. 6i-n, p, r, s and Suppl. Fig. 9g-o). Furthermore, TEM revealed a marked reduction in pinocytotic vesicles in motor nerve ECs, along with restored, sealed TJs (Fig. 6t and Suppl. Fig. 9p-r’). Thus, preventing neutrophil infiltration in peripheral nerves promotes vascular integrity.

We next evaluated the relationship between neutrophil infiltration and activation of the myeloid compartment in *SOD1^G93A^* nerves. Neutrophil depletion mitigated macrophage dysregulation, leading to a reduction in F4/80^+^ MΦ counts in the affected tissues (Fig. 6q and Suppl. Fig. 10a-c). High-dimensional unsupervised spectral flow cytometry revealed a reduction in the activated macrophage subset (MF3), accompanied by a corresponding increase in the resting resident pool (MF1), while the frequency of inflammatory monocytes remained unchanged (Fig. 6u-v and Suppl. Fig. 10f-h). Supervised analysis further confirmed the increase in rMΦ (F4/80^+^ Ly6C^−^) and a parallel decrease in aMΦ (F4/80^+^ Ly6C^low^) subpopulations, with no significant changes in infiltrating monocytes (Suppl. Fig. 10d, e). Conversely, neutrophil numbers and extravasation remained elevated in the sciatic nerves of *SOD1^G93A^; Ccr2^-/-^* mice, indicating that neutrophil recruitment occurred independently of monocyte infiltration (Suppl. Fig. 10i-n).

Because vascular inflammation and BNB impairment improved upon neutrophil depletion, we examined whether this translated into amelioration of ALS-like phenotypes in *SOD1^G93A^* mice. Semithin motor nerve sections revealed fewer degenerating axon fibers in neutrophil-depleted mutants compared to untreated controls at the symptomatic stage (P120), although total axon numbers remained unchanged (Fig. 6w-y). Amelioration of axonal pathology was accompanied by a moderate but significant preservation of motor neurons, while astrogliosis and microgliosis remained elevated (Suppl. Fig. 9s-u). However, these improvements did not alter overall disease progression in *SOD1^G93A^* mice, as motor performance — assessed by grip strength and rotarod tests— and body weight decline were not rescued (Suppl. Fig. 9v-x).

We conclude that EC activation in ALS peripheral nerves is a potentially reversible process, as shown by its attenuation upon neutrophil depletion. Our findings suggest a detrimental feedback loop between EC activation and neutrophil infiltration that perturbs resident macrophages, disrupts the BNB, and accelerates axonal degeneration independently of monocyte recruitment (Fig. 6z).

### Non-cell autonomous impact of ALS mutations on the nerve endothelium

We have detected ultrastructural abnormalities accompanied by early upregulation of PLVAP in the nerve endothelium of *TDP-43^Q331K^* mice, similar to *SOD1^G93A^* mutants, demonstrating that these changes are not limited to an individual ALS model (Fig.1l, r and Fig. 4t). Unlike *SOD1^G93A^*, which is ubiquitously expressed^69^, *TDP-43^Q331K^* is driven by the murine prion promoter, active in neurons and glia^68,108,109^. Immunostaining for Myc-tagged *TDP-43^Q331K^* showed nuclear localization in spinal cord parenchymal cells, but never in ECs (Fig. 7a, b, e). A similar pattern was observed in the sciatic nerve, where rare Myc^+^ nuclei—possibly fibroblasts— were detected, with no expression in ECs (Fig. 7c-e). These findings were further confirmed by quantitative RT-PCR for human *TDP-43^Q331K^* on whole sciatic nerve or spinal cord extracts. In line with the immunostaining results, mutant transcript levels were higher in the spinal cord than in the sciatic nerve but remained undetectable in FACS-purified ECs from either tissue (Fig. 7f). We concluded that *TDP-43^Q331K^* is absent from ECs in the CNS and PNS, consistent with the reported activity of the transgenic promoter.

**Fig. 7:**
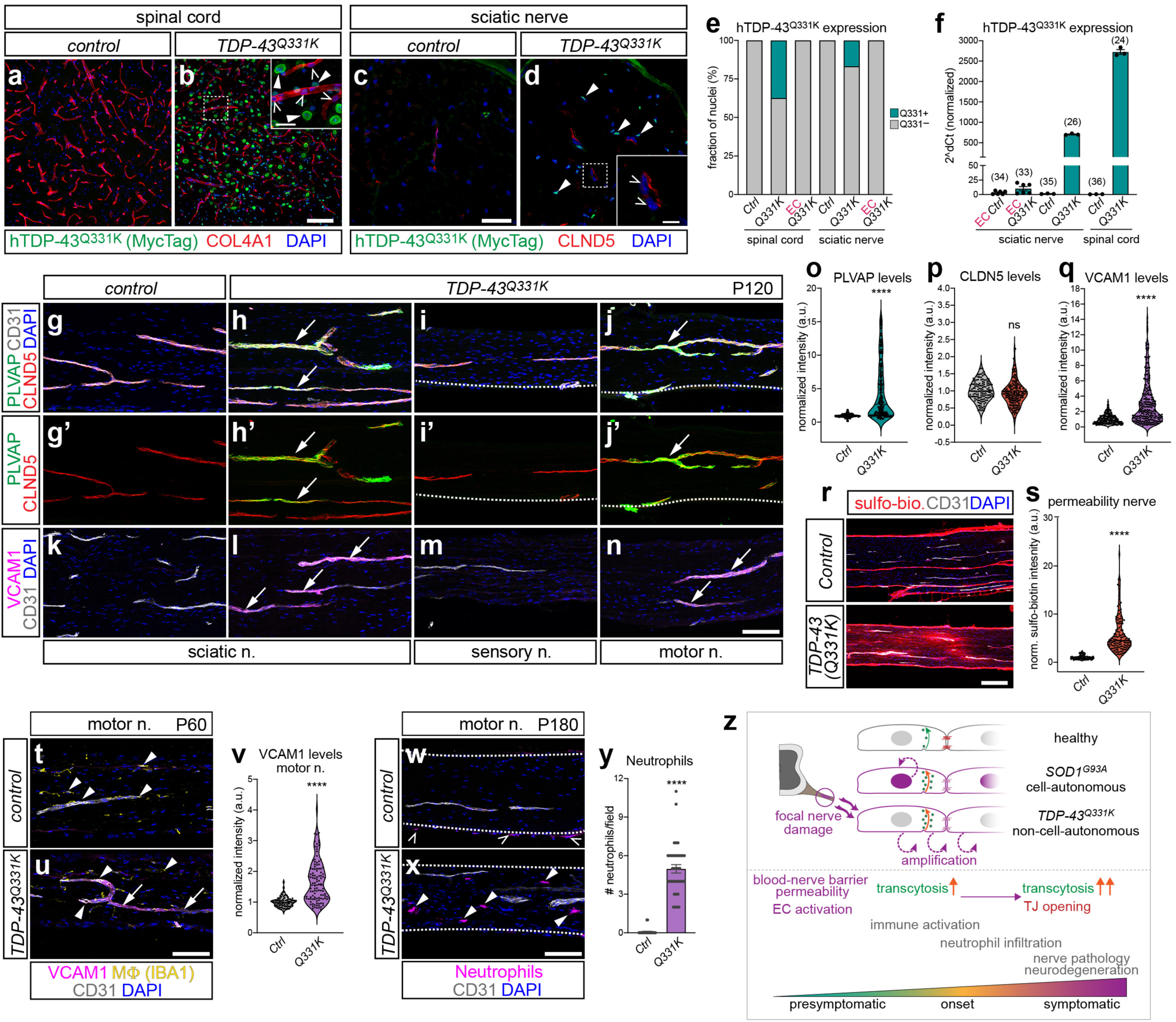
Endothelial abnormalities arise non-cell autonomously in *TDP-43^Q331K^* nerves. **a**, **b**, Spinal cords of control (**a**) and *TDP-43^Q331K^* (**b**) mice stained for Myc-Tag to reveal the TDP-43^Q331K^ transgene. TDP-43^Q331K^ is detected in nuclei of parenchymal cells (neurons and glia, closed arrowheads in inset in **b**) but not in EC nuclei (open arrowheads in inset). Myc-Tag does not label non-transgenic controls (**a**). COL4A1 marks vessels; DAPI, nuclei. **c**, **d**, Transverse sections of control (**c**) and *TDP-43^Q331K^* (**d**) sciatic nerves stained for Myc-Tag. *TDP-43^Q331K^*is not detected in EC nuclei (open arrowheads in inset in **d**). Sparse Myc-Tag^+^ nuclei are visible in the nerve parenchyma (arrowheads). CLDN5 marks endoneurial vessels; DAPI, nuclei. **e**, Percentage of Myc-tag^+^ nuclei (green, Q331K^+^) in the spinal cord and sciatic nerves of control and *TDP-43^Q331K^* mice. EC nuclei are Myc-tag^−^. **f**, *TDP-43^Q331K^*transgene expression measured by qRT-PCR in whole spinal cords and sciatic nerves of control and *TDP-43^Q331K^*mice, or in FACS-purified ECs from each tissue. Values represent 2^–ΔCt normalized to the EC-negative fraction. Average cycle threshold (Ct) are indicated above each bar. **g-j’**, Endoneurial vessels of sciatic nerves from P120 control (**g**, **g’**) and *TDP-43^Q331K^*mice (**h**, **h’**), or sensory (**i**, **i’**) and motor (**j**, **j’**) *TDP-43^Q331K^*nerves stained for PLVAP and CLDN5. PLVAP is upregulated in mutant sciatic and motor nerves (**h**, **h’**, **j**, **j’**, arrows). PLVAP and CLDN5 are shown separately in **g’-j’**. Dashed contours in **i** and **j** outline the endoneurium. CD31 marks vessels; DAPI, nuclei. **k-n**, Endoneurial vessels of sciatic, sensory, and motor nerves of controls or *TDP-43^Q331K^* mice stained for VCAM1. VCAM1 is upregulated in *TDP-43^Q331K^*sciatic and motor nerves (**l**, **n**, arrows). CD31, vessels; DAPI, nuclei. **o**, **p**, PLVAP (**o**) and CLDN5 (**p**) signal in sciatic nerve endoneurial vessels at P120, normalized to average control value. **q**, VCAM1 signal in sciatic nerve endoneurial vessels at P120, normalized to average control value. **r**, Sciatic nerves of P180 control (upper panel) and *TDP-43^Q331K^* (lower panel) mice injected with sulfo-biotin. The tracer leaks from vessels into the nerve parenchyma in mutants. CD31, vessels; DAPI, nuclei. **s**, Extravascular sulfo-biotin signal in control and *TDP-43^Q331K^* sciatic nerves, normalized to average control value. **t-u**, Endoneurial vessels of motor nerves from presymptomatic P60 *TDP-43^Q331K^* mice (**u**) upregulate VCAM1 (arrows). At this stage, IBA1^+^ macrophages (arrowheads) are comparable between control (**t**) and mutant (**u**) nerves. **v**, VCAM1 signal in motor nerve endoneurial vessels at P60, normalized to average control value. **w**, **x**, Neutrophils identified by Ly-6B.2 (7/4) staining in motor nerves of control (**w**) and *TDP-43^Q331K^*(**x**) mice. Extravascular neutrophils are present in mutants (**x**, closed arrowheads). Dashed contours outline the endoneurium. Few neutrophils are visible in the epineurium of controls (**w,** open arrowheads). **y**, Neutrophil counts in motor and sciatic nerves of control and *TDP-43^Q331K^* mice at P180. Mean ± SEM. **z**, Model: The nerve endothelium senses and amplifies focal ALS-related nerve damage, promoting neuroinflammation and neurodegeneration. EC activation and BNB disruption are driven by non-cell autonomous (*TDP-43^Q331K^*) and cell-autonomous (*SOD1^G93A^*) mechanisms. Aberrant upregulation of transcytosis during the presymptomatic phase precedes TJ opening. Scale bars: a, b, 100μm; inset in b, 25μm; c, d, 25μm; inset in d, 5μm; g-n, 100μm; r, 200μm; t, u, 100μm; w, x, 100μm

Despite this expression pattern, in sciatic and motor nerves of *TDP-43^Q331K^*mice at onset stage (P120; Fig. 1m), but not in sensory nerves, we observed the full spectrum of endothelial abnormalities identified in *SOD1^G93A^* mutants, including upregulation of PLVAP, which preceded changes in CLDN5 levels (Fig. 7g-j’, o, p) —supporting TEM findings showing early accumulation of pinocytotic vesicles in ECs before the loosening of TJs (Fig. 1l, r and Suppl. Fig. 2u-v’). EC activation, marked by VCAM1 induction in endoneurial vessels, was already significant in pre-symptomatic mutants (P60), anticipating the increase in macrophage numbers, which became conspicuous at later stages (Fig. 7k-n, q, t-v and Suppl. Fig. 11a-e). Vascular inflammation in *TDP-43^Q331K^*nerves was further evidenced by elevated expression of VWF and ICAM1 in endoneurial ECs, coinciding with PLVAP upregulation (Suppl. Fig. 11f-h). Similar to *SOD1^G93A^*mice, *TDP-43^Q331K^* mutants displayed BNB leakage of sulfo-biotin in motor and sciatic nerves, but not sensory nerves (Fig. 7r, s and Suppl. Fig. 1i-l). In contrast, BSCB permeability remained intact despite astro-microgliosis and motor neuron loss in the spinal cord of symptomatic *TDP-43^Q331K^* mice (P180) (Suppl. Fig. 11m-s). At this stage, neutrophils were found to extravasate into motor and sciatic nerves (Fig. 7w-y and data not shown). In *TDP-43^Q331K^* mice, endothelial phenotypes were penetrant and reached a severity comparable to *SOD1^G93A^*in the affected areas but were less widespread throughout the nerve. The more focal nature of this damage likely reflects the milder motor neuron pathology of *TDP-43^Q331K^*mice compared to the *SOD1^G93A^* model^68^.

Thus, despite lacking *TDP-43^Q331K^* expression, ECs in peripheral nerves developed vascular abnormalities similar to those observed in *SOD1^G93A^* mice, where the mutant transgene is present ubiquitously. To further confirm this finding, we took advantage of the design of the *TDP-43^Q331K^* construct, which is flanked by loxP sites, to excise the mutant transgene specifically from ECs via Cre-mediated recombination using *Cdh5*^*CreER*110^. *Cdh5^CreER^; TDP-43^Q331K^* mice were treated with tamoxifen at P30 and analyzed at P135 (post-onset) (Suppl. Fig. 11t-w). EC activation and BNB alterations in these mutants remained comparable to *TDP-43^Q331K^* mice (Suppl. Fig. 11x-z and data not shown), providing additional evidence that the nerve endothelium becomes dysfunctional even in the absence of a direct effect of ALS genes in ECs.

These results imply that vascular defects in peripheral nerves can arise non-cell autonomously, with ECs responding to ALS-related changes and signals derived from their tissue microenvironment. Notably, axonal degeneration was minimal at the onset of vascular abnormalities in *TDP-43^Q331K^* mice (Suppl. Fig. 2o-t), underscoring the high sensitivity of the nerve endothelium to ALS-associated pathology.

## DISCUSSION

Peripheral nerve abnormalities —including axonal injury, defects in retrograde transport, neuromuscular junction alterations, and neuroinflammation— have long been considered central features of disease progression^47,48,111–113^. In this study, we identify the BNB as both a sensor and an active facilitator of ALS progression. We demonstrate that disease-associated alterations in nerve ECs are not only central to the pathophysiology but also reversible through targeted modulation of immune cell infiltration, opening potential therapeutic avenues to delay disease progression. Here, we provide evidence that nerve ECs not only react to early peripheral pathology but also may actively propagate disease signals, thereby amplifying neuroinflammation and axonal degeneration. In addition, we delineate EC-specific molecular signatures that may serve as biomarkers of BNB dysfunction in ALS and potentially in other PNS disorders. Our findings align with the concept that ECs are sensitive to perturbations in both systemic circulation and their local tissue environment^114^, and mount rapid responses that can become maladaptive in chronic disease states^7,115^.

### The peripheral nerve vasculature is a gateway for motor neuron degeneration

Diagnostic nerve biopsies obtained from living patients —unlike postmortem specimens, which have typically been used to study ALS-related vascular changes in the brain and spinal cord— provid a unique window into the early phases of disease^51,116^. Unexpectedly, in motor nerves we observed vascular alterations that preceded overt axonal loss, supporting the early involvement of BNB dysfunction in ALS pathogenesis. The vulnerability of the PNS vasculature is supported by findings of BNB alterations in vasculitic and inflammatory neuropathies^58,59,117,118^, as well as in chemotherapy-induced peripheral neuropathy, where excessive vasoconstriction in nerve blood vessels causes hypoxia and pain^119^. In line with these observations, BNB disruption in presymptomatic ALS mice results in leakage of plasma proteins such as fibrinogen, which impairs axonal regeneration and myelination^120^ and activates innate immune pathways that exacerbate neuronal injury^121,122^. BBB and BSCB disruption in ALS has been studied mainly through histology and permeability assays, while transcriptional characterization has been lacking^14,19,20,123^. By profiling ECs from spinal cord and peripheral nerves, we observed minimal transcriptional changes in spinal cord even at symptomatic stages, when astro- and microgliosis are already evident. In contrast, peripheral nerve ECs showed robust alterations already at presymptomatic phases. Notably, nerve ECs a higher “activated” tone at baseline compared to spinal cord ECs, which may predispose them to earlier and stronger pathological responses. Under ALS conditions, this baseline priming becomes amplified, with nerve ECs showing increased expression of inflammatory mediators such as VCAM1 and ICAM1, which are broadly implicated in vascular inflammation^76,124,125^. By comparison, neither VCAM1, ICAM1, nor PLVAP —a marker linked to increased transcellular permeability— are induced in spinal cord ECs of mutant mice, highlighting the selective vulnerability of peripheral nerve vasculature. Consistent with this, gene ontology analyses highlighted enrichment of NF-κB and TNF-α signaling pathways, which are major regulators of EC activation^76,126,127^. Notably, endothelial-specific inhibition of NF-κB has been shown to reduce tissue inflammation^128^, suggesting a promising therapeutic approach for ALS-related vascular dysfunction.

### Organotypic endothelial identity underlies vulnerability of peripheral nerve vasculature in ALS

Single-cell RNA-sequencing studies have revealed extensive heterogeneity among ECs across species and disease states^3,77,129^. We found that in ALS peripheral nerves, capillary endothelial subtypes are the primary responders to disease: venous capillaries show pronounced upregulation of genes linked to inflammation, activation, and increased permeability, whereas barrier capillaries largely downregulate genes involved in transport and vascular homeostasis. Venous capillaries are known to be susceptible in CNS vascular pathologies. For instance, they are the site of origin for cerebral cavernous malformations^130,131^ and show the strongest inflammatory and neoangiogenic response and contribute to in EAE^94,132^. We identified equivalent populations of capillary venous ECs in both nerve and spinal cord based on shared markers; however, the nerve subset exhibited a distinctive baseline profile, marked by elevated expression of pro-inflammatory and activation-associated genes. In ALS, these nerve venous ECs further upregulated a set of genes (e.g., *Plvap*, *Vwf*, *Aqp1*, *Tacr1, Aplnr*) not induced in the corresponding spinal cord subset during EAE inflammation^94^, revealing a unique, nerve-specific transcriptional response to ALS-related damage. Venous markers VWF and VCAM1, which are elevated in aged CNS vasculature and implicated in EC activation, atherosclerosis, and stroke^114,133–135^, increased in the pre-symptomatic phase exclusively in motor and sciatic nerve ECs of both *SOD1* and *TARDBP* mutant mice, but remained unchanged in the spinal cord vasculature. A similar pattern was observed for TACR1 (NK1R), which mediates inflammatory signaling and cytokine production^136^, and for LRG1, an IL6-induced secreted glycoprotein involved in cardiovascular disease and innate immunity that promotes neutrophil chemotaxis and fibrosis^137^. Collectively, these findings highlight the activated state of the nerve endothelium consistent with the organotypic features of the capillary–venous EC subset and its selective vulnerability in ALS.

### Enhanced transcellular permeability underlies early BNB disruption

The two cardinal mechanisms of barrier maintenance —transcellular and paracellular transport— appear differentially sensitive to pathological stress. Our data indicate that enhanced transcellular route, rather than TJ disruption, is the primary driver of BNB breakdown in presymptomatic ALS, with structural alterations in EC contacts becoming evident only at more advanced stages. In pre-symptomatic ALS models, focal PLVAP activation in nerve vasculature marks early vascular insult. This is accompanied by selective downregulation of MFSD2A, a key inhibitor of caveolae-mediated transcytosis, and increased cytoplasmic localization of CAV1, consistent with electron microscopy evidence of elevated vesicular transport. Collectively, these changes confirm a pathological surge in transcytosis preceding junctional breakdown. Similar dynamics are observed in stroke models, where transcellular leakage precedes TJ disruption^138^. Notably, PLVAP+ vessels still retain tight junction markers such as CLDN5, and we observe ECs in a hybrid PLVAP^−^/CLDN5^+^ state, suggesting a partial transition toward a non-barrier phenotype rather than complete transcriptional reprogramming. This aligns with evidence of a reduction in Wnt tone, a key driver of barrier induction and maintenance^72,139,140^, despite the pathway remain active —as evidenced by LEF1 expression in ALS ECs (data not shown)— indicating that ECs retain core barrier identity while selectively deregulating specific molecular components that perturb BNB integrity. Together, these observations establish a mechanistic framework in which early, graded dysregulation of transcellular permeability underlies selective BNB vulnerability in ALS. This early response of the transcytotic pathway provides a potential window for therapeutic intervention. Temporary modulation of transcellular permeability could facilitate targeted delivery of neuroprotective factors^60^, and strategies such as overexpression of the transcytosis suppressor MFSD2A or inhibition of vesicular transport have demonstrated efficacy in rescuing BBB alterations in brain disorders^141–143^.

### Endothelial–immune crosstalk drives peripheral nerve inflammation in ALS

Endothelial–immune interactions actively shape tissue inflammation and disease progression in neurological disorders, including ALS^144,145^. In ALS nerves, endothelial activation drives early upregulation of adhesion molecules and inflammatory mediators, facilitating immune cell recruitment. Recruited immune cells then feed-back on ECs, amplifying barrier dysfunction and sustaining inflammation ^106,146^. High-dimensional flow cytometry revealed expansion of neutrophils, inflammatory monocytes, and activated macrophages in the peripheral nerves of *SOD1^G93A^*mice, paralleling EC transcriptional shifts toward a pro-inflammatory state. Resident macrophages, essential for maintaining vascular homeostasis in the PNS^60^, acquire an activated phenotype in ALS, consistent with previous reports^53,54,57^. Our analysis revealed that resident macrophages, normally perivascular in healthy nerve, transition from a homeostatic to an activated state in situ, without recruitment of circulating monocytes. Activated macrophages detach from vessels, become phagocytic, and undergo an inflammatory switch, accompanied by relocalization within the endoneurium. This process may be driven by downregulation of CX3CL1 on endoneurial ECs, a ligand for CX3CR1 expressed by resident macrophages, and a genetic modifier in ALS^147^. Nerve EC activation was accompanied by substantial infiltration of neutrophils that modulate the activation state of macrophage populations. Neutrophils localize near degenerating axons and vessels, amplifying vascular dysfunction and inflammation^106^, and compromising barrier integrity via protease and ROS release and downregulation of TJ proteins such as CLDN5 ^148^. Neutrophil depletion restored BNB integrity and alleviated axons degeneration, without however improving motor symptoms. These results highlights neutrophils as key modulators of ALS pathology in peripheral nerves and suggests that therapeutic strategies targeting neutrophils, alone or combined with interventions to normalize EC function, may offer a promising avenue^55,149^.

### Non–cell-autonomous ALS signals trigger endothelial activation

Mutant TDP-43 can cell-autonomously regulate angiogenesis and barrier function, and its overexpression in brain ECs induces neuroinflammation and BBB alterations^37–39,150^. However, pTDP-43 aggregates in the spinal cord do not consistently correlate with BSCB leakage^32^, leaving it unclear whether vascular dysfunction arises from direct effects in ECs or indirectly via signals from neurons, glia, or other surrounding cells.

In our study, EC activation in the *TDP-43^Q331K^* model occurs in motor —but not sensory— nerves prior to detectable axonal degeneration, even in the absence of mutant protein in ECs. These observations indicate that ECs respond non–cell autonomously to ALS-associated signals, likely originating from vulnerable motor axons or the surrounding microenvironment in a pre-degenerative state. In *SOD1^G93A^* mice, where all cells express the mutant protein, vascular pathology is quantitatively more severe although all hallmarks of vascular dysfunction are shared across these two models, suggesting that both cell-autonomous and non–cell autonomous mechanisms contribute to peripheral EC activation and BNB disruption in ALS. Together, our data indicate that peripheral nerve ECs act as sensors of early ALS-related damage, integrating pathological cues from the tissue microenvironment and mounting maladaptive responses. Elucidating the nature of these activating signals and the relative contributions of intrinsic and extrinsic mechanisms to endothelial dysfunction in the PNS remains an important open question, with direct relevance for defining pathological pathways and identifying potential windows for therapeutic intervention.

## Acknowledgements

We thank Ralf Adams for *Cdh5(PAC)-CreERT2* mice; Cesare Covino and the Advanced Microscopy Laboratory (ALEMBIC) of San Raffaele Hospital for expertise and instrumentation. Funding sources: AriSLA Foundation pilot grant (NeVALS), Italian Ministry of Health GR-2016-02364327, Giovanni Armenise-Harvard Foundation Career Development Award.

## Author contributions

G.P.B. designed experiments, collected and analyzed data with the help of A.B., A.M., I.B., and L.M. S.D. performed computational analysis with the help of SdP. C.B. performed high-dimensional flow cytometry and data analysis. P.P. conducted nerve morphology and electron microscopy studies. D.Bos. performed cell sorting. G.N. and C.B. provided nerve samples from mutant mice. N.R. selected and analyzed clinical cases. M.I. provided critical comments for immunological studies. A.Q. collected human nerve samples with the help of N.R, and analyzed morphological data. D.B. supervised the study, analyzed data and wrote the manuscript with G.P.B.

## MATERIALS AND METHODS

### Sample size and Statistical analysis

Statistical significance is reported as follows: ns, not significant (p >0.05); * p <0.05; ** p <0.01; *** p <0.001; **** p <0.0001.

**Fig. 1j:** N, TEM images (≥3 from each motor nerve biopsy). One-way ANOVA with Dunnett’s test, p<0.001 Mild (includes 2 ALS “normal” and 4 “mild” cases) vs. Control (3 non-ALS neuropathies/myopathy cases); p<0.0001 Severe (includes 2 ALS “moderate” and 4 “severe” cases) vs. Control.

**Fig. 1k:** N, TEM images from motor nerves, control (8 nerves), *SOD1^G93A^* P60 (4), P90 (6), P120 (8). One-way ANOVA with Dunnett’s test, (ns) p=0.74 *SOD1^G93A^* P60 vs. control; p<0.0001 *SOD1^G93A^* P60 and P120 vs. control.

**Fig. 1l:** N, TEM images from motor nerves, control (9 nerves P60-P180), *TDP-43^Q331K^* P60 (2), P120 (4), P180 (48). One-way ANOVA with Dunnett’s test, p<0.0001 *TDP-43^Q331K^* P60, P120, and P180 vs. control.

**Suppl. Fig. 2e, f:** N, sections/mice: control (9/7 mice at P60-P120), *SOD1^G93A^* P60 (9/4), P90 (11/5), P120-150 (6/4). One-way ANOVA with Dunnett’s test. Total fiber counts (**e**), (ns) p=0.24 *SOD1^G93A^*P60 vs. control; p<0.05 *SOD1^G93A^* P90 vs. control; p<0.001 *SOD1^G93A^* P120-150 vs. control. Degenerating fibers/total (**f**), (ns) p=0.66 *SOD1^G93A^* P60 vs. control; p<0.0001 *SOD1^G93A^* P90 and P120-150 vs. control.

**Suppl. Fig. 2g, h:** N, mice: 16-26 per genotype at each stage. One-way ANOVA with Sidak’s test. Rotarod test (**g**), (ns) p=0.94 *SOD1^G93A^* vs. control at P60; (ns) p=0.16 *SOD1^G93A^*vs. control at P90; p<0.0001 *SOD1^G93A^* vs. control at P120. BDW measurement (**h**), (ns) p=0.42 *SOD1^G93A^* vs. control at P60; (ns) p=0.43 *SOD1^G93A^* vs. control at P90; p<0.0001 *SOD1^G93A^* vs. control at P120.

**Suppl. Fig. 2s, t:** N, sections/mice: control (13/6 mice at P60-P120), *TDP-43^Q331K^* P60 (16/6), P120 (6/4), P180 (6/6). One-way ANOVA with Dunnett’s test. Total fiber counts (**s**), (ns) p=0.79 *TDP-43^Q331K^*P60 vs. control; (ns) p=0.45 *TDP-43^Q331K^* P120 vs. control; p<0.01 *TDP-43^Q331K^*P180 vs. control. Degenerating fibers/total (**t**), p<0.01 *TDP-43^Q331K^* P60 vs. control, p<0.001 *TDP-43^Q331K^*P120 and P180 vs. control.

**Fig. 2f:** N, tiled regions of interest (ROIs) spanning the sciatic nerve parenchyma from 5-6 mice per genotype/stage. Control values are pooled from P60-P120. One-way ANOVA with Dunnett’s test, p<0.05 *SOD1^G93A^* P60 vs. control; p<0.0001 *SOD1^G93A^*P90 and P120 vs. control.

**Fig. 2g:** N, tiled ROIs spanning the spinal cord (4 per spinal cord) from 5-6 mice per genotype/stage. Control values are pooled from P60-P120. One-way ANOVA with Dunnett’s test, (ns) p=0.98 *SOD1^G93A^* P60 vs. control; (ns) p=0.74 *SOD1^G93A^* P90 vs. control; p<0.0001 *SOD1^G93A^*P120 vs. control.

**Suppl. Fig. 3o:** N, spinal cord images (∼10 per spinal cord) from 5-6 mice per genotype/stage. Control values are pooled from P60-P150. One-way ANOVA with Dunnett’s test, (ns) p=0.67 *SOD1^G93A^* P60 vs. control; (ns) p=0.22 *SOD1^G93A^* P90 vs. control; p<0.0001 *SOD1^G93A^*P150 vs. control.

**Suppl. Fig. 3p, q:** N, tiled ROIs spanning spinal cords from 6 control mice (P60-P150), 2 *SOD1^G93A^* P60, 2 *SOD1^G93A^* P90, and 3 *SOD1^G93A^* P150. One-way ANOVA with Dunnett’s test. GFAP staining (**p**), (ns) p=0.58 *SOD1^G93A^*P60 vs. control; p<0.0001 *SOD1^G93A^* P90 and P120 vs. control. IBA1 staining (**q**), (ns) p=0.74 *SOD1^G93A^* P60 vs. control; p<0.01 *SOD1^G93A^* P90 vs. control; p<0.0001 *SOD1^G93A^* P150 vs. control.

**Suppl. Fig. 3v:** N, spinal cord vessels from 5-8 mice per genotype/stage. Control values are pooled from P60-P120. One-way ANOVA with Dunnett’s test, (ns) p=0.89 *SOD1^G93A^* P60 vs. control; p<0.01 *SOD1^G93A^* P90 vs. control; p<0.0001 *SOD1^G93A^*P120 vs. control.

**Suppl. Fig. 3w:** N, sciatic nerve vessels from 7-12 mice per genotype/stage. Control values are pooled from P60-P120. One-way ANOVA with Dunnett’s test, (ns) p=0.12 *SOD1^G93A^* P60 vs. control; (ns) p=0.82 *SOD1^G93A^* P90 vs. control; (ns) p=0.43 *SOD1^G93A^* P120 vs. control.

**Fig. 3j, k:** N, vessels from tiled ROIs spanning sciatic nerves from 5-6 mice per genotype/stage. Control values are pooled from P60-P120. One-way ANOVA with Dunnett’s test. PLVAP staining (**j**), p<0.0001 *SOD1^G93A^* P60, P90 and P120 vs. control. CLDN5 staining (**k**): (ns) p=0.72 *SOD1^G93A^* P60 vs. control; (ns) p=0.079 *SOD1^G93A^* P90 vs. control; p<0.0001 *SOD1^G93A^* P120 vs. control.

**Fig. 3l, m:** N, spinal cord vessels from 8 control and 9 *SOD1^G93A^* mice. Unpaired t test. PLVAP staining (**l**), (ns) p=0.56 *SOD1^G93A^* vs. control. CLDN5 staining (**m**): p<0.0001 *SOD1^G93A^* vs. control.

**Fig. 3p:** N, tiled regions ROIs spanning the sciatic nerve parenchyma from 4 control and 5 *SOD1^G93A^* mice. Unpaired t test, p<0.0001 *SOD1^G93A^* vs. control.

**Fig. 3s:** N, tiled regions ROIs spanning the nerve parenchyma from 5 control and 6 *SOD1^G93A^* mice. Pooled measurements from sciatic and motor nerves. Unpaired t test, p<0.0001 *SOD1^G93A^*vs. control.

**Fig. 3w, x:** N, vessels from tiled ROIs spanning sciatic nerves from 3 control and 4 *SOD1^G93A^*mice. Unpaired t test, p<0.0001 *SOD1^G93A^* vs. control.

**Fig. 3y:** N, nerve biopsies from control (2) and ALS (3) cases. p<0.01 ALS vs. control.

**Suppl. Fig. 4c:** N, vessels from tiled ROIs spanning sciatic nerves from 6 mice per genotype/stage. Unpaired t test, p<0.0001 *SOD1^G93A^*vs. control.

**Suppl. Fig. 4f:** N, vessels from tiled ROIs spanning sciatic nerves from 4 mice per genotype/stage. Unpaired t test, (ns) p=0.22 *SOD1^G93A^*vs. control.

**Suppl. Fig. 4i:** N, vessels from tiled ROIs spanning sciatic nerves from 4-5 mice per genotype/stage. Control values are pooled from P60-P190. One-way ANOVA with Dunnett’s test, (ns) p>0.99 *SOD1^G93A^* P60 vs. control; (ns) p=0.73 *SOD1^G93A^* P90 vs. control; p<0.0001 *SOD1^G93A^*P140 vs. control.

**Suppl. Fig. 4l:** N, ROIs spanning the sciatic nerve parenchyma from 4-5 mice per genotype/stage. One-way ANOVA with Dunnett’s test, (ns) p=0.07 P60 vs. control; p<0.0001 *SOD1^G93A^*P90 and P140 vs. control.

**Suppl. Fig. 4o:** N, tiled regions ROIs spanning the sciatic nerve parenchyma from 3 mice per genotype. Unpaired t test, p<0.0001 *SOD1^G93A^* vs. control.

**Suppl. Fig. 4r:** N, spinal cord vessels from 3 mice per genotype. Unpaired t test, (ns) p=0.06 *SOD1^G93A^* vs. control.

**Suppl. Fig. 4v:** N, vessels (18-26) from 4-5 mice per genotype/stage.

**Suppl. Fig. 4y:** N, vessels (23) from 2 wild-type mice.

**Fig. 4s, t:** N, CLDN5+ vessels (>100) from 4-5 mice per genotype/stage. Controls are pooled from different stages.

**Suppl. Fig. 5g, h:** N, tiled regions ROIs spanning the sciatic nerve parenchyma from 4-5 mice per genotype/stage. One-way ANOVA with Sidak’s test. (ns) p>0.9 CAR4^−^ vs. CAR4^+^ control; p<0.0001 CAR4^−^ vs. CAR4^+^ *SOD1^G93A^*; p<0.001 CAR4^−^ vs. CAR4^+^ *TDP-43^Q331K^*.

**Fig. 5a, b:** N, vessels from tiled ROIs spanning sciatic nerves from 5 control and 4 *SOD1^G93A^* mice at P60, and 4 control and 6 *SOD1^G93A^*mice at P120. One-way ANOVA with Sidak’s test, p<0.0001 *SOD1^G93A^* vs. control.

**Fig. 5g:** N, vessels from tiled ROIs spanning sciatic nerves from 7-8 mice per genotype/stage. Control values are pooled from different stages. One-way ANOVA with Dunnett’s test, p=0.0035 *SOD1^G93A^* P60 vs. control; p<0.0001 *SOD1^G93A^* P90 and P120 vs. control.

**Fig. 5l:** N, reporter-labeled cells (MF) counted in 55-80 fields of view spanning sciatic nerves from 3-4 mice per genotype/stage. Control values are from P60. Statistical annotations are color-coded by cell labeling. Mixed-effect analysis with Tukey’s test, *SOD1^G93A^*P60 vs. control; *SOD1^G93A^* P90 vs. P60; *SOD1^G93A^*P120 vs. P90.

**Fig. 5m:** N, 3 independent sciatic nerve preparations per genotype, each containing nerves pooled from 2–4 mice. Unpaired t test, p<0.01 *SOD1^G93A^* vs. control.

**Fig. 5o:** N, 4 independent sciatic nerve preparations per genotype, each containing nerves pooled from 2–4 mice. Unpaired t test, (ns) p=0.14 DC *SOD1^G93A^* vs. control; p<0.0001 MF3 *SOD1^G93A^* vs. control; p<0.01 other cell clusters *SOD1^G93A^* vs. control.

**Fig. 5q:** N, sciatic nerves from 6 mice per genotype. Data pooled from 3 independent experiments. Unpaired t test, p<0.0001 rMF *SOD1^G93A^* vs. control; p<0. 01 aMF *SOD1^G93A^*vs. control; p<0.001 monocytes *SOD1^G93A^* vs. control.

**Fig. 5s:** N, sciatic nerves from 4 mice per genotype. One-way ANOVA with Tukey’s test for indicated pairwise comparisons.

**Fig. 5z:** N, vessels from tiled ROIs spanning sciatic nerves from 3-4 mice per genotype. One-way ANOVA with Sidak’s test, (ns) p=0.8 PLVAP *SOD1^G93A^; Ccr2^-/-^* vs. *SOD1^G93A^*; (ns) p=0.31 VCAM1 *SOD1^G93A^; Ccr2^-/-^* vs. *SOD1^G93A^*.

**Suppl. Fig. 7h:** N, spinal cord vessels from 3 mice per genotype. Unpaired t test, (ns) p=0.32 *SOD1^G93A^* vs. control.

**Suppl. Fig. 7k:** N, spinal cord vessels from 3 control and 4 *SOD1^G93A^* mice. Unpaired t test, (ns) p=0.52 *SOD1^G93A^* vs. control.

**Suppl. Fig. 7q:** N, vessels from tiled ROIs spanning sciatic nerves (6 control, 7 *SOD1^G93A^*). Unpaired t test, p<0.0001 *SOD1^G93A^* vs. control.

**Suppl. Fig. 7t:** N, reporter-labeled cells (MF) counted in 22-36 fields of view spanning motor nerves (3 control, 4 *SOD1^G93A^*). Statistical annotations are color-coded by cell labeling. Mixed-effect analysis with Sidak’s test, *SOD1^G93A^* vs. control.

**Suppl. Fig. 8c, e:** N, sciatic nerves from 5 mice per genotype. Unpaired t test, *SOD1^G93A^* vs. control.

**Suppl. Fig. 8j:** N, fields of view for MF counts, spanning sciatic nerves from 5 control, 5 *SOD1^G93A^*, and 6 *SOD1^G93A^; Ccr2^-/-^* mice. One-way ANOVA with Sidak’s test, p<0.01 *SOD1^G93A^; Ccr2^-/-^* vs. *SOD1^G93A^*.

**Fig. 6b:** N, sciatic nerves from 11 control and 7 *SOD1^G93A^* mice. Data pooled from 3 independent experiments. Unpaired t test, p<0.01 *SOD1^G93A^* vs. control.

**Fig. 6d:** N, Ly-6B.2 (7/4)-FITC^+^ neutrophils counted in 35-53 fields of view spanning motor nerves from 4-6 mice per genotype/stage. One-way ANOVA with Dunnett’s test, (ns) p=0.103 *SOD1^G93A^* P60 vs. control; p<0.0001 *SOD1^G93A^* P90 and P120 vs. control.

**Fig. 6o:** N, Ly-6B.2 (7/4)-FITC^+^ (or Ly6G^+^) neutrophils counted in 19-31 fields of view spanning sciatic nerves from 8 control and 11 *SOD1^G93A^*/isotype or anti-Ly6G-treated mice. One-way ANOVA with Tukey’s test, p<0.0001 *SOD1^G93A^*/isotype-treated vs. control; p=0.07 *SOD1^G93A^*/anti-Ly6G-treated vs. control; p<0.0001 *SOD1^G93A^*/anti-Ly6G-treated vs. *SOD1^G93A^*/isotype-treated.

**Fig. 6p:** N, vessels from tiled ROIs spanning sciatic nerves from 8 (control) and 11 (*SOD1^G93A^*/isotype or anti-Ly6G-treated) mice. One-way ANOVA with Tukey’s test, p<0.0001 all pairwise comparisons.

**Fig. 6q:** N, fields of view for MF counts, spanning sciatic nerves from 8 control and 11 *SOD1^G93A^*/isotype or anti-Ly6G-treated mice. One-way ANOVA with Tukey’s test, p<0.0001 all pairwise comparisons.

**Fig. 6r:** N, vessels from tiled ROIs spanning sciatic nerves from 8 control and 11 *SOD1^G93A^*/isotype or anti-Ly6G-treated mice. One-way ANOVA with Tukey’s test, p<0.0001 *SOD1^G93A^*/isotype-treated vs. control; (ns) p=0.0504 *SOD1^G93A^*/anti-Ly6G-treated vs. control; p<0.0001 *SOD1^G93A^*/anti-Ly6G-treated vs. *SOD1^G93A^*/isotype-treated.

**Fig. 6s:** N, vessels from tiled ROIs spanning sciatic nerves from 8 control and 11 *SOD1^G93A^*/isotype or anti-Ly6G-treated mice. One-way ANOVA with Tukey’s test, p<0.0001 *SOD1^G93A^*/isotype-treated vs. control; p<0.0001 *SOD1^G93A^*/anti-Ly6G-treated vs. control; p<0. 001 *SOD1^G93A^*/anti-Ly6G-treated vs. *SOD1^G93A^*/isotype-treated.

**Fig. 6t:** N, TEM images from sciatic nerves of 9 control, 13 *SOD1^G93A^*/isotype treated, and 17 *SOD1^G93A^*/anti-Ly6G-treated) mice. One-way ANOVA with Tukey’s test, p<0.0001 *SOD1^G93A^*/isotype-treated vs. control; (ns) p=0.367 *SOD1^G93A^*/anti-Ly6G-treated vs. control; p<0.0001 *SOD1^G93A^*/anti-Ly6G-treated vs. *SOD1^G93A^*/isotype-treated.

**Fig. 6v:** N, sciatic nerves from 6 mice per genotype. One-way ANOVA with Tukey’s test. <u>Neutrophils</u>, p=0.013 *SOD1^G93A^*/isotype-treated vs. control; p=0.20 *SOD1^G93A^*/anti-Ly6G-treated vs. control; p<0.001 *SOD1^G93A^*/anti-Ly6G-treated vs. *SOD1^G93A^*/isotype-treated; <u>Monocytes</u>, p=0.01 *SOD1^G93A^*/isotype-treated vs. control; p<0.001 *SOD1^G93A^*/anti-Ly6G-treated vs. control; (ns) p=0.25 *SOD1^G93A^*/anti-Ly6G-treated vs. *SOD1^G93A^*/isotype-treated; <u>MF1</u>, p<0.0001 *SOD1^G93A^*/isotype-treated vs. control; p<0.05 *SOD1^G93A^*/anti-Ly6G-treated vs. control; p<0.05 *SOD1^G93A^*/anti-Ly6G-treated vs. *SOD1^G93A^*/isotype-treated; <u>MF3</u>, p<0.0001 *SOD1^G93A^*/isotype-treated vs. control; p<0.001 *SOD1^G93A^*/anti-Ly6G-treated vs. control; p<0.001 *SOD1^G93A^*/anti-Ly6G-treated vs. *SOD1^G93A^*/isotype-treated.

**Fig. 6x, y:** N, motor nerves from 9 control, 13 *SOD1^G93A^*/isotype treated, and 17 or *SOD1^G93A^*/anti-Ly6G-treatedmice. One-way ANOVA with Tukey’s test. Total fiber counts (**x**), p<0.001 *SOD1^G93A^*/isotype-treated vs. control; p<0.05 *SOD1^G93A^*/anti-Ly6G-treated vs. control; (ns) p=0.32 *SOD1^G93A^*/anti-Ly6G-treated vs. *SOD1^G93A^*/isotype-treated. Degenerating fibers/total (**y**), p<0.0001 *SOD1^G93A^*/isotype-treated vs. control; p<0.001 *SOD1^G93A^*/anti-Ly6G-treated vs. control; p=0.0014 *SOD1^G93A^*/anti-Ly6G-treated vs. *SOD1^G93A^*/isotype-treated.

**Suppl. Fig. 9c:** N, Ly-6B.2 (7/4)-FITC^+^ neutrophils counted in 35-46 fields of view of spinal cords from 4 control and 6 *SOD1^G93A^* mice. Unpaired t test, (ns) p=0.302 *SOD1^G93A^*vs. control.

**Suppl. Fig. 9f:** N, sciatic nerves from 6 control, 7 *SOD1^G93A^*/isotype treated, and 8 *SOD1^G93A^*/anti-Ly6G-treated mice. Data pooled from 2 independent experiments. One-way ANOVA with Tukey’s test, p<0.01 *SOD1^G93A^*/isotype-treated vs. control; (ns) p=0.66 *SOD1^G93A^*/anti-Ly6G-treated vs. control; p<0.001 *SOD1^G93A^*/anti-Ly6G-treated vs. *SOD1^G93A^*/isotype-treated.

**Suppl. Fig. 9m:** N, Ly-6B.2 (7/4)-FITC^+^ (or Ly6G^+^) neutrophils counted in 13-21 fields of view spanning motor nerves from 8 control and 11 *SOD1^G93A^*/isotype or anti-Ly6G-treated mice. One-way ANOVA with Tukey’s test, p<0.0001 *SOD1^G93A^*/isotype-treated vs. control; p<0.05 *SOD1^G93A^*/anti-Ly6G-treated vs. control; p<0.0001 *SOD1^G93A^*/anti-Ly6G-treated vs. *SOD1^G93A^*/isotype-treated.

**Suppl. Fig. 9n:** N, vessels from tiled ROIs spanning sciatic nerves from 8 control and 11 *SOD1^G93A^*/isotype or anti-Ly6G-treated mice. One-way ANOVA with Tukey’s test, p<0.0001 *SOD1^G93A^*/isotype-treated vs. control; p<0.05 *SOD1^G93A^*/anti-Ly6G-treated vs. control; p<0.01 *SOD1^G93A^*/anti-Ly6G-treated vs. *SOD1^G93A^*/isotype-treated.

**Suppl. Fig. 9o:** N, vessels from tiled ROIs spanning sciatic nerves from 8 control and 11 *SOD1^G93A^*/isotype or anti-Ly6G-treated mice. One-way ANOVA with Tukey’s test, p<0.0001 *SOD1^G93A^*/isotype-treated vs. control; p<0.0001 *SOD1^G93A^*/anti-Ly6G-treated vs. control; (ns) p=0.43 *SOD1^G93A^*/anti-Ly6G-treated vs. *SOD1^G93A^*/isotype-treated.

**Suppl. Fig. 9s:** N, spinal cord images from 11 control, 11 *SOD1^G93A^*/isotype-treated, and 13 *SOD1^G93A^*/anti-Ly6G-treated) mice. One-way ANOVA with Tukey’s test, p<0.0001 *SOD1^G93A^*/isotype-treated vs. control; p<0.0001 *SOD1^G93A^*/anti-Ly6G-treated vs. control; p<0.05 *SOD1^G93A^*/anti-Ly6G-treated vs. *SOD1^G93A^*/isotype-treated.

**Suppl. Fig. 9t, u:** N, tiled ROIs spanning spinal cords from 6 control, 7 *SOD1^G93A^*/isotype treated, and 8 *SOD1^G93A^*/anti-Ly6G-treated mice. One-way ANOVA with Tukey’s test. GFAP staining (**t**), p<0.0001 *SOD1^G93A^*/isotype-treated vs. control; p<0.0001 *SOD1^G93A^*/anti-Ly6G-treated vs. control; (ns) p=0.209 *SOD1^G93A^*/anti-Ly6G-treated vs. *SOD1^G93A^*/isotype-treated. IBA1 staining (**u**), p<0.0001 *SOD1^G93A^*/isotype-treated vs. control; p<0.0001 *SOD1^G93A^*/anti-Ly6G-treated vs. control; (ns) p=0.82 *SOD1^G93A^*/anti-Ly6G-treated vs. *SOD1^G93A^*/isotype-treated.

**Suppl. Fig. 9w:** N, mice; 16 control, 17 *SOD1^G93A^*/isotype treated, and 18 *SOD1^G93A^*/anti-Ly6G-treated mice. One-way ANOVA with Tukey’s test, p<0.0001 *SOD1^G93A^*/isotype-treated vs. control; p<0.0001 *SOD1^G93A^*/anti-Ly6G-treated vs. control; (ns) p=0.98 *SOD1^G93A^*/anti-Ly6G-treated vs. *SOD1^G93A^*/isotype-treated.

**Suppl. Fig. 9w:** N, mice; 16 control, 17 *SOD1^G93A^*/isotype treated, and 18 *SOD1^G93A^*/anti-Ly6G-treated mice. One-way ANOVA with Tukey’s test, p<0.0001 *SOD1^G93A^*/isotype-treated vs. control; p<0.0001 *SOD1^G93A^*/anti-Ly6G-treated vs. control; (ns) p>0.99 *SOD1^G93A^*/anti-Ly6G-treated vs. *SOD1^G93A^*/isotype-treated.

**Suppl. Fig. 9x:** N, mice; 18 control, 15 *SOD1^G93A^*/isotype treated, and 18 *SOD1^G93A^*/anti-Ly6G-treated mice. Unpaired t test, p<0.01 *SOD1^G93A^*/anti-Ly6G-treated vs. control at 16 weeks; p<0.05 p<0.0001 *SOD1^G93A^*/isotype-treated vs. control at 16 weeks. (ns) p>0.05 all other pairwise comparisons.

**Suppl. Fig. 10e:** N, sciatic nerves from 7 control, 6 *SOD1^G93A^*/isotype treated, and 6 *SOD1^G93A^*/anti-Ly6G-treated mice. Data pooled from 2 independent experiments One-way ANOVA with Tukey’s test. <u>rMF</u>, p<0.0001 *SOD1^G93A^*/isotype-treated vs. control; p<0.05 *SOD1^G93A^*/anti-Ly6G-treated vs. control; p<0.05 *SOD1^G93A^*/anti-Ly6G-treated vs. *SOD1^G93A^*/isotype-treated. <u>aMF</u>, p<0.05 *SOD1^G93A^*/isotype-treated vs. control; (ns) p=0.89 *SOD1^G93A^*/anti-Ly6G-treated vs. control; p<0.05 *SOD1^G93A^*/anti-Ly6G-treated vs. *SOD1^G93A^*/isotype-treated. <u>Mon</u>, p<0.001 *SOD1^G93A^*/isotype-treated vs. control; p<0.01 *SOD1^G93A^*/anti-Ly6G-treated vs. control; (ns) p=0.55 *SOD1^G93A^*/anti-Ly6G-treated vs. *SOD1^G93A^*/isotype-treated.

**Fig. 10h:** N, sciatic nerves from 6 mice per genotype. One-way ANOVA with Tukey’s test. <u>MF2</u>, p<0.01 *SOD1^G93A^*/isotype-treated vs. control; p<0.001 *SOD1^G93A^*/anti-Ly6G-treated vs. control; (ns) p=0.17 *SOD1^G93A^*/anti-Ly6G-treated vs. *SOD1^G93A^*/isotype-treated; <u>MF4</u>, p<0.0001 *SOD1^G93A^*/isotype-treated vs. control; p<0.001 *SOD1^G93A^*/anti-Ly6G-treated vs. control; (ns) p=0.39 *SOD1^G93A^*/anti-Ly6G-treated vs. *SOD1^G93A^*/isotype-treated; <u>MF5</u>, (ns) p=0.11 *SOD1^G93A^*/isotype-treated vs. control; (ns) p=0.94 *SOD1^G93A^*/anti-Ly6G-treated vs. control; (ns) p=0.066 *SOD1^G93A^*/anti-Ly6G-treated vs. *SOD1^G93A^*/isotype-treated; <u>MF6</u>, p<0.01 *SOD1^G93A^*/isotype-treated vs. control; (ns) p=0.12 *SOD1^G93A^*/anti-Ly6G-treated vs. control; (ns) p=0.35 *SOD1^G93A^*/anti-Ly6G-treated vs. *SOD1^G93A^*/isotype-treated.

**Fig. 10l:** N, Ly-6B.2 (7/4)-FITC^+^ neutrophils counted in 19-29 fields of view spanning sciatic nerves from 4 control, 6 *SOD1^G93A^*, and 6 *SOD1^G93A^; Ccr2^-/-^*mice. One-way ANOVA with Tukey’s test, p<0.0001 *SOD1^G93A^* vs. control; p<0.0001 *SOD1^G93A^; Ccr2^-/-^* vs. control; (ns) p=0.87 *SOD1^G93A^; Ccr2^-/-^* vs. *SOD1^G93A^*.

**Fig. 10n:** N, sciatic nerves from 4 mice per genotype. One-way ANOVA with Tukey’s test, p<0.0001 *SOD1^G93A^* vs. control; p<0.0001 *SOD1^G93A^; Ccr2^-/-^* vs. control; (ns) p=0.53 *SOD1^G93A^; Ccr2^-/-^* vs. *SOD1^G93A^*.

**Fig. 7e:** >500 total nuclei counted in sciatic nerve or spinal cord sections from 3 mice per genotype. All EC nuclei observed in the spinal cords (n=122) or sciatic nerves (n=41) of *TDP-43^Q331K^* mice were Myc-Tag^−^.

**Fig. 7f:** N, technical replicates from 2 independent experiments.

**Fig. 7o, p, q:** N, vessels from tiled ROIs spanning sciatic nerves from 10 mice per genotype. Unpaired t test. (**o**) p<0.0001 (**p**) (ns) p=0.0517 (**q**) p<0.0001 *TDP-43^Q331K^* vs. control.

**Fig. 7s:** N, tiled ROIs spanning the sciatic nerve parenchyma from 4 control and 6 *TDP-43^Q331K^* mice. Unpaired t test, p<0.0001 *TDP-43^Q331K^*vs. control.

**Fig. 7v:** N, vessels from tiled ROIs spanning motor nerves from 5 control and 6 *TDP-43^Q331K^* mice. Unpaired t test, p<0.0001 *TDP-43^Q331K^*vs. control.

**Fig. 7y:** N, Ly-6B.2 (7/4)-FITC^+^ neutrophils counted in 29-37 fields of view spanning sciatic and motor nerves from 5 control and 6 *TDP-43^Q331K^* mice. Unpaired t test, p<0.0001 *TDP-43^Q331K^*vs. control.

**Suppl. Fig. 11e:** N, fields of view for MF counts, spanning sciatic nerves from 3 control and 4 *TDP-43^Q331K^* mice. Unpaired t test, p<0.0001 *TDP-43^Q331K^* vs. control.

**Suppl. Fig. 11g, h:** N, vessels from tiled ROIs spanning sciatic nerves from 7 control and 10 *TDP-43^Q331K^* mice. Unpaired t test, p<0.0001 *TDP-43^Q331K^* vs. control.

**Suppl. Fig. 11n:** N, tiled ROIs spanning the sciatic nerve parenchyma from 4 control and 6 *TDP-43^Q331K^* mice. Unpaired t test, (ns) p=0.26 *TDP-43^Q331K^* vs. control.

**Suppl. Fig. 11q, r:** N, tiled ROIs spanning spinal cords from 4 control and 6 *TDP-43^Q331K^* mice. Unpaired t test, p<0.0001 *TDP-43^Q331K^* vs. control.

**Suppl. Fig. 11s:** N, tiled ROIs spanning spinal cords from 7 control and 5 *TDP-43^Q331K^*mice. Unpaired t test, (ns) p=0.78 *TDP-43^Q331K^* vs control.

**Suppl. Fig. 11y, z:** N, vessels from tiled ROIs spanning sciatic nerves from 2 control, 2 *TDP-43^Q331K^*, and 4 *Cdh5^CreER^*; *TDP-43^Q331K^*. One-way ANOVA with Tukey’s test. (**y**) p<0.0001 *TDP-43^Q331K^*vs. control; p<0.0001 *Cdh5^CreER^*; *TDP-43^Q331K^* vs. control; (ns) p=0.12 *Cdh5^CreER^*; *TDP-43^Q331K^* vs. *TDP-43^Q331K^*. (**z**) p<0.0001 *TDP-43^Q331K^* vs. control; p<0.0001 *Cdh5^CreER^*; *TDP-43^Q331K^* vs. control; (ns) p=0.061 *Cdh5^CreER^*; *TDP-43^Q331K^*vs. *TDP-43^Q331K^*.

#### Mice

*SOD1^G93A^* (B6.Cg-Tg(SOD1*G93A)1Gur/J; Jax #004435)^69^. *SOD1^G93A^* transgene copy numbers across generations were controlled by quantitative PCR using with SYBR select master mix (Themo, Cat #4472908) on genomic DNA with primers (forward) 5’-GAAACCGCGACTAACAATCAAAGTG-3’ and (reverse) 5’-CATCAGCCCTAATCCATCTGATGC-3’ for *SOD1*, and (forward) 5’-CAGGTGGGCTCCAGCATT-3’ and (reverse) 5’-TCACCAAGTGTCATTTCTGCCTTTG-3’ for *APOB* as a normalizer. Male and female *SOD1^G93A^* mice were classified as presymptomatic (60 ± 5 days), onset (90 ± 5 days), or symptomatic (120 ± 5 days) based on neuroscore^151^, body weight measurements, and behavioral tests, followed by post-mortem motor neuron and axonal fiber counts. This progression is consistent with previous studies^53^. For consistency, mice are referred to as P60, P90, and P120, with the specific age indicated when analyzed at other time points. *TDP-43^Q331K^*(6.Cg-Tg(Prnp-TARDBP*Q331K)103Dwc/J; Jax #017933)^68^ were staged as presymptomatic (60 ± 10 days), onset (120 ± 10 days), and symptomatic (180 ± 10 days)^68,70^, with both males and females included in the analysis. *Cdh5^CreER^* (Cdh5(PAC)-CreERT2)^110^. *R26-Tomato^LSL^* (Ai14; Jax #007914)^152^. *Ccr2^RFP^Cx3cr1^GFP^*(Jax #032127)^153^. *Ccr2^-/-^* (Jax #004999)^103^. Tamoxifen (Tmx; Sigma-Aldrich Cat# T5648) was dissolved at 10mg/ml in 1:9 ethanol:corn oil (Sigma-Aldrich Cat# S5007) and administered intraperitoneally (i.p.) to *Cdh5^CreER^; TDP-43^Q331K^* mice immediately after weaning for 7 consecutive days at 100mg/kg body weight (BDW). All mouse lines were maintained on C57Bl6/N strain. Both males and females were analyzed unless otherwise specified. Mice were maintained in pathogen-free facilities under standard housing conditions with continuous access to food and water. All experimental procedures and handling were approved by the Animal Research Committee of IRCCS San Raffaele Hospital and performed in compliance with IACUC guidelines.

#### Behavioral analysis

Rotarod test: All mice were trained for 1 week before testing. Mice were placed on an accelerated rotarod (Ugo Basile) from 5 to 40 rotations per minute for a maximum of 300 seconds. A minimum of 5 trials were performed in each session, and the longest latency to fall, normalized to BDW, was recorded^70^. Grip test: aximal force (g) exerted by all four limbs on the grid was measured at least 5 times using a grip strength meter (Ugo Basile). The highest and lowest values were discarded, and the average of the remaining measurements was recorded. BDW was measured every 2 weeks from 2 months of age until the symptomatic stage. For behavioral studies involving neutrophil depletion, only male *SOD1^G93A^* mice were included.

#### Human Biopsy samples

The anterior motor branch of the obturator nerve was sampled and processed according to standardized protocols^116^. Motor nerves from non-ALS neuropathy or myopathy cases were used as controls after applying established exclusion criteria^116^. All biopsies were obtained from Institute of Experimental Neurology (INSPE) tissue bank. Patients provided informed consent to the study. All experimental protocols were approved by San Raffaele Scientific Institute Ethical Committee (Milan, Italy). Axonopathy (demyelination and degeneration) was assessed on semithin sections as part of the clinical diagnostic routine. Alterations in endoneurial vessels (including accumulation of pinocytotic vesicles, tight junction changes, cellular hypertrophy and hyperplasia, basal lamina detachment, and microfilament accumulation) were scored blindly on corresponding TEM sections by two independent researchers, and the average score for each feature was calculated.

#### Electron microscopy and morphological studies

Mice were sacrificed by cervical dislocation and sciatic nerve, femoral nerves with quadriceps and saphenous nerve branches were collected and fixed with 2% glutaraldehyde in 0.12M phosphate buffer, postfixed with 1% osmium tetroxide, dehydrated with alcohol and embedded in Epon (Fluka). Transverse semithin sections (0.5–1μm) of mouse and human nerve samples were stained with toluidine blue and examined by light microscopy. Ultrathin (100–120 nm) sections were stained with uranyl acetate and lead citrate, and examined by TEM (Talos L120C G2).

#### Immunohistochemistry

After dissection, nerves were fixed in 4% paraformaldehyde (PFA) on ice for 2 hours, washed extensively in PBS, cryoprotected in 30% sucrose (in PBS) overnight (ON), embedded in OCT (Sakura Cat# 4583) and cryosectioned at 16-18μm thickness for longitudinal sections and 10μm thickness for transverse sections. For spinal cord analysis, mice were trans-cardiac perfused with saline followed by 4% PFA and dissected and further fixed with 4% PFA ON, cryoprotected with 30% sucrose (in PBS) ON and cryosectioned at 20-25μm thickness. For immunostaining, nerve sections were permeabilized with 0.2% Triton X-100 and blocked in 5% donkey serum diluted in PBS for 1 hr. Spinal cord sections were permeabilized with 0.3% Triton X-100 and blocked in 5% donkey serum diluted in PBS for 1 hr. Primary antibodies were incubated in same solution ON at 4°C. After 3 washes in PBS of 5 min each, secondary antibodies were incubated in 0.1% Triton X-100 and 5% donkey serum diluted in PBS at RT for 2 hrs. After 3 washes in PBS, sections were stained with DAPI and mounted with Fluoromount Aqueous Mounting Medium (Sigma-Aldrich Cat# F4680). For PLVAP, Claudin-5, ZO-1 and VE-cadherin immunostaining, nerves were embedded directly in OCT and frozen on dry ice cooled 2-methylbutane. Sections were fixed with pre-cooled methanol for 5-7 min at 4°C and incubated with primary and secondary antibodies diluted in 1% BSA and 0.01% Triton X-100. For Choline Acetyl Transferase (ChAT) and Myc Immunostaining spinal cords sections were subjected to antigen retrieval with Tris-EDTA buffer for 10 min at 95°C (10mM Tris base, 1 mM EDTA, 0.05% Tween 20 pH 9.0). Samples were routinely imaged with Olympus Fluoview FV3000RS or Leica TCS SP8 SMD FLIM Laser Scanning Confocal microscopes. MAVIG RS-G4 resonant scanning confocal microscope (VivaScope GmbH Research) was used for imaging the entire length of longitudinal nerve sections.

Primary antibodies used for immunostaining were: rat anti-CD31 (1:200; BD Pharm Cat# 550274), goat anti-CD31 (1:1000; R&D Systems Cat# AF3628), rabbit anti-RFP (1:1000; MBL Cat# PM005), rat anti-PLVAP (1:100; BD Pharm Cat# 553849), rabbit anti-claudin5 (1:400, Thermo Fisher Scientific Cat# 34-1600), goat anti-Podocalyxin (1:400; R&D Systems Cat# AF1556), rat anti-CD144 (1:100; BD Pharm Cat# 550548), rabbit anti-VWF (1:200; Dako Cat# A008229-5), chicken anti-GFP (1:500; Abcam Cat# ab13970), rabbit anti-GFP (1:1000; Thermo Fisher Scientific Cat# A6455), rabbit anti-Fibrinogen (1:10000; Dako Cat# A0080), rabbit anti ZO-1 (1:300; Thermo Fisher Scientific Cat# 40-2300), rat anti-VCAM1 (1:100; BD Pharm Cat# 550547), goat anti-ICAM1 (1:150; R&D Systems Cat# AF796), goat anti-Car4 (1:400; R&D Systems Cat# AF2414), , Rabbit anti-Iba1 (1:500; Wako Cat# 019-19741), rabbit anti-GFAP (1:400; Dako Cat# Z0334), mouse anti-NeuN (1:100; Millipore Cat# MAB377), rabbit anti-ChAT (1:2000; Abcam Cat#AB178850), mouse anti-Myc (1:100; Cell signaling Cat#2276), rat anti-F4/80 (1:250; Abcam Cat# AB6640), rabbit anti-Caveolin (1:300; Cell signaling Cat# 3267), rabbit anti-Collagen IV (1:500; Biorad Cat# 2150-1470), rat anti-Ly6g (1:100; BD Pharm Cat# 551459), rat anti-7/4 FITC (1:200; Abcam Cat# AB53453), rabbit anti-Mfsd2A (1:400, Cell signaling Cat# 80302). Secondary antibodies: goat or donkey anti-rabbit/rat/goat/mouse Alexa Fluor 488-, 555-, 647-conjugated (1:500; Thermo Fisher Scientific).

#### Whole nerve staining and tissue clearing

Nerves were fixed overnight in 4% PFA, and the epineurium was manually removed. Nerves were incubated in PBSGT (0.2% gelatin, 0.5% TritonX-100, 0.01% Thimerosal, Sigma-Aldrich Cat# T8784 in PBS) for 2 days at RT on a rocker. Primary antibodies were diluted in PBSGT with 10mg/mL saponin and incubated at RT for 7 days on a rocker. Samples were washed 6 times over one day with PBSGT, then incubated with secondary antibodies in PBSGT with 10mg/mL saponin for 2 days at 37°C on a rocker. After 6 washes in PBSGT, nerves were stored in PBS at 4°C until clearing with X-CLARITY Tissue Clearing System (Logos Biosystems). Samples were incubated in Hydrogel Solution (Logos Bio-systems Cat# C1310X) for 24 hours at 4°C, transferred in X-CLARITY Polymerization System for 3 hrs at 37°C, then washed several times in Electrophoretic Tissue Clearing Solution (Logos Biosystems Cat# C13001). Nerves were clarified using the X-CLARITYTissue Clearing System II (Logos Biosystems C30001) to remove lipids from the tissue, applying a current of 0.6A at 37°C for 4 hours. Samples were washed extensively in PBS and incubated in X-Clarity Mounting Solution (Logos Bio-systems Cat# C13101) for 1 hour at RT before imaging with a Nikon A1 MP multiphoton microscope with Coherent Chameleon Ultra II using Apo LWD 25 X (NA 1.1) Water Plan objective. 3D-image stacks were analyzed with Arivis 4D software. Primary antibodies used for whole nerve staining were a mix of goat anti-CD31 (1:500; R&D Systems Cat# AF3628) and goat anti-Podocalyxin (1:1000; R&D Systems Cat# AF1556). Secondary antibodies (from Thermo Fisher Scientific) were donkey anti-goat AlexaFluor-555 (1:1000; Cat# A21432).

#### Vascular permeability assays

Adult mice received 200μl of EZ-Link Sulfo-NHS-LC-Biotin (Thermo Fisher Scientific Cat# 21335) (20mg/ml in PBS) via tail vein injection and were sacrificed after 30 min to allow circulation of the tracer. Nerves and spinal cords were harvested for immunostaining and the dye was detected using AlexaFluor 555- or 647-streptavidin (1:500; Thermo Fisher Scientific Cat# S32355, S32357). Mice that received BSA-FITC (Invitrogen Cat# A23015) (1.5mg/ml in PBS; 10μl/g BDW) via tail vein injection were sacrificed after 2-3 hours for nerve harvesting^60^. Mice tail vein injected with HRP (Sigma Aldrich Cat# P8250; 500mg/kg BDW in PBS) were sacrificed after 30 min. HRP was detected on nerve sections using TSA Fluorescein system (Akoya Biosciences Cat# NEK701A001KT).

#### Endothelial cell isolation from sciatic nerve

Sciatic nerves were harvested in DMEM/F-12 medium (Gibco Cat# 11039021). The epineurium was manually stripped before dissociation for bulk RNA-seq in order to enrich for endoneurial ECs, while it was not removed for scRNA-seq to capture EC from the entire nerve vasculature. Nerves were pooled by genotype, minced on ice in DMEM/F-12 using a razor blade and rotated for 30 min at 37°C in dissociation buffer [Collagenase Type II (50ml/ml from 200mg/ml stock; Gibco Cat# 17101015), Hyaluyronidase (1:50 from 1% stock; Sigma-Aldrich Cat# H3506), DNase I (2.5μg/ml; Sigma-Aldrich Cat# DN25) in DMEM/F- 12] with trituration by pipetting every 10 min. The reaction was blocked with 5% FBS/2mM EDTA in PBS w/o Ca2^+^/Mg2^+^. The cell suspension was passed through a 70mm cell strainer and centrifuged at 300 xg for 10min (4°C). The pellet was washed twice in FACS buffer [5% FBS in PBS w/o Ca2^+^/Mg2^+^], resuspended in 0.5% BSA/PBS w/o Ca2^+/^Mg2^+^ and incubated with myelin removal beads (Miltenyi Biotech Cat# 130-096-731) for 15 min, and passed through LS columns (Miltenyi Biotech Cat# 130-042-041) to remove myelin debris. The flowthrough cell suspension was collected and stained with rat anti-CD31 APC (1:50; BD Pharmingen Cat#551262) and rat anti-CD45.2 V405 (1:250; BD Biosciences Cat# 560697) at 4°C for 30 min with intermittent tapping. Samples were washed twice with FACS buffer and resuspended in the same buffer. ECs (CD31^+^/CD45^−^) were sorted with BD FACSAria Fusion flow cytometer into 400μl of Trizol-LS (Thermo Fisher Scientific Cat # 10296010) for bulk RNA-seq or DMEM/F-12 medium for scRNA-seq.

#### EC isolation from spinal cord

Spinal cords were dissected from the same mice used for sciatic nerve EC isolation described above and pooled by genotype. Each spinal cord was chopped on ice into 3-4 pieces in DMEM/F-12 medium. Tissue dissociation was performed with the Miltenyi Adult Brain Dissociation kit (Cat# 130-107-677). Samples were incubated with the dissociation enzyme mix, placed on Miltenyi gentleMACS Octo dissociator for 30 min, and passed through a 70μm cell strainer. After red blood cell removal, samples were resuspended in 2% BSA/PBS w/o Ca2^+^/Mg2^+^ and stained with with anti-CD31 and anti-CD45.2 antibodies for FACS-purification of CD31^+^/CD45^−^ ECs as indicated above. ECs were sorted in 400μl of Trizol-LS for total RNA isolation and bulk RNA-seq.

#### Bulk RNA-sequencing and bioinformatic analysis

For each replicate, 6-7 sciatic nerves and 3-4 spinal cords were pooled by genotype and dissociated as described above: Control (non-transgenic) P90 sciatic nerves (3 independent replicates), *SOD1^G93A^* P90 sciatic nerves (3 replicates), Control P120 sciatic nerves (3 independent replicates), *SOD1^G93A^* P120 sciatic nerves (3 replicates), Control P90 spinal cords (3 replicates), *SOD1^G93A^* P90 spinal cords (3 replicates), Control P120 spinal cords (4 replicates), *SOD1^G93A^*P120 sciatic nerves (4 replicates). An average of 20,000 ECs were purified by FACS for each replicate. Total RNA was prepared as per manufacturer’s instructions and resuspended in 10-12ml molecular-grade water. After quantification and quality control with Qubit 3.0 Fluorometer (Thermo Fisher Scientific) and Agilent 2100 Bioanalyzer, RNA-seq libraries were generated using the SMART-Seq Ultra Low Input RNA Kit (Illumina) and sequenced on an Illumina NovaSeq6000 system (single-end, 100bp read length). Sequencing reads were trimmed using Trimmomatic (v0.39)^154^ to remove sequencing adapters and exclude low-quality reads, and aligned to the mm10 reference genome using STAR aligner (v2.5.3a)^155^. Transcript quantification (total counts per gene) was performed using FeatureCounts (v1.6.4)^156^ based on mapping to exons annotated in GENCODE Mus musculus basic gene annotation (version M16), excluding multimapping reads. For each comparison, only genes with average expression greater than 3 RPKM (Reads Per Kilobase per Million mapped reads) across samples were retained for downstream analysis. Differential gene expression analysis was conducted using the DESeq2 package (v1.30.1)^157^. To visualize gene expression patterns, RPKM values were transformed to z-scores across samples using the pheatmap R package. GO analysis of DEGs (padj ≤0.05) was performed using Enrichr^158^. Enriched terms (padj <0.01) were selected across relevant gene-set libraries. The resulting list was further curated to prioritize the most significant and representative processes, excluding redundant or overly generic terms. GSEA was performed using all expressed genes (cpm > 1) as input, with the clusterProfiler package (v4.2.2)^159^. Enrichment, volcano, GSEA and expression Dot plot were generated with ggplot2 (v4.2.3). EWCE (Expression Weighted Cell-type Enrichment) package (v1.2.0)^160^ was applied to DEGs showing consistent changes at P90 and P120 using the integrated control/*SOD1^G93A^*EC nerve scRNAseq dataset as a reference. Cell-type abundance was inferred from bulk RNA-seq results with CIBERSORTx19^161^ using raw count matrices as input. For spinal cord samples, reference profiles were derived from the Brain Tabula Muris (https://figshare.com/articles/dataset/5829687). For sciatic nerve samples, reference signatures were obtained from an in-house sciatic nerve single-cell atlas.

#### Single-cell sequencing and bioinformatic analysis

Approximately 20 intact sciatic nerves were pooled for each group (Control vs. *SOD1^G93A^* and dissociated as described above. Single-cell suspensions were stained as indicated above with a mix of antibodies to label ECs (rat anti-CD31 APC; 1:50) along with myeloid cells (rat anti-CD45.2 V405; 1: 400), mesenchymal cells (rabbit anti-NG2 Alexa Fluor488; Millipore Cat# AB5320A4; 1:200), and Schwann cells (mouse anti O4 PE; Miltenyi Biotec Cat# 130-117-357; 1:250). All CD31⁺ ECs were sorted, along with a proportion of other labeled and unlabeled cells, to ensure representative inclusion of various nerve cell types without over-representing the most abundant population, particularly Schwann cells. Sorted cells were counted manually and with an automatic cell counter, and resuspended in 2% FBS/DMEM/F-12 at a concentration of 1000 cells/ml. Cells were subsequently encapsulated and processed using the Chromium platform (10X Genomics) with the Chromium Single Cell 3’ Library & Gel Bead Kit v3. After quality controls and quantification on TapeStation instrument (Agilent), libraries were sequenced on Illumina NovaSeq6000 system. From 3 independent cell purification and scRNA-seq experiments, we generated 2 Control and 4 *SOD1^G93A^* samples. Raw sequencing data were demultiplexed using the mkfastq application from CellRanger (v6.1.2, 10x Genomics). Sequencing libraries were aligned to the 10x Genomics default mm10 reference genome (version 2020-A) using the CellRanger pipeline^162^. The resulting raw gene expression matrices were processed using the Seurat R package (v5.0.0)^163^, with default parameters unless otherwise specified. Only cells (unique barcodes) that expressed a minimum of 300 genes (considering only genes detected in at least 5 cells) and contained over 1000 unique molecular identifiers (UMIs) were retained. Cells with more than 15% of total reads mapping to mitochondrial genes were excluded. After filtering, a total of 35,793 single cells were retained, comprising 16,962 cells from non-transgenic control samples and 18,831 cells from *SOD1^G93A^* samples. Following quality control, datasets were integrated using Seurat’s Harmony-based integration (v0.1.1)^164^ with SCTransform normalization^165^ and batch correction regressing out mitochondrial gene expression to remove technical variation. Principal component analysis was performed (with RunPCA) and the number of PCs was selected after inspection with ElbowPlot (nPCs=50). Cell clustering was performed using FindNeighbors and FindClusters with a resolution of 0.1. Cell clusters were annotated based on expression of canonical marker genes and visualized on UMAP plots with the “FeaturePlot” and DimPlot functions. From this dataset, ECs (3,062 cells) were extracted, re-integrated and re-clustered following the procedure described above with a clustering resolution of 0.5. Rare contaminating cells were removed. Marker genes for each EC subcluster, and DEGs between control and *SOD1^G93A^* samples for each EC subtype cluster were detected using the FindAllMarkers function (thresh.use=0.1, min.pct=0.01, min.diff.pct=-Inf e test.use = "wilcox"). Endoneurial EC selected based on known marker genes^64^ were re-integrated and re-clustered with a resolution of 0.2. Marker gene detection, cell-type annotation and DEGs analysis were performed as described above. To assess gene expression patterns, average log-normalized expression values were transformed to z-scores across cell populations using the pheatmap R package. Gene signatures were generated using the AddModuleScore function in Seurat to quantify module expression scores across cells. Single cell data were visualized using UMAP. Alluvial plot was generated using the geom_alluvium function from the ggalluvial R package (v0.12.3)^166^.

To compile a single-cell atlas of the sciatic nerve endothelium, Seurat v4.0 workflow was used to integrate 5 previously published scRNA-seq datasets of the intact mouse sciatic nerve^64,89–92^, along with the control nerve samples obtained in this study. Variable genes (n=2000) were identified in each sample using the *FindVariableFeatures* function. Integration anchors were computed with *FindIntegrationAnchors* (dims = 1:30) and used to integrate the datasets via the *IntegrateData* function. The integrated dataset was then scaled and centered using *ScaleData*, regressing out unwanted sources of variation such as mitochondrial gene expression and dataset of origin. Cell clustering was performed using *FindNeighbors* and *FindClusters* with a resolution of 0.1. ECs annotated based on the expression of marker genes were extracted and re-integrated following the same workflow, using a clustering resolution of 0.2. The contribution of each dataset was as follows: Bhat et al. (514 ECs), Gerber et al. (1544 ECs), Zhao et al. (828 ECs), Toma et al. (380 ECs), Wolbert et al. (488 ECs), this study (1201 ECs). Small cell clusters that could not be annotated with known EC-subtype marker genes were eliminated yielding to a total of 4894 ECs in the atlas. Dendrogram visualization of hierarchical clustering of EC subtypes was performed with pheatmap R package (v1.0.12)^167^ using the mean expression values of the top 200 marker for each subtype cluster, ranked by log2 fold change.

The reanalysis of scRNA-seq dataset of spinal cord endothelium (GSE210776)^94^ was performed with the standard Seurat workflow (v5.0.0). Integrated analysis was carried out using the IntegrateLayers function, with CCAIntegration method. The integrated object was then used to compute a nearest-neighbor graph (FindNeighbors, dims = 1:30, reduction = "integrated.cca"). Cell clustering was performed using FindClusters at resolution 0.1. UMAP was applied for visualization (RunUMAP, dims = 1:30, reduction = "integrated.cca"). Cldn5⁺ ECs were extracted and re-clustered applying the same workflow with 0.15 resolution. To compare the transcriptomic profiles of sciatic nerve vs. spinal cord ECs we performed cross-dataset label transfer using the FindTransferAnchors and TransferData functions (dims = 1:30) from the Seurat package to projected EC subtype annotations from the sciatic nerve EC atlas onto the spinal cord single-cell dataset.

#### RNA purification and quantitative RT-PCR from human motor nerve biopsies

Nerve biopsies flash-frozen in liquid nitrogen were stored at the San Raffaele Hospital-INSPE tissue bank. Tissue homogenates were prepared using Precellys Evolution tissue homogenizer equipped with Cryolys Evolution cooling system (Bertin Technologies). Briefly, frozen nerves were transferred into tubes containing beads and 500µl of TRIzol (Invitrogen, Cat# 15596026) and homogenized at 6000 rpm for 4 × 30 s with 60 s pauses. Lysates were centrifuged, and supernatants were collected for RNA extraction following the standard TRIzol protocol. cDNA was synthesized from 0.3-1mg RNA with M-MLV reverse transcriptase (Thermo Fisher Scientific Cat# 28025013) and random primers (Thermo Fisher Scientific Cat# N8080127), diluted to 40μl in molecular biology-grade water and used for real-time quantitative PCR with SYBR Select Master Mix (Thermo Fisher Scientific Cat# 4472913). RT-PCR reactions were performed in triplicate. Primers: *MFSD2A* forward 5’-CTCCTGGGCCATCATGCTCTC-3’; reverse 5’-GGCCACCAAAGATGAGAAA-3’. *GAPDH* forward 5’-ATCACCATCTTCCAGGAGCGAG-3’; reverse 5’-GGGCAGAGATGATGACCCTTTTG-3’.

#### Western blotting

Mouse nerves were flash-frozen with liquid nitrogen immediately after harvesting, triturated manually with pestels on dry ice, and resuspended in SDS-lysis buffer (150mM NaCl, 50mM TrisHCl pH7.4, 10mM EDTA, 1% Tx-100, 0.1% SDS; 150μl/ nerve) supplemented with protease (Roche Cat# 4693132001) and phosphatase (Thermo Fisher Scientific Cat# 78420) inhibitor cocktails, followed by bath sonication for 30 seconds at full power. Lysates were incubated on a thermomixer at 700 rpm for 30 min at 4°C and centrifuged at 13,000g for 15 min (4°C). Supernatants were collected and quantified with Bradford protein assay (Bio-Rad Cat# 5000006). Protein samples were run on handcast 8-10% SDS-PAGE gels or NuPAGE 4-12% Bis-Tris gels (Thermo Fisher Scientific) and transferred on nitrocellulose membranes (Amersham/Merck Cat# GE10600002). Antibodies used for immunoblotting were: goat mouse anti-Vinculin (1:5000, Millipore Cat# 05-386), mouse anti-Caveolin (1:1000; BD Pharm Cat# 610407), mouse anti-pY14 Caveolin (1:1000; BD Pharm Cat# 611339), rabbit anti-Claudin5 (1:1000, Thermo Fisher Scientific Cat# 34-1600).

#### *In vivo* neutrophil depletion

Neutrophils were depleted by intraperitoneal injection of InVivoMAb anti-mouse Ly6G antibody (clone 1A8; Catalog #BE0075-1, BioXcell) at a dose of 10 mg/kg BDW every four days. Control mice received an equivalent dose of InVivoMAb rat IgG2a isotype control antibody, anti-trinitrophenol (clone 2A3; Catalog #BE0089, BioXcell). Neutrophil depletion was confirmed by flow cytometry analysis of peripheral blood collected from the retro-orbital sinus of anesthetized mice the day before and the day after antibody administration. Blood was drawn into sodium heparin-coated hematocrit capillaries, and red blood cells were lysed using ACK Lysing Buffer. The resulting single-cell suspensions were washed twice with PBS and used for subsequent staining and analysis. Only males were used for this experiment to ensure consistency in body weight measurement and behavioral tests across groups.

#### Immune cell isolation and flow cytometry

Single-cell suspensions from mouse sciatic nerves were prepared as described above. For flow cytometry staining, cells were plated at a maximum density of 2×10^6^ cells per well in a 96-well U-bottom tissue culture plate. Cell viability was assessed with Viability 405/520 fixable dye (Miltenyi, Cat #130-130-404) or LIVE/DEAD Fixable Far-Red dye (Invitrogen, Cat #L34973). Staining of surface markers was carried out at 4°C for 30 min in the presence of anti-CD16/32 blocking antibody (Fc-block, BD Biosciences #553142) with Brilliant Stain buffer (BD Biosciences Cat #566349) according to manufacturer instructions, using the following antibodies: anti-CD45 (clone 30-F11, BioLegend Cat #103111), anti-CD45.2 (clone 104, BD Biosciences Cat #564616), anti-CD4 (clone RM4-5, eBioscience Cat #48-0042-82), anti-CD8a (clone 53-6.7, BD Biosciences Cat #612898), anti-CD80 (clone 16-10A1, BioLegend Cat #104738), anti-CD86 (clone GL1, BD Bioscience Cat #564199), anti-IA-IE (clone M5/114.15.2, BD Biosciences Cat #748846), anti-H2-kB (clone AF6-88.5, BD Biosciences Cat #742861), anti-F4/80 (clone BM8, BioLegend Cat #123110), anti-CD11c (clone N418, BioLegend Cat #117319), anti-CD11b (clone M1/70, BD Pharmingen Cat #557960), anti-CD68 (clone FA-11, BioLegend Cat #137015), anti-CCR5 (clone HM-CCR5, BioLegend Cat #107005), anti-CX3CR1 (clone SA011F11, BD Biosciences Cat #149029), anti-Ly6G (clone 1A8, BioLegend Cat #127624), anti-CCR3 (clone J073E5, BioLegend Cat #144508), anti-Ly6C (clone HK1.4, BioLegend Cat #128014). Flow cytometry was performed in FACS buffer [2mM EDTA, 2% FBS (Corning) in PBS] using BD FACSymphony A5 Cell Analyzer or Cytek Aurora. Flow cytometry data were analyzed with FlowJo Software (Treestar).

#### Image Analysis, quantification and statistics

Images used for quantification were acquired using the same microscope settings and adjusted to the same background levels whenever possible. Images were processed with Fiji/ImageJ, Adobe Photoshop and Adobe Illustrator. 3D rendering and videos were generated with Arivis 4D software. Mean intensity values and px area of various antigens were measured with Fiji/ImageJ on a binary mask of either CD31 or Podocalyxin staining, as indicated. Vessel width was measured by line tool from Fiji/ImageJ and plotted in px length. For permeability assays the extravascular intensity value of sulfo-biotin/HRP/BSA in nerves or spinal cord were quantified after subtracting its signal from vessels masked by CD31. CAV1 staining were generated from individual confocal images of control or SOD1G93A nerve ECs acquired at 60× magnification with 1.5× optical zoom. Microglia and astrocyte activation in *SOD1^G93A^* and *TDP43^Q331K^* mice were quantified by measuring IBA1^+^ or GFAP^+^ px area, respectively. Macrophages or neutrophils were manually counted by F4-80 or Ly-6B.2 (7/4) staining, respectively, within the same px area across samples. CAR4^+^ and PLVAP^+^ vessels were counted manually within random areas of the same size in control, *SOD1^G93A^*or *TDP43^Q331K^* nerve sections. Profile intensity plots for Motor neurons were manually counted in lumbar spinal cord sections based on NeuN or ChAT staining. Myc^+^ nuclei were counted within a fixed area of spinal cord or sciatic nerve transverse sections from *TDP43^Q331K^* mice. GFP^+^ or RFP^+^ cells labeled by *Cx3cr1^GFP^*/*Ccr2^RFP^* reporter were counted after anti-GFP/RFP staining within the same area of longitudinal nerve sections across control and mutant samples. Nerve fibers were counted manually in areas of 300 px^2^ from semithin sections. Pinocytotic vesicles were manually counted in TEM images of ECs and normalized by cytoplasmic area. Statistical analysis was performed with GraphPad Prism V8.4.0 software. Statistical details are found in the figure legends.

---

**Supplementary Fig. 1:**
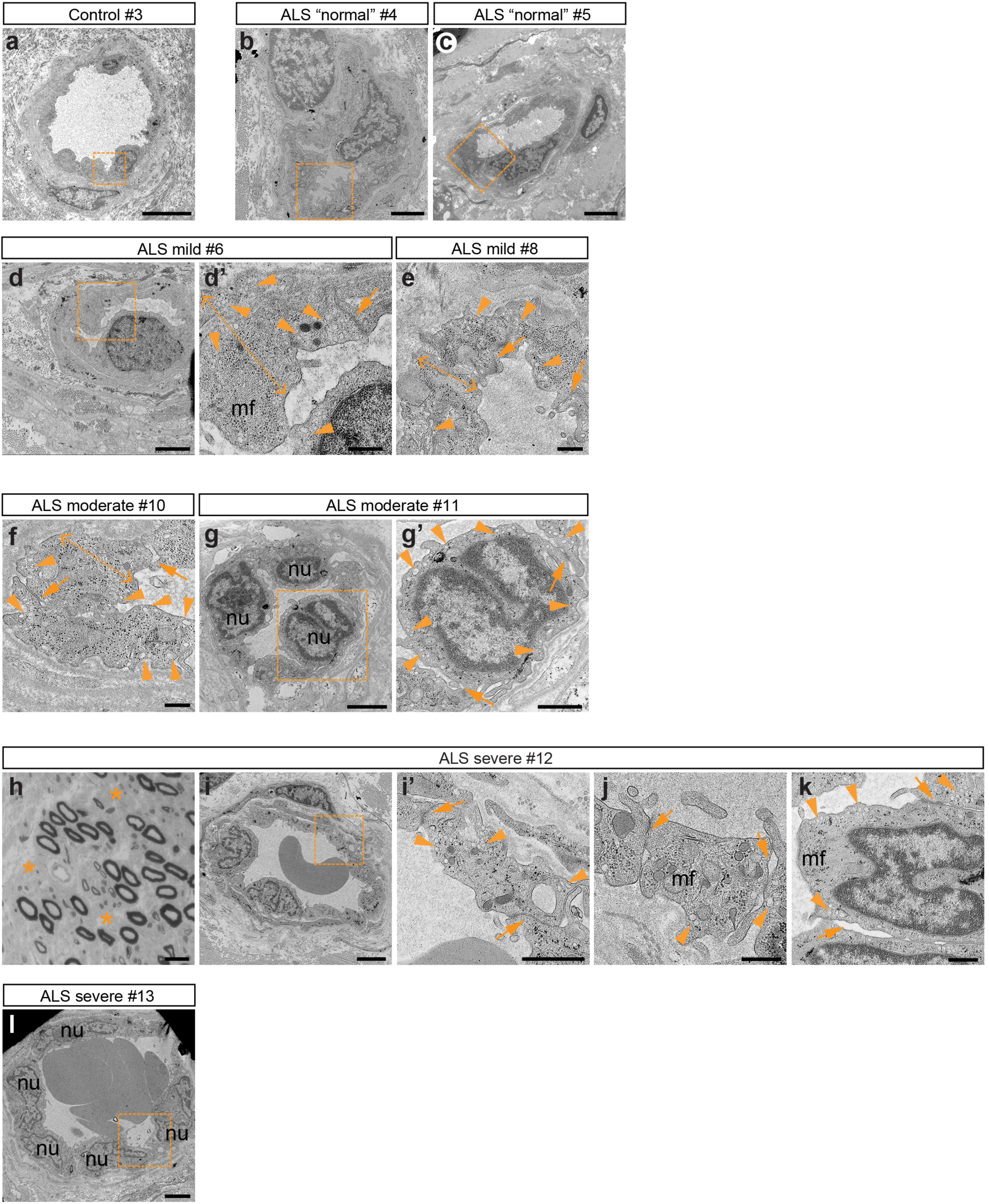
Ultrastructural endothelial defects in patient motor nerve biopsies. TEM images of control (myopathy) *obturator* nerve biopsy (**a**) and sporadic ALS cases of varying severity (**b-g’**, **i-l**). Accumulation of pinocytic vesicles (closed arrowheads) is visible in all ALS samples. TJs (arrows) are partially open in severe cases (**j**, **k**). Hypertrophy is evident in **d**, **d’**, **e**, **f** (double open arrowheads span the endothelium). EC hyperplasia is visible in **g**, **l** (nu, nuclei). Microfilament bundles (mf) are present in **d’**, **j**, **k**. Boxed areas in **a**, **b**, **c**, **l** are magnified in Fig. 1d’, e’, f’, h’, respectively. Boxed areas in **d**, **g, i** are magnified in **d’**, **g’ i’**. **h**, Toluidine Blue-stained semithin transverse sections of diagnostic *obturator* nerve biopsies from severe ALS case with marked axon loss (asterisks). This sporadic patient carried the *SOD1* p.G148S mutation while genetic changes were not identified for the other cases. Scale bars: a, 5μm; b, c, d, g, i, l, 2μm; g’, i’, 1μm; d’, e, f, j, k, 500nm; h, 20μm

**Supplementary Fig. 2:**
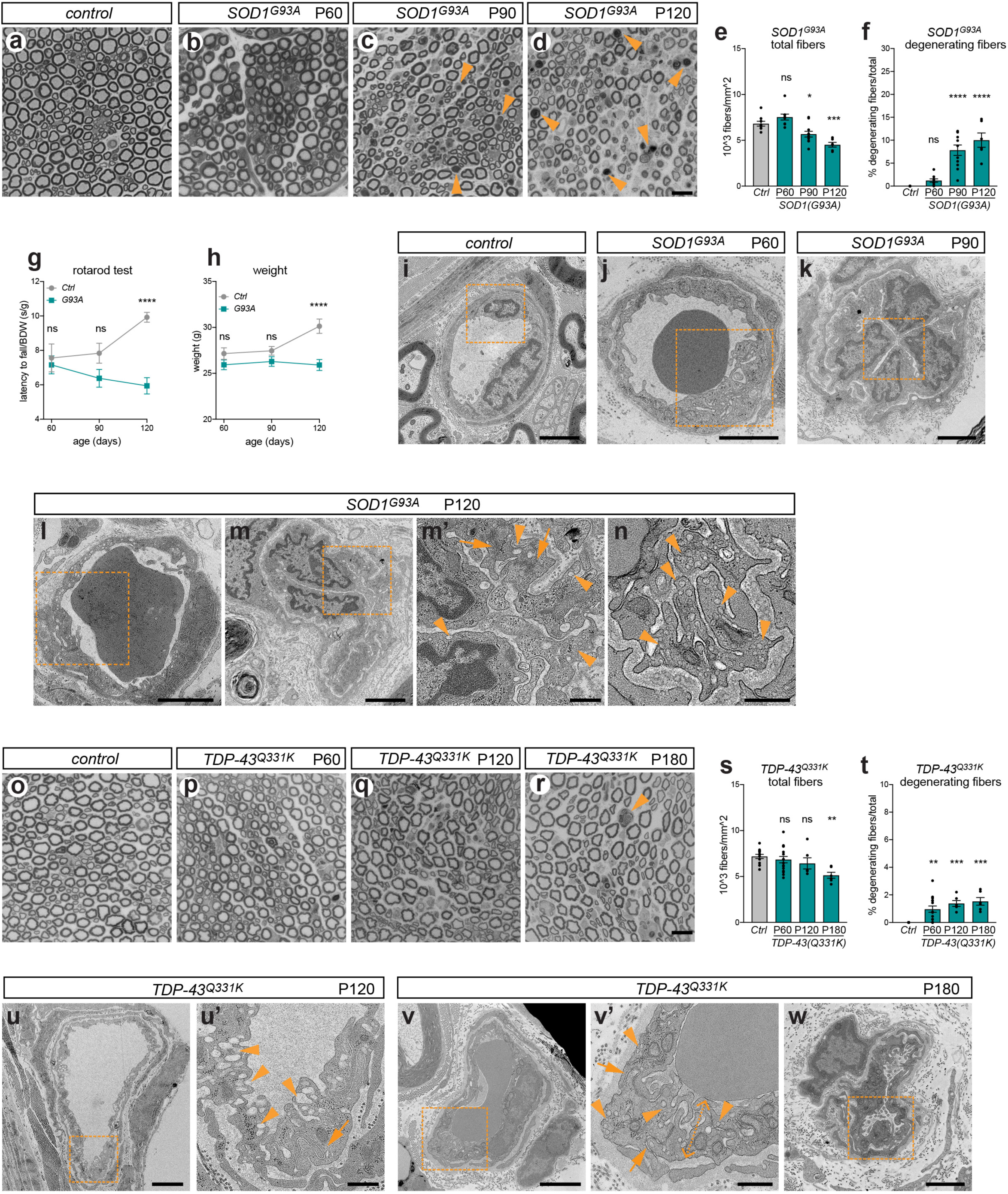
Endothelial alterations precede axonal degeneration in *SOD1^G93A^* and *TDP-43^Q331K^* ALS mice. **a-d**, Semithin transverse sections of control (**a**) and *SOD1^G93A^* (**b-d**) *quadriceps* motor nerves during disease progression. Arrowheads point to degenerating fibers. **e**, **f**, Axon density (**e**) and percentage of degenerating axons (**f**) assessed in semithin sections from control and *SOD1^G93A^* motor nerves. Mean ± SEM. **g**, Rotarod motor performance of control and *SOD1^G93A^* mice. Mean ± SEM; normalized to body weight, BDW). **h**, Body weight measurement in control and *SOD1^G93A^* mice. Mean ± SEM. **i-n**, TEM images of endoneurial vessels in motor nerves of control (**i**) and *SOD1^G93A^* (**j-n**) mice at the indicated stage. Boxed areas in **i-l** are magnified in Fig. 1n-q. Abundant cytoplasmic vesicles are visible in *SOD1^G93A^* ECs (arrowheads in **m’**, **n**). TJs are preserved (arrows, **m’**). **o-r**, Semithin transverse sections of control (**o**) and *TDP-43^Q331K^* (**p-r**) motor nerves during disease progression. (arrowhead in **r** points to degenerating fiber). **s**, **t**, Axon density (**s**) and percentage of degenerating axons (**t**) assessed in semithin sections from control and *TDP-43^Q331K^* motor nerves. Mean ± SEM. Mild axonal loss occurs in the early-symptomatic phase (P180), with only a few actively degenerating fibers visible (compare to *SOD1^G93A^*, **f**). Increased inter-fiber distance in **r** is a sign of edema. **u-w**, TEM images of endoneurial vessels in motor nerves of *TDP-43^Q331K^*mice at P120 (onset) and P180 (early-symptomatic). Boxed areas in **u**, **v** are magnified in **u’**, **v’**, respectively. Arrowheads indicate cytoplasmic vesicles. Arrows point to preserved TJs. Double open arrowheads mark hypertrophy (**v’**). The boxed area in **w** is magnified in Fig. 1r. Scale bars: a-d, o-r, 20μm; i, j, k, l, m, u, v, w, 2μm; m’, n, u’, v’, 500nm

**Supplementary Fig. 3:**
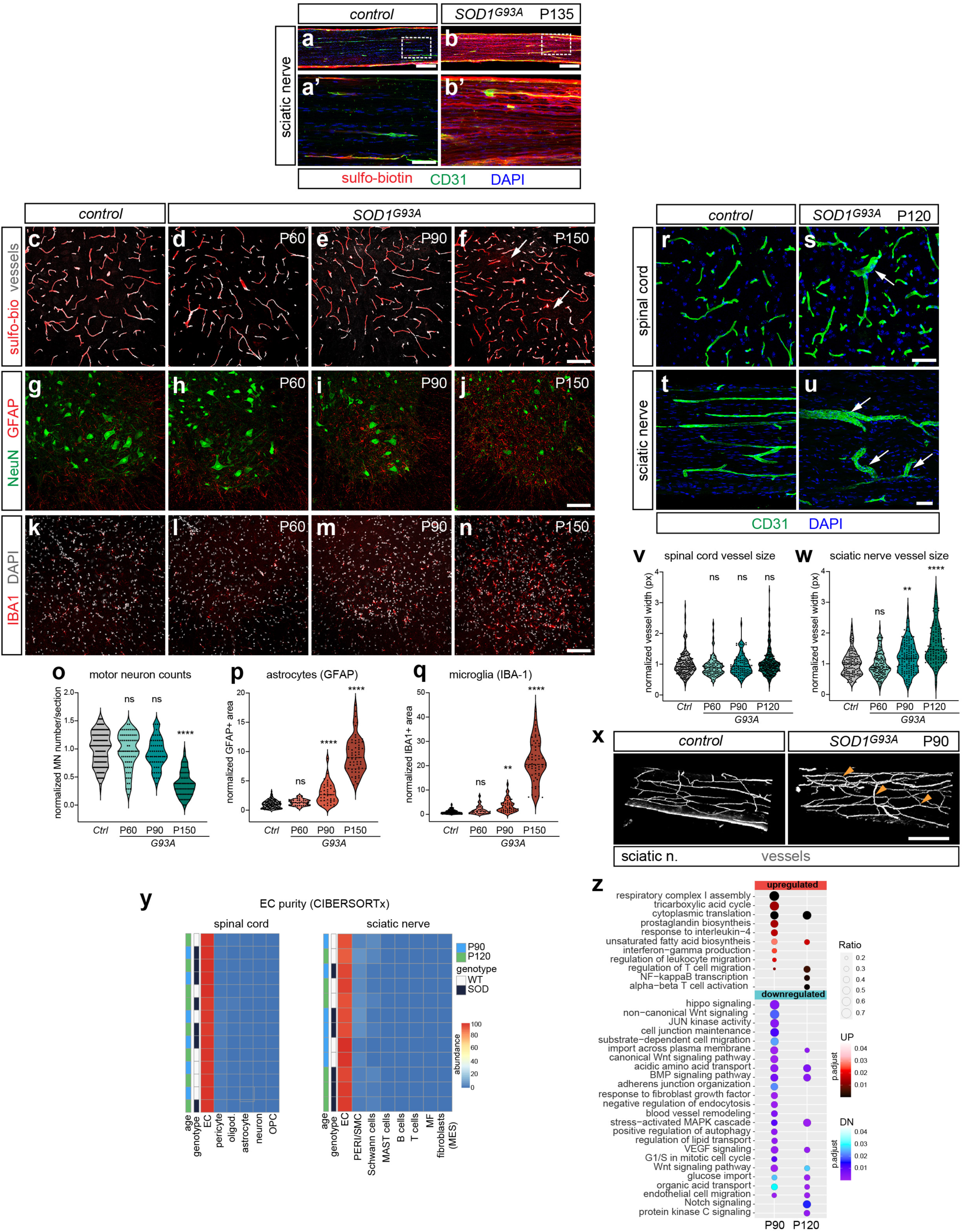
Enhanced microvascular barrier defects in *SOD1^G93A^* peripheral nerves compared to the spinal cord. **a-b’**, Longitudinal views of sciatic nerves from the same control and *SOD1^G93A^* mice at P135 as in Fig. 1k-l’ (where spinal cords are shown). Sulfo-biotin leakage is widespread in nerves. Boxed areas in **a**, **b** are magnified in **a’**, **b’**. **c-f**, Spinal cords from control and *SOD1^G93A^* mice at P60 (**d**), P90 (**e**), and P150 (**f**) following sulfo-biotin injection. The tracer remains confined to the vasculature until late disease stage, when focal leakage is observed (arrows, **f**). Vessels are labeled with Podocalyxin. **g-j**, Control and *SOD1^G93A^* spinal cords showing motor neurons in the ventral horn (NeuN) and reactive astrocytes (GFAP). Motor neuron loss and astrocytosis are conspicuous in the symptomatic phase (P150, **j**). **k-n**, Control and *SOD1^G93A^* spinal cords showing proliferation/activation of microglia (IBA1) in the ventral horn of mutant mice. DAPI, nuclei. **o**, Number of NeuN^+^ motor neurons in spinal cord sections of control and *SOD1^G93A^* mice at progressive stages, normalized to average control value. **p**, **q**, Quantification of astrocytes (GFAP^+^) (**p**) and microglia (IBA1^+^) (**q**) in control and *SOD1^G93A^* spinal cords. Immunoreactivity area normalized to average control value. **r-u**, Spinal cord (**r**, **s**) and sciatic nerve (**t**, **u**) vessels from P120 control and *SOD1^G93A^*mice stained for CD31. Vessel dilation is frequent in *SOD1^G93A^* nerves (**u**, arrows) but rare in the spinal cord (**s,** arrow). DAPI, nuclei. **v**, **w**, Vessel diameter in spinal cord (**v**) and sciatic nerve (**w**) of control and *SOD1^G93A^* mice across disease stages, normalized to average control value. **x**, 3D rendering from two-photon imaging of sciatic nerve vasculature stained for CD31/Podocalyxin in P90 control and *SOD1^G93A^* mice. Mutants show discrete architectural changes, including increased anastomotic branching (arrows). **y**, Cell type composition inferred from bulk RNA-seq data using CIBERSORTx confirms efficient EC purification. Color scale indicates cell abundance. Tabula Muris and sciatic nerve scRNA-seq datasets were used to deconvolve spinal cord and nerve EC transcriptomes, respectively (see Methods). **z**, GSEA identifies upregulated (red) and downregulated (blue) pathways in *SOD1^G93A^* at P90 and P120. Scale bars: a, b, 200μm; a’, b’, 50μm; c-n, 100μm; r-u, 50μm; x, 250μm.

**Supplementary Fig. 4:**
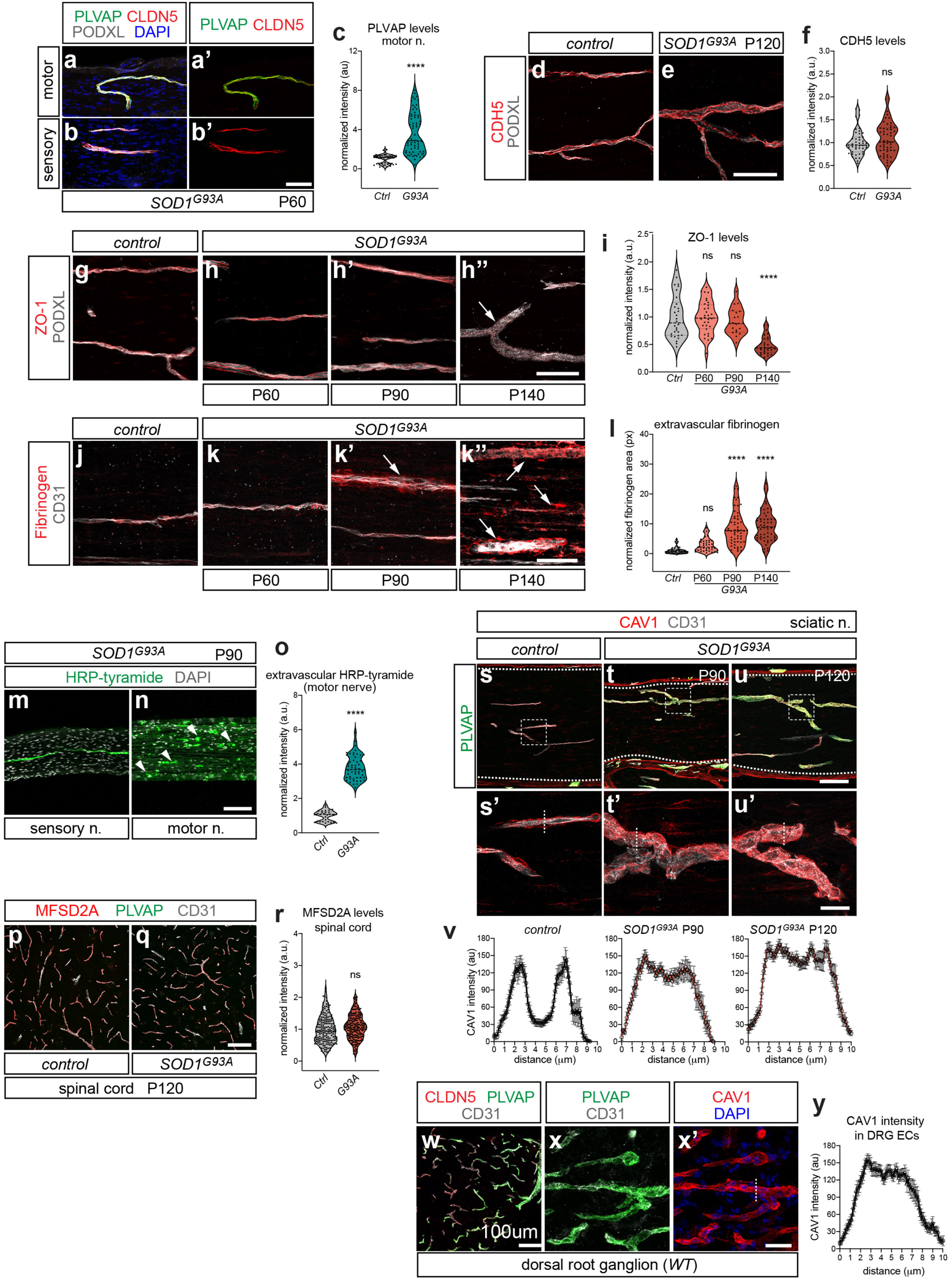
Deregulated transcytosis at the blood-nerve barrier precedes endothelial junctional alterations. **a-b’**, Endoneurial vessels in *quadriceps* (motor) (**a**, **a’**) and *saphenous* (sensory) (**b**, **b’**) nerves from P60 *SOD1^G93A^* mice stained for PLVAP, CLDN5, and PODXL (marks all vessels). DAPI, nuclei. **a’**, **b’** show only PLVAP and CLDN5 staining. **c**, PLVAP signal in endoneurial vessels of P60 control and *SOD1^G93A^*motor nerves, normalized to average control value. **d**, **e**, Endoneurial vessels in control (**d**) and *SOD1^G93A^* (**e**) sciatic nerves at P120 stained for VE-cadherin (CDH5) and PODXL (marks all vessels). **f**, CDH5 signal in endoneurial vessels of control and *SOD1^G93A^*sciatic nerves, normalized to average control value. **g-h’’**, ZO-1 staining in endoneurial vessels of control (**g**) and *SOD1^G93A^*(**h**, **h’**, **h”**) sciatic nerves at progressive stages. Control is P90. PODXL, pan-vessel marker. ZO1 decreases in *SOD1^G93A^*at advanced stage (**h”**, arrow). **i**, ZO-1 signal in endoneurial vessels of control and *SOD1^G93A^* sciatic nerves during disease progression, normalized to average control value. **j-k”**, Fibrinogen staining in endoneurial vessels of control (**j**) and *SOD1^G93A^* (**k**, **k’**, **k”**) sciatic nerves at progressive stages. Control is P90. CD31, pan-vessel marker. Fibrinogen extravasation is evident at P90 and increases at P140 (arrows). **l**, Extravascular fibrinogen signal in control and *SOD1^G93A^* sciatic nerves at progressive stages, normalized to average control value. **m**, **n**, S*aphenous* (sensory, **m**) and *quadriceps* (motor, **n**) nerves from P90 *SOD1^G93A^* mice injected systemically with HRP, detected by tyramide amplification. HRP is confined to vessels in sensory nerves but extravasates in motor nerves and is phagocytosed by macrophages (**n**, arrowheads). DAPI, nuclei. **o**, Extravascular HRP-tyramide signal in control and *SOD1^G93A^*motor nerves (P90), normalized to average control value. **p-q**, Spinal cord vessels from control (**p**) and *SOD1^G93A^*(**q**) mice at P120 stained for MFSD2A, PLVAP and CD31. MFSD2A expression remains unchanged in mutants; PLVAP is not induced. **r**, MFSD2A signal in spinal cord vessels of control and *SOD1^G93A^* mice at P120, normalized to average control value. **s-u’**, Sciatic nerves from control (**s**) and *SOD1^G93A^*at P90 (**t**) and P120 (**u**) stained for Caveolin-1 (CAV1), PLVAP and CD31. Dashed contours outline the endoneurium. Boxed areas in **s**, **t**, **u** are magnified in **s’**, **t’**, **u’**, showing only CAV1 and CD31. Representative dashed lines spanning vessel width in **s’**, **t’**, **u’** used to measure CAV1 signal profile in **v**. **v**, CAV1 signal intensity profiles. In control vessels, CAV1 shows two distinct peaks at the plasma membrane; in *SOD1^G93A^* nerves, a diffuse profile indicates increased cytoplasmic distribution. **w-x’**, Dorsal root ganglion (DRG) vessels from wild-type mice stained for PLVAP, CLDN5 and CD31 (**w**). Higher magnification views show PLVAP and CD31 (**x**), and CAV1 with DAPI (nuclei) (**x’**). Representative dashed line spanning vessel width in **x’** used to measure CAV1 signal profile in **y**. **y**, CAV1 signal intensity profiles in PLVAP^+^ DRG vessels show a diffuse pattern, consistent with cytoplasmic localization. Scale bars: a-b’, 50μm; d, e, 50μm; g-h”, 50μm; j-k”, 50μm; m, n, 100μm; p, q, 100μm; s-u, 100μm; s’-u’, 25μm; w, 100μm; x, x’, 25μm.

**Supplementary Fig. 5:**
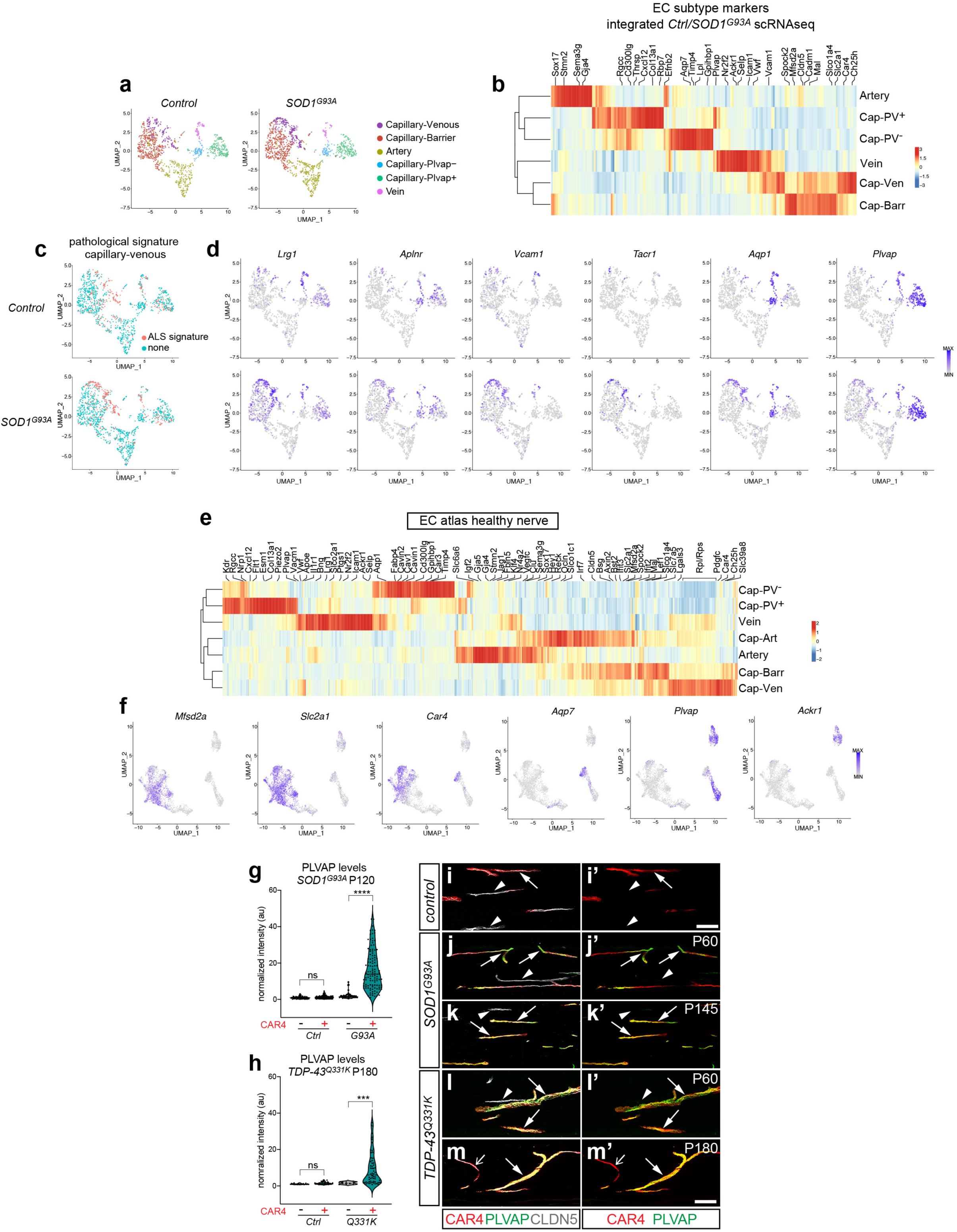
Single-cell profiling of the nerve endothelium in control and *SOD1^G93A^* mice. **a**, UMAP plots of control and *SOD1^G93A^* nerve EC datasets color-coded by subtype. **b**, Expression heatmap (z-score) of EC subtype-marker genes defined in a previous sciatic nerve scRNA-seq dataset{Bhat, 2024 #1}. **c**, Pathological gene signature identified in venous capillaries, visualized on UMAP plots of control and *SOD1^G93A^* nerve ECs using thresholded binary labeling (CellID). **d**, UMAP plots of control and *SOD1^G93A^* nerve ECs showing expression of representative genes from the capillary-venous pathological signature. **e**, Expression heatmap (z-score) of top-100 marker genes for EC subtypes from healthy sciatic nerve EC atlas. **f**, Expression of EC subtype-marker genes visualized in UMAP plots of healthy sciatic nerve EC atlas. **g**, **h**, PLVAP signal in CAR4^+^ and CAR4^−^ endoneurial vessels of sciatic nerves from P120 control and *SOD1^G93A^* mice (**g**) and P180 control and *TDP-43^Q331K^* mice (**h**), normalized to average control value. **x-m’**, Sciatic nerves from control (**i, i’**), *SOD1^G93A^* (**j-k’**), and *TDP-43^Q331K^* (**l-m’**) mice at the indicated stages. Control corresponds to P120. CAR4^+^/CLDN5^+^ venous capillaries (arrows) upregulate PLVAP in mutants. CLDN5^+^/CAR4^−^ vessels remain PLVAP^−^ (arrowheads). **i’-m’** show only CAR4 and PLVAP staining. Scale bars: i-m’, 100μm.

**Supplementary Fig. 6:**
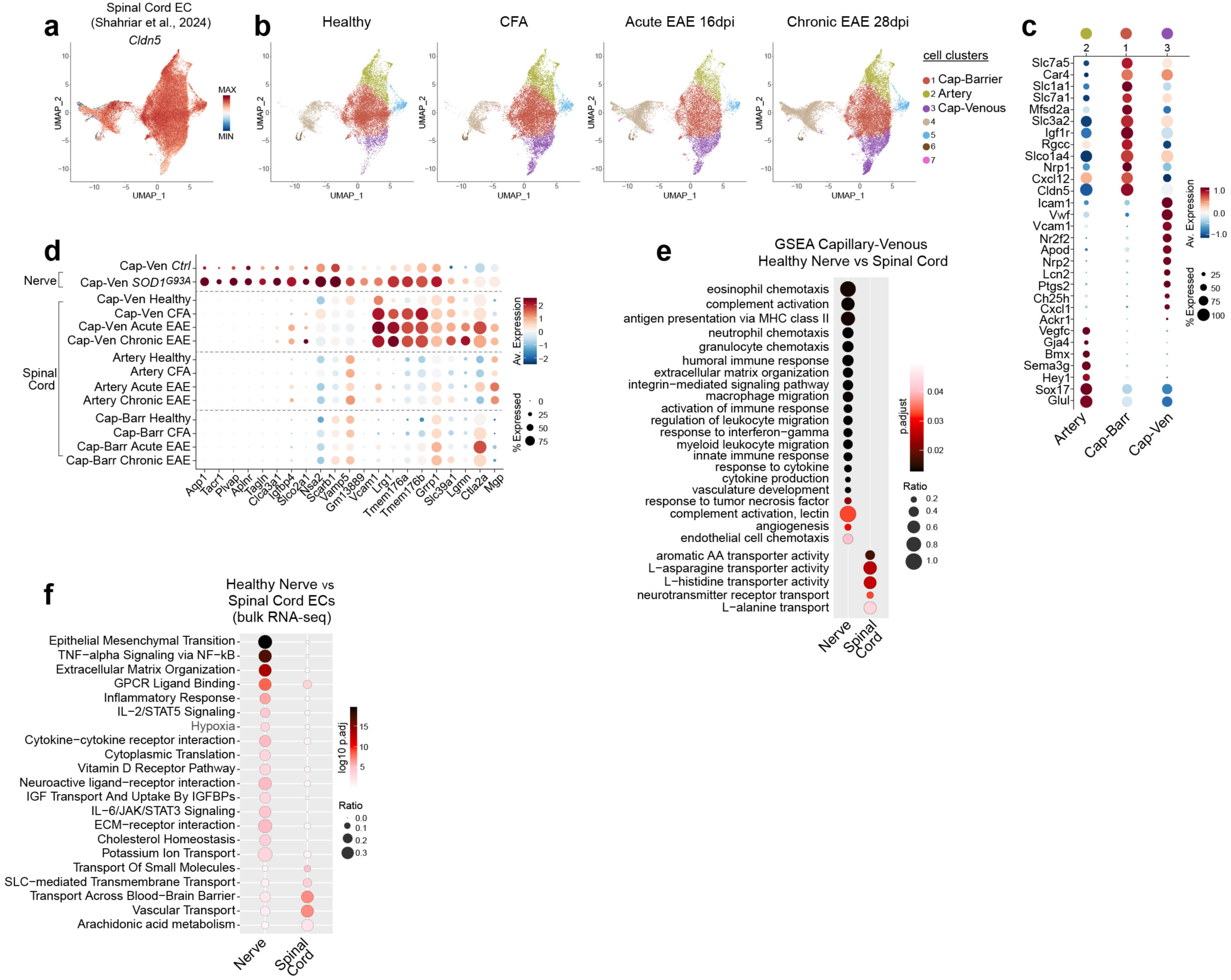
Distinct specialization and reactivity of venous capillaries in spinal cord and nerve. **a**, UMAP plot of integrated spinal cord scRNA-seq EC dataset{Shahriar, 2024 #3} showing expression of *Cldn5*. Only *Cldn5*^+^ cells from the original dataset were retained for the analysis. **b**, Split UMAP plots for healthy, CFA-treated, acute EAE (16-17 days postimmunization, d.p.i.), and early chronic EAE (28-29 d.p.i.) color-coded by unsupervised cell clusters. Clusters 1, 2, and 3 correspond to capillary-barrier, artery, and capillary-venous ECs, respectively. **c**, Dot plot showing expression of marker genes used for EC cluster annotation. **d**, Dot plot showing expression of top DEGs from the nerve capillary-venous pathological signature (padj ≤0.05 and Log2FC ≥0.5 in *SOD1^G93A^*vs. control) in spinal cord EC subtypes under healthy, CFA-treated, acute, and chronic EAE conditions. **e**, GSEA of biological processes in capillary-venous ECs from healthy nerve (from nerve EC atlas) versus spinal cord{Shahriar, 2024 #3}. **f**, GO enrichment of biological processes in ECs from healthy nerve and spinal cord based on bulk RNA-seq (control samples).

**Supplementary Fig. 7:**
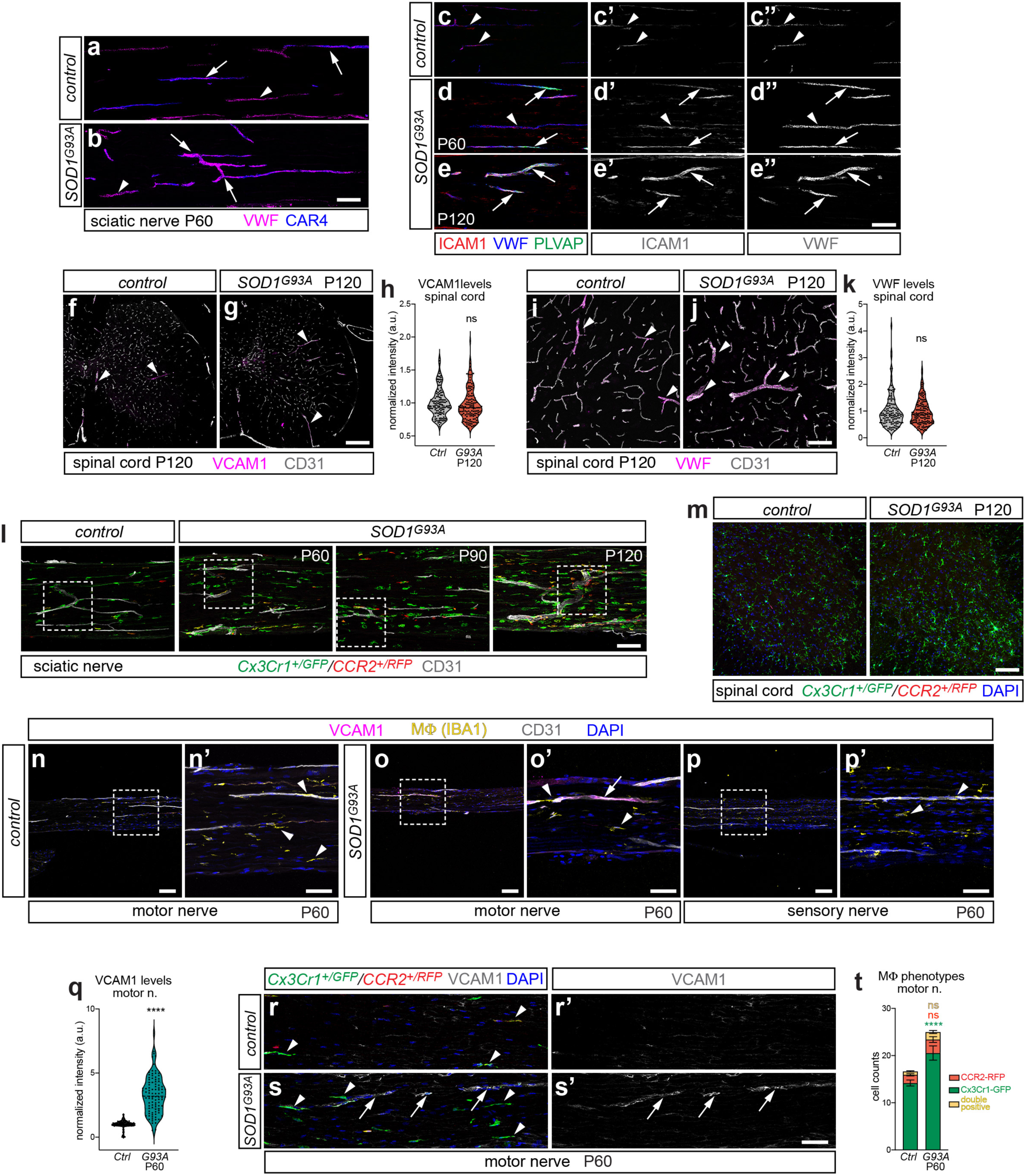
Endothelial activation in ALS nerves, but not spinal cord, coincides with resident macrophage activation. **a**, **b**, Upregulation of VWF in CAR4^+^ capillaries of *SOD1^G93A^* vs. control sciatic nerves at P60 (arrows). Arrowheads point to VWF^+^/CAR4^−^ vessels. **c-e”**, Sciatic nerves from control (**c-c”**) and *SOD1^G93A^*mice at P60 (**d-d”**) and P120 (**e-e”**). ICAM1 and VWF are upregulated in PLVAP+ endoneurial vessels in mutants (arrows). VWF also increases in PLVAP^−^ vessels (arrowheads, **d-d”**). ICAM1 and VWF staining are shown separately in **c’-e’** and **c”-e”**, respectively. **f**, **g**, VCAM1 marks larger veins in both control (**f**) and *SOD1^G93A^* (**g**) spinal cords (arrowheads) and is not upregulated in the microvasculature of mutants at P120. CD31 marks all vessels. **h**, VCAM1 signal in control and *SOD1^G93A^* spinal cord vasculature, normalized to average control value. **i**, **j**, VWF marks larger veins in both control (**i**) and *SOD1^G93A^* (**j**) spinal cord (arrowheads) and is not upregulated in the microvasculature of mutants at P120. CD31 marks all vessels. **k**, VWF signal in control and *SOD1^G93A^* spinal cord vasculature, normalized to average control value. **l**, Sciatic nerves of control and *SOD1^G93A^* mice at the indicated stages, expressing *Cx3cr1^GFP^*/*Ccr2^RFP^* reporter. CD31 marks vessels. Boxed areas are magnified in Fig. 5h-k. **m**, Spinal cord of control and *SOD1^G93A^* mice (P120) expressing *Cx3cr1^GFP^*/*Ccr2^RFP^* reporter. GFP^+^ microglia expand in mutants while RFP^+^ monocytes are absent. **n-p’**, Motor nerves from P60 control (**n**, **n’**) and *SOD1^G93A^*(**o**, **o’**) mice, or *SOD1^G93A^* sensory nerves (**p**, **p’**). Focal induction of VCAM1 in endoneurial vessels is visible in mutant motor nerves (arrowhead in **o’**), but not sensory nerves. IBA1 labels macrophages (arrowheads). Boxed areas in **n**, **o**, **p** are magnified in **n’**, **o’**, **p’**. **q**, VCAM1 signal in endoneurial vessels of P60 control and *SOD1^G93A^* motor nerves, normalized to average control value. **r-s’**, Motor nerves from P60 control and *SOD1^G93A^* mice expressing *Cx3cr1^GFP^*/*Ccr2^RFP^* reporter. VCAM1 marks activated endoneurial vessels in mutants (arrows in **s**, **s’**). Nearby GFP^+^ resident macrophages remain elongated but increase in numbers compared to controls (arrowheads). VCAM1 staining is show separately in **r’**, **s’**. **t**, Number of GFP^+^, RFP^+^, GFP^+^/RFP^+^ double-positive MΦ in control and *SOD1^G93A^* motor nerves (P60). Mean ± SEM. Scale bars: a, b, 100μm; c-e”, 100μm; f, g, 200μm; I, j, 100μm; l, 100μm; m, 100μm; n, o, p, 100μm; n’, o’, p’, 50μm; r-s’, 50μm.

**Supplementary Fig. 8:**
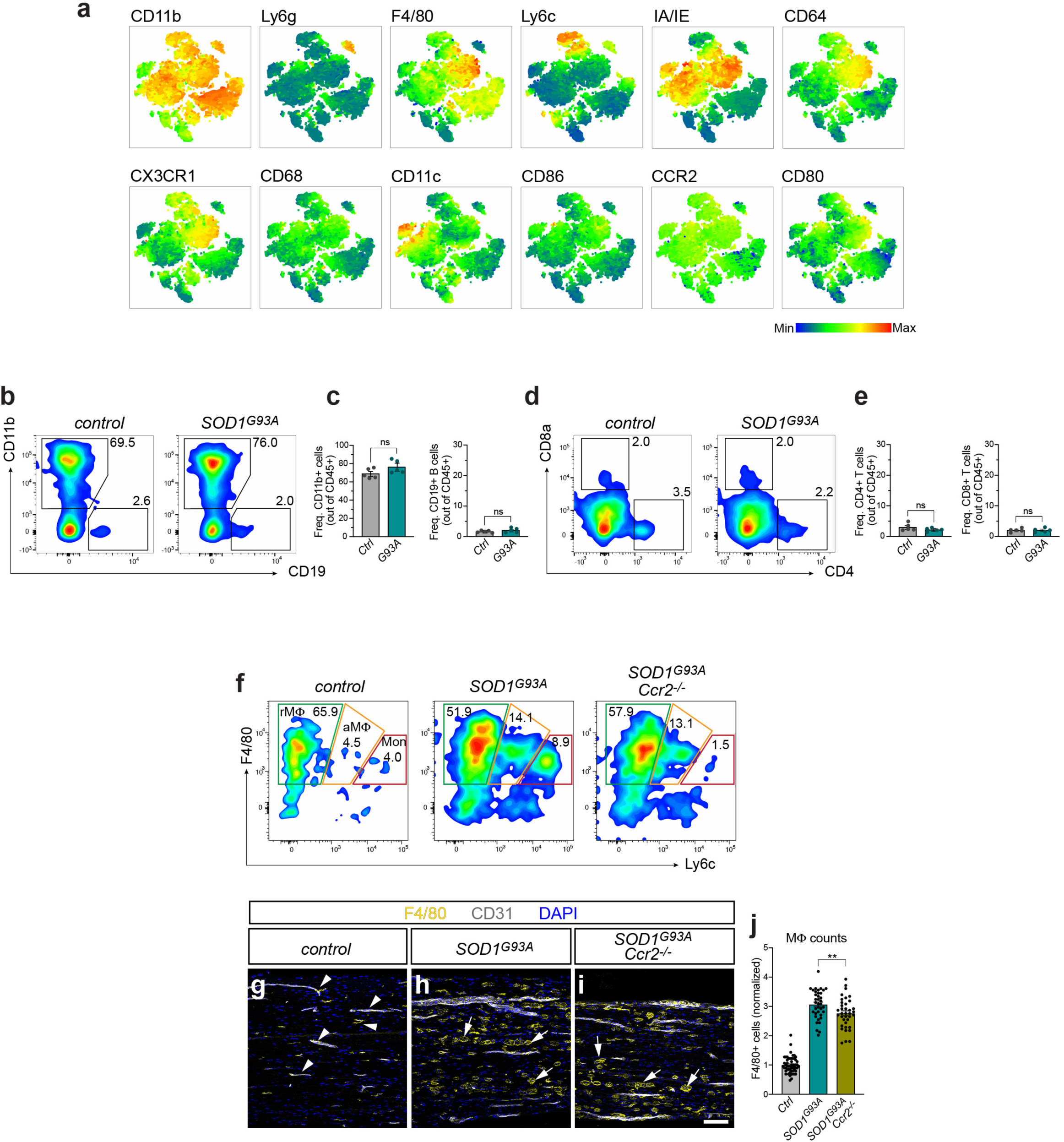
Nerve-resident macrophages drive immune remodeling in *SOD1^G93A^* mice without lymphocyte recruitment. **a**, t-SNE maps of nerve myeloid cells showing antigen expression profiles used to identify populations by unsupervised clustering. **b**, Flow cytometry plots showing near-absent CD19^+^ B cells in sciatic nerves of control and *SOD1^G93A^* mice at P120. **c**, Frequency of CD11b^+^ myeloid cells (left) and CD19^+^ B cells (right) in control and *SOD1^G93A^* nerves. Mean ± SEM. **d**, Flow cytometry plots showing negligible presence of CD4^+^ and CD8^+^ T cells in sciatic nerves of control and *SOD1^G93A^*mice at P120. **e**, Frequency of CD4^+^ (left) and CD8^+^ T cells (right) in control and *SOD1^G93A^* nerves. Mean ± SEM. **f**, Flow cytometry plots showing resident macrophages (rMΦ), activated macrophages (aMΦ), and monocytes (Mon) discriminated based on F4/80 and Ly6c levels in sciatic nerves from control, *SOD1^G93A^*, and *SOD1^G93A^; Ccr2^-/-^* mice at P120. Monocyte infiltration is prevented in *SOD1^G93A^; Ccr2^-/-^* mice. The percentage of aMΦ is not reduced in *SOD1^G93A^; Ccr2^-/-^* compared to *SOD1^G93A^*indicating that they primarily derive from rMΦ. **g-i**, Macrophages identified by F4/80 staining in sciatic nerves from control, *SOD1^G93A^*, and *SOD1^G93A^; Ccr2^-/-^* mice at P120. CD31 marks vessels; DAPI, nuclei. Macrophages are elongated in controls (arrowheads) and become globular/phagocytic in mutants (arrows). **j**, Macrophage counts in *SOD1^G93A^*, and *SOD1^G93A^; Ccr2^-/-^* sciatic nerves based on F4/80 staining. Mean (normalized to control) ± SEM. Scale bar: g-i, 100μm.

**Suppl. Fig. 9:**
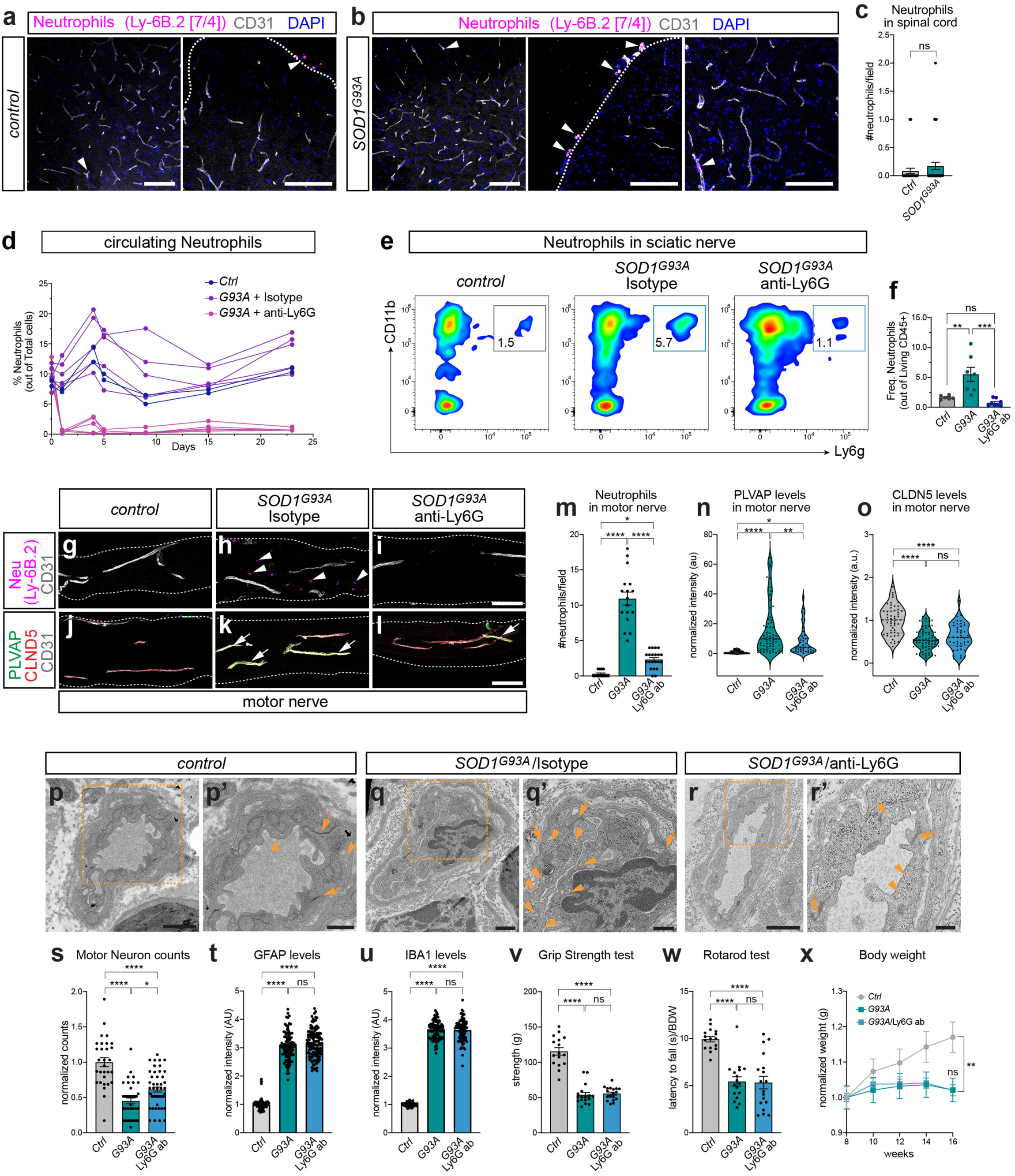
Neutrophil infiltration in *SOD1^G93A^* nerves, but not spinal cord, contributes to vascular damage. **a**, **b**, Neutrophils identified by Ly-6B.2 (7/4) staining in the spinal cord of control (**a**) and *SOD1^G93A^* (**b**) mice at P120. Few neutrophils (arrowheads) are seen within the vasculature or associated with the meninges (a, left; b, middle) in both genotypes. CD31 marks vessels; DAPI, nuclei. Dashed contours outline the spinal cord parenchyma. **c**, Number of neutrophils infiltrating the spinal cord of control and *SOD1^G93A^*mice. Extravascular neutrophils are rare, with no significant difference between genotypes. **d**, Percentage of neutrophils in peripheral blood assessed during anti-Ly6G treatment. Circulating neutrophils are similar between control and *SOD1^G93A^*mice and are depleted by anti-Ly6G treatment. **e**, Flow cytometry plots showing increased neutrophils (CD11b^+^ Ly6G^+^) in the sciatic nerves of *SOD1^G93A^* mice (P120) and their depletion following anti-Ly6G treatment. **f**, Frequency of neutrophils in sciatic nerves of control, *SOD1^G93A^*/isotype-treated and *SOD1^G93A^*/anti-Ly6G-treated mice at P120. **g-i**, Neutrophils [Ly-6B.2 (7/4), arrowheads] detected in *SOD1^G93A^ quadriceps* nerves (**h**), but not in control (**g**) or *SOD1^G93A^*/anti-Ly6G-treated (**i**) mice at P120. CD31, vessels. **j-l**, PLVAP induction in CLDN5^+^ endoneurial vessels of *SOD1^G93A^ quadriceps* nerves at P120 (**k**, arrows) is reduced in *SOD1^G93A^*/anti-Ly6G-treated mice (**l**). PLVAP is absent in control (**j**). CD31, vessels. **m**, Neutrophil counts [Ly-6B.2 (7/4) staining] in motor nerves of control, *SOD1^G93A^*/isotype-treated and *SOD1^G93A^*/anti-Ly6G-treated mice at P120. Mean ± SEM. **n**, **o**, PLVAP (**n**) and CLDN5 (**o**) signal in motor nerve endoneurial vessels at P120, normalized to average control value. **p-r’**, TEM images of endoneurial vessels from control (**p**, **p’**), *SOD1^G93A^*/isotype-treated (**q**, **q’**) and *SOD1^G93A^*/anti-Ly6G-treated (**r**, **r’**) motor nerves. Boxed areas in **p**, **q**, **r** are magnified in **p’**, **q’**, **r’**. Pinocytotic vesicles (arrowheads) are frequent in *SOD1^G93A^* (**q’**) but not in controls (**p’**) and anti-Ly6G-treated mutants (**r’**). Partial opening of EC junctions (arrows) is observed in *SOD1^G93A^* (**q’**) mutants but not in controls (**p’**) and anti-Ly6G-treated mice (**r’**). **s**, Number of ChAT^+^ motor neurons in spinal cord sections of control, *SOD1^G93A^*/isotype-treated and *SOD1^G93A^*/anti-Ly6G-treated mice at P120. Mean (normalized to control) ± SEM. **t**, **u**, Quantification of astrocytes (GFAP^+^) (**t**) and microglia (IBA1^+^) (**u**) in the spinal cord of control, *SOD1^G93A^*/isotype-treated and *SOD1^G93A^*/anti-Ly6G-treated mice at P120. Mean (normalized to control) ± SEM. **v**, All-limb grip strength test in control, *SOD1^G93A^*/isotype-treated and *SOD1^G93A^*/anti-Ly6G-treated mice at P120. Mean ± SEM. **w**, Rotarod motor performance test in control, *SOD1^G93A^*/isotype-treated and *SOD1^G93A^*/anti-Ly6G-treated mice at P120. Mean ± SEM; normalized to body weight, BDW). **x**, Body weight measurement over time in control, *SOD1^G93A^*/isotype-treated and *SOD1^G93A^*/anti-Ly6G-treated mice. Mean (normalized to initial BDW) ± SEM. Anti-Ly6G treatment started at 12 weeks (P90). Scale bars: a-b, 100μm; g-l, 100μm; p, q, 1μm; r, 2μm; p’, q’, r’, 500nm.

**Suppl. Fig. 10:**
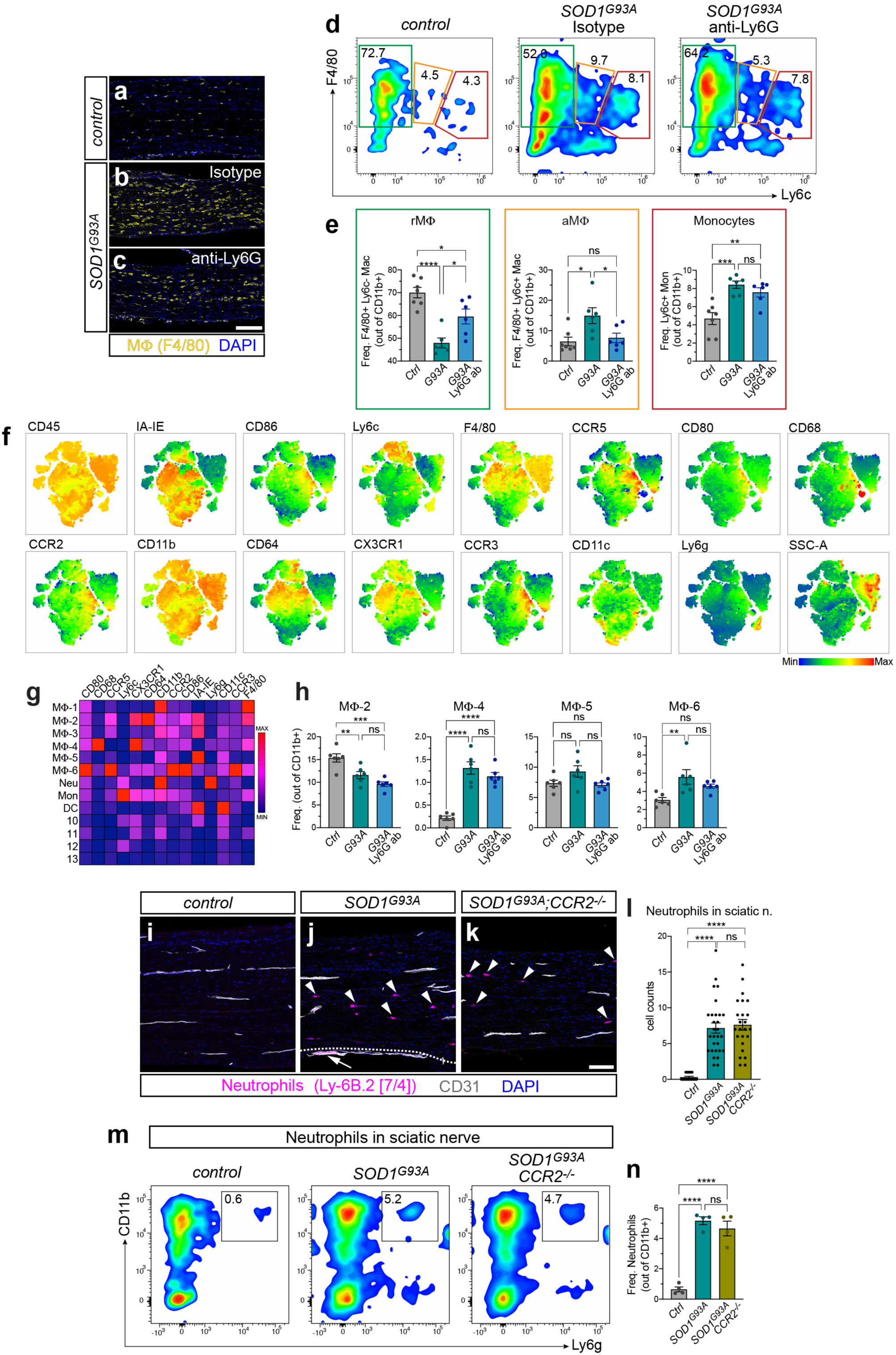
Neutrophils modulate macrophage activation in nerves independently of monocyte recruitment. **a-c**, Macrophages identified by F4/80 staining in sciatic nerves from control (**a**), *SOD1^G93A^*/isotype-treated (**b**) and *SOD1^G93A^*/anti-Ly6G-treated (**c**) mice at P120. DAPI, nuclei. Quantification shown in Fig. 6q. **d**, Flow cytometry plots from sciatic nerves of control, *SOD1^G93A^*/isotype-treated and *SOD1^G93A^*/anti-Ly6G-treated mice at P120 showing resident macrophages (rMΦ, green gate), activated macrophages (aMΦ, orange gate), and monocytes (red gate) discriminated based on F4/80 and Ly6c levels. **e**, Frequency of rMΦ, aMΦ and monocytes (gated from CD11b^+^ cells) from sciatic nerves. Mean ± SEM. **f**, t-SNE maps of nerve myeloid cells showing antigen expression profiles used for unsupervised clustering. **g**, Normalized MFI of surface markers used to define myeloid cell clusters in unsupervised analysis. **h**, Frequency of MΦ2-6 subsets in sciatic nerves identified by unsupervised t-SNE analysis (gated from CD11b^+^ cells). MΦ2 are non-activated (resident) macrophages, reduced in *SOD1^G93A^* mutants. MΦ4 and MΦ6 are activated subsets that increase in mutants. No significant differences are observed between *SOD1^G93A^*/isotype-treated and *SOD1^G93A^*/anti-Ly6G-treated mice. Oher cell clusters are shown in Fig. 6u, v. Mean ± SEM. **i-k**, Neutrophils identified by Ly-6B.2 (7/4) staining in sciatic nerves from control (**i**), *SOD1^G93A^* (**j**), and *SOD1^G93A^; Ccr2^-/-^* (**k**) mice at P120. CD31 marks vessels; DAPI, nuclei. Infiltrating neutrophils (arrowheads) are present in both mutant genotypes (**j**, **k**) but not in controls (**i**). **l**, Neutrophil counts in sciatic nerves based on Ly-6B.2 (7/4) staining. Neutrophils increase in both *SOD1^G93A^* and *SOD1^G93A^; Ccr2^-/-^* mutants, with no significant difference between the two genotypes. Mean (normalized to control) ± SEM. **m**, Flow cytometry plots from the sciatic nerves of control, *SOD1^G93A^*, and *SOD1^G93A^; Ccr2^-/-^* mice at P120 showing a comparable increase of neutrophils (CD11b^+^ Ly6G^+^) in both mutant genotypes. **n**, Frequency of neutrophils in sciatic nerves of control, *SOD1^G93A^*, and *SOD1^G93A^; Ccr2^-/-^*. Mean ± SEM. Scale bars: a-c, 200μm; i-k, 100μm.

**Suppl. Fig. 11:**
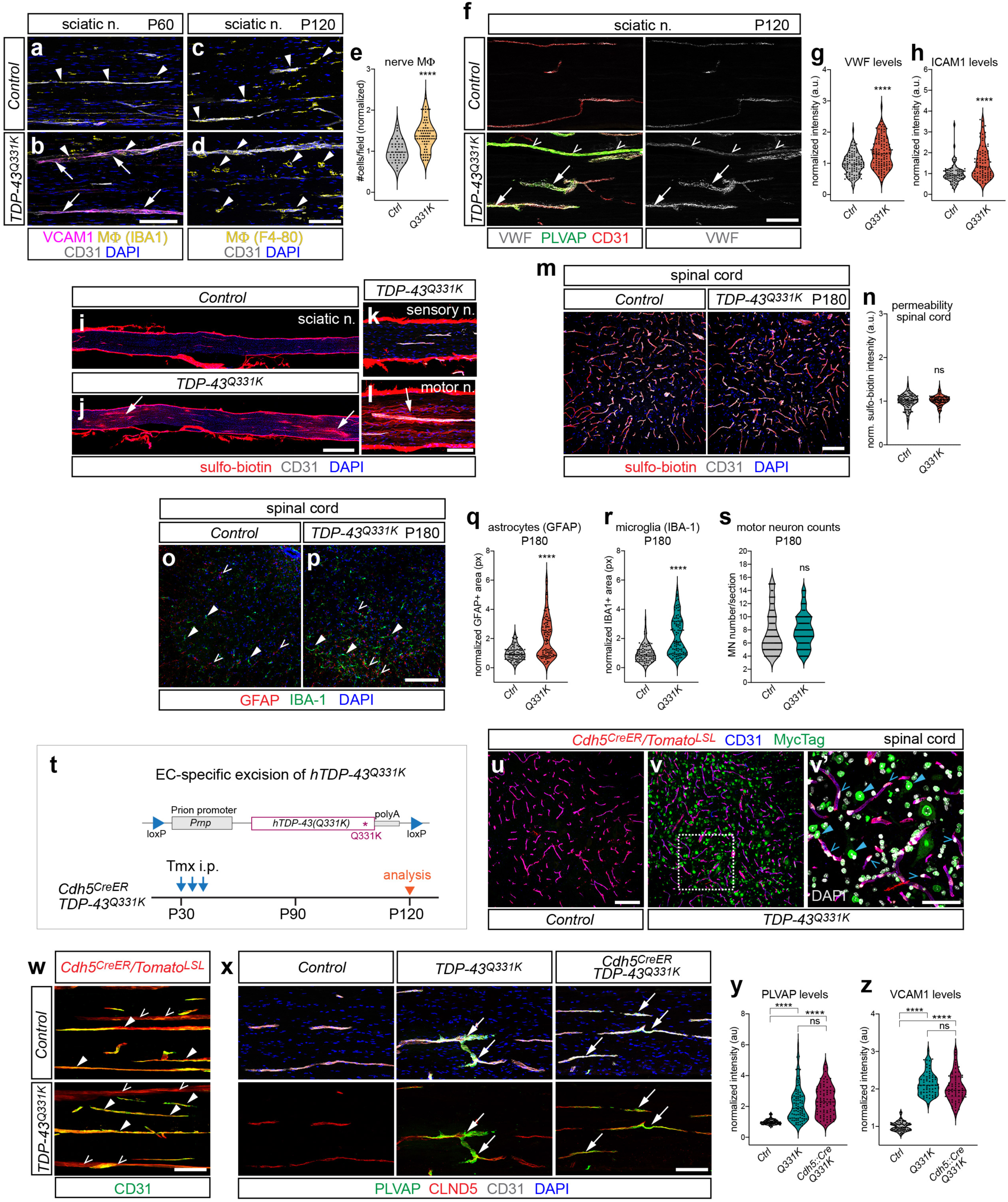
Vascular defects in peripheral nerves but not the spinal cord of *TDP-43^Q331K^* mice. **a**, **b**, Sciatic nerve endoneurial vessels upregulate VCAM1 in P60 *TDP-43^Q331K^* mice (**b**, arrows) compared to controls (**a**). Macrophages (IBA1^+^, arrowheads) are comparable between genotypes at this stage. CD31 marks vessels; DAPI, nuclei. **c**, **d**, Macrophages (F4/80^+^, arrowheads) acquire a globular/phagocytic morphology in P120 *TDP-43^Q331K^* sciatic nerves (**d**) compared to controls (**c**). CD31, vessels; DAPI, nuclei. **e**, Macrophage counts (F4/80^+^) in sciatic nerves of control and *TDP-43^Q331K^*mice at P120, normalized to average control value. **f**, Upregulation of VWF and PLVAP in sciatic nerve endoneurial vessels from P120 *TDP-43^Q331K^*mice (arrows). Some PLVAP vessels maintain low VWF expression (open arrowheads). CD31 marks all vessels. **g**, **h**, VWF (**g**) and ICAM1 (**h**) signal in endoneurial vessels of control and *TDP-43^Q331K^*sciatic nerves at P120, normalized to average control value. **i-l**, Sciatic nerves from control mice (**i**), and sciatic (**j**), *saphenous* (sensory, **k**), and *quadriceps* (motor, **l**) nerves from P180 *TDP-43^Q331K^*mice injected with sulfo-biotin. The tracer diffuses into the sciatic and motor nerve parenchyma in mutants (**j**, **i**, arrows). CD31, vessels; DAPI, nuclei. **m**, Spinal cords from P180 control and *TDP-43^Q331K^* injected with sulfo-biotin. The tracer is retained in the vasculature in both genotypes. CD31, vessels; DAPI, nuclei. **n**, Extravascular sulfo-biotin signal in control and *TDP-43^Q331K^* spinal cords at P180, normalized to average control value. **o**, **p**, Spinal cords from P180 control (**o**) and *TDP-43^Q331K^* (**p**) mice stained for GFAP and IBA1 to reveal reactive astrocytes (open arrowheads) and microglia (closed arrowheads), respectively. **q**, **r**, Quantification of astrocytes (GFAP^+^) (**q**) and microglia (IBA1^+^) (**r**) in P180 control and *TDP-43^Q331K^* spinal cords. Immunoreactivity area normalized to average control value. **s**, Number of ChAT^+^ motor neurons in spinal cord hemisections of P180 control and *TDP^Q331K^* mice. **t**, Schematic of *hTDP-43^Q331K^*transgene driven by the murine prion promoter and flanked by loxP sites. Bottom: Experimental timeline for Cre-dependent excision of *TDP-43^Q331K^* from ECs with *Cdh5^CreER^*. **u-v’**, Spinal cord vessels labeled by lox-STOP-lox (LSL) Tomato reporter driven by *Cdh5^CreER^* in control (**u**) and *TDP-43^Q331K^* mice (**v**, **v’**). Myc-Tag marks *TDP-43^Q331K^*-positive nuclei in mutants (**v’**, closed arrowheads) but is absent in EC nuclei (**v’**, open arrowheads). The boxed area in **v** in magnified in **v’**. CD31, vessels. **w**, *Cdh5^CreER^*-mediated recombination in sciatic nerve ECs (CD31^+^) confirmed by vascular Tomato reporter labeling (closed arrowheads). Tomato also labels the Cdh5^+^ perineurium (CD31^−^, open arrowheads). **x**, PLVAP induction (arrows) in CLDN5^+^ endoneurial sciatic nerve vessels is comparable between *TDP-43^Q331K^* and *Cdh5^CreER^*; *TDP-43^Q331K^* mutants (P120). CD31, marks all vessels; DAPI, nuclei. Lower panels show only PLVAP and CLDN5 staining. **y**, **z**, PLVAP (**y**) and VCAM1 (**z**) signal in endoneurial vessels of control, *TDP-43^Q331K^*, and *Cdh5^CreER^*; *TDP-43^Q331K^* sciatic nerves at P120, normalized to average control value. Scale bars: a-d, 100μm; f, 100μm; i, j, 500μm; k, l, 100μm; m, 100μm; o, p, 100μm; u, v, 100μm; v’, 50μm; w, 100μm; x, 100μm.

